# Pluripotency and Transcriptional Pausing Disrupt Circadian Rhythms to Facilitate the Enrichment of Cancer Stem Cells

**DOI:** 10.64898/2026.07.10.737844

**Authors:** Edward A Gonzalez, Dahui Wang, Maciej Jeziorek, Shahd Mohamed, Lauren S. Sherman, Premananda Indic, Patricia Soteropoulos, Mainul Hoque, Seth Raphael Goldman, Karen Adelman, Lanjing Zhang, Pranela Rameshwar, Jean-Pierre Etchegaray

**Author notes:** These authors contributed equally (Edward A Gonzalez, Dahui Wang, Maciej Jeziorek).

## Abstract

Triple negative breast cancer (TNBC) is the most aggressive breast cancer subtype, enriched for cancer stem cells (CSCs), which are the cause of tumor recurrence. CSCs are responsible for tumor initiation and propagation; however, the molecular mechanisms underlying their formation remain largely unclear. We show that carboplatin treated TNBC cells lose their circadian rhythms and promote the enrichment of CSCs. Notably, genetic ablation of the circadian clock by itself, without carboplatin treatment, facilitated the formation of CSCs along with their ability to generate 3D tumorspheres and mouse tumors enriched with CSCs. Mechanistically, we identified an antagonistic interplay between the circadian clock and pluripotency, whereby the core pluripotent factor OCT4 disrupts the expression of circadian timekeeping genes to disable the circadian clock. Furthermore, we uncovered a transcriptional pausing program, controlling the circadian clock, to be perturbed in carboplatin treated TNBC cells. Concordantly, based on gene expression analysis from The Cancer Genome Atlas (TCGA), we found that the uncoupling of transcriptional pausing and the circadian clock correlated with low survivability across diverse cancer types. Moreover, cancer patients with poor prognosis exhibit low expression of the core timekeeping genes *Clock*, *Npas2, Bmal1* and *Rorc*. Lastly, we restored circadian rhythms in *Oct4* deficient TNBC cells and impaired their ability to generate tumorspheres and decreased the number of CSCs in mouse tumors. Overall, our findings demonstrate an unprecedented mechanism for the formation of CSCs that is dependent on the loss of circadian rhythms and thereby has eminent implications for developing new cancer therapies.

**SIGNIFICANT STATEMENT:** Circadian rhythms are absent in pluripotent stem cells; however, their presence or absence in cancer stem cells has remained undetermined. Here, we implemented a carboplatin-based paradigm to enrich for cancer stem cells. We found that upon carboplatin treatment, triple negative breast cancer cells lost their circadian rhythms. Strikingly, the formation of cancer stem cells is diminished by partial restoration of circadian cycles achieved by knocking down the core pluripotency gene *Oct4*. Mechanistically, we observed an alteration of transcriptional pausing in cancer stem cells that may be implicated in the destruction of the circadian clock.

## INTRODUCTION

Despite significant advances in early diagnosis and therapeutic treatments, cancer remains a clinical challenge. A power force for cancer initiation, progression and relapse relies in the formation of cancer stem cells (CSCs) representing a minor subpopulation of whole cells within a given tumor (<5%) (1, 2). CSCs are self-renewing malignant cells with multipotent capacity and capable of tumor initiation, propagation, metastasis, and relapse. Conventional cancer therapies target the bulk of tumor cells, but are unable to target CSCs, which is currently the most challenging aspect in cancer treatment. The formation of CSCs can be induced upon reprogramming of tumor cells in response to diverse tumor microenvironment and therapeutic pressures, which contributes to tumor heterogeneity and growth(3–8). Earlier findings indicated that gene expression signatures in poorly differentiated aggressive human tumors are reminiscent to expression profiles observed in ESCs (9). This observation is further substantiated by recent analyses describing similar gene regulatory programs between somatic cellular reprogramming into induced pluripotent stem cells (iPSCs) and dedifferentiation of tumor cells into CSCs (10). However, the underlying cellular and molecular mechanisms that trigger such dedifferentiation towards CSCs remain largely unclear. Through mechanisms that are not yet fully understood, circadian rhythms are lost during somatic cell reprogramming into iPSCs (11). A definitive, unified mechanism underlying the loss of circadian rhythms in this process remains under active investigation(12). Consistently, circadian rhythms are absent in embryonic stem cells (ESCs) (11, 13). While the core circadian timekeepers *Clock, Bmal1, Per,* and *Cry* are still expressed in pluripotent stem cells, their ability to maintain a circadian feedback oscillator is disabled. ESCs share striking similarities of physiology and gene expression profiles with CSCs(9). Furthermore, parallels of transcriptional and epigenetic regulatory programs have been suggested between somatic cell reprogramming into iPSCs and dedifferentiation of cancer cells into CSCs(10, 14, 15). Therefore, we reasoned that loss of circadian rhythms could facilitate dedifferentiation or reprogramming of cancer cells into CSCs. Additionally, chemotherapeutic agents including platinum-based drugs such as carboplatin were shown to induce the enrichment of CSCs, possibly through dedifferentiation of cancer cells into CSCs (16–22). Differentiated cancer cells are capable of reprograming themselves and acquire stem cell-like properties under chemotherapeutic pressures leading to resistance and relapse, which highlights their plasticity. We used triple negative breast cancer (TNBC) cells subjected to carboplatin treatment as a model system to study the formation of CSCs and their interconnection with the circadian clock.

Circadian rhythms are cell-autonomous molecular clocks composed of autoregulatory transcription-translation feedback loops that are intrinsic to individual cells and integrated to their specific physiological needs (23). About 43% of all protein coding genes follow a circadian expression profile that is cell type specific (24). The central components of the circadian clock include transcription factors activating the expression of their own repressors within 24-hour periodicity. Generally, the transcriptional machinery of the central timekeeping feedback loop consists of the transcription activators, Circadian Locomotor Output Cycles Kaput (CLOCK) and Brain and Muscles ARNT-like Protein 1 (BMAL1), which as heterodimers bind to regulatory regions on *Period* (*Per*) genes, coding for PER1, PER2, PER3 proteins, and *Cryptochrome* (*Cry*) genes, coding for CRY1, CRY2 proteins. Upon their accumulation in the cytosol, PER-CRY complexes enter the nucleus to repress the activity of CLOCK:BMAL1 heterodimer. A second feedback loop consists of two sets of nuclear orphan receptors functioning as transcriptional activators RAR-related orphan receptors (RORα/β) and repressors Nuclear Receptor Subfamily Group D 1/2 (REV-ERBα/β), and whose expression is activated by the CLOCK-BMAL1 heterodimer. These factors compete for REV-ERB/ROR-response elements (RRE) within regulatory sequences of core timekeeping genes including *Bmal1* and *Cry1* to fine-tune their circadian expression (25, 26). Because the circadian clock is intrinsic to the physiology of individual cells, its disruption can interfere with cellular homeostasis leading to changes in gene expression that are conducive to tumorigenesis. The World Health Organization (WHO) designated the disruption of circadian rhythms as a likely carcinogenic factor, which sparked a global interest to understand the molecular bases connecting the circadian clock with cancer (27–29). However, how disrupted circadian rhythms impact the formation of CSCs, which are conducive to tumor initiation, propagation, metastasis, and relapse, remains a major question. Here, we show that loss of circadian rhythms, via genetic ablation or carboplatin treatment, facilitates the formation of CSCs capable of generating 3D tumorspheres and mouse tumors with enriched CSCs. Mechanistically, we identified an interconnection between the circadian clock, transcriptional pausing and the pluripotency gene network that becomes dysfunctional in breast cancer patients. Moreover, uncoupling the interconnection of the circadian clock with transcriptional pausing correlates with poor survivability in a large population of patients from diverse types of cancers.

## RESULTS

### Carboplatin-treated TNBC cells lost their circadian rhythms, which resulted in the enrichment of CSC-like cells

Recent studies show that chemotherapeutic agents, including the platinum-based drug carboplatin, can promote dedifferentiation of cancer cells towards a CSC-like state (5, 16, 17, 19, 30–32). Thus, we implemented a carboplatin-based paradigm to trigger reprogramming of TNBC cells towards CSCs (**Fig. 1A**). Concordantly, we found that carboplatin treated TNBC cells (MDA-MB-231) show increased levels of the stem cell reporter Oct4-GFP, as determined by fluorescent microscopy (**Fig. 1, B** and **C**), and flow cytometry (**Fig. 1D**). Additionally, carboplatin treatment led to an increase of CSC-associated cell surface markers, including CD133 and CD44^high^/CD24^low^ (**Fig. 1D**). We further validated the enrichment of CSC-like cells by examining the expression of genes associated with cellular dedifferentiation such as epithelial-to-mesenchymal transition (EMT). A time-course qPCR analysis during carboplatin treatment shows a progressive upregulation of the EMT promoting genes *Ck19* (*Krt19*), *Snai1* and *Zeb1* (**Fig. 1E**). Concomitantly, carboplatin treated TNBC cells generated significantly larger tumorspheres compared to untreated controls (**Fig. 1, F** and **G**). Therefore, these results establish carboplatin treatment of TNBC cells as a robust experimental paradigm for the enrichment of CSCs. Using this approach, we found over time a gradual attenuation of circadian rhythms in TNBC cells (MDA-MB-231 and HS578T) treated with carboplatin (**Fig. 1H**, and **Fig. S1**, **A** and **B**). The circadian rhythms in untreated TNBC cells were confirmed by real-time monitoring of bioluminescence from two distinct circadian reporters, *Bmal1*-luc or *Per2*-luc (**Fig. S1**, **A** and **B**). Additionally, qPCR analysis revealed a circadian oscillatory pattern in the expression of the timekeeping genes *Bmal1, Per2, Cry2, Rora*, and *Rev-Erba* (**Fig. S1C**). We confirmed that *Clock* mRNA does not exhibit an oscillatory expression profile, as previously reported(33). Moreover, apart from TNBC cells, carboplatin treatment also promoted the disruption of circadian rhythms in osteosarcoma cells (U2OS) that correlated with the upregulation of canonical cancer stem cell markers including *Abcg2, Cd44, Nanog,* and *Oct4* (**Fig. S1, D** and **E**)(34). Collectively, these data demonstrate a loss of circadian rhythms in carboplatin-treated TNBC cells, which promoted the enrichment of CSC-like cells.

**Figure. 1.**
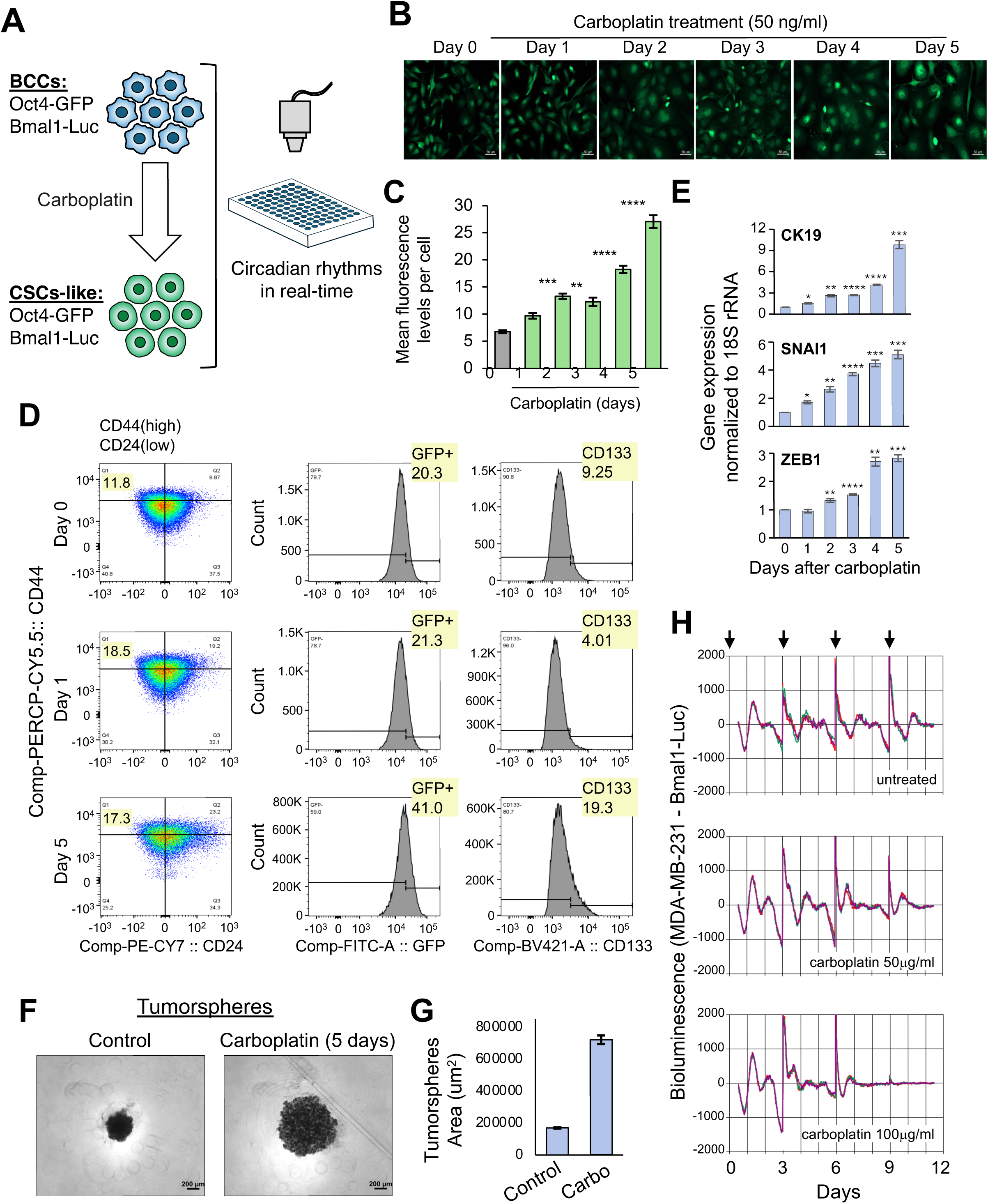
Carboplatin Treatment Disrupts Circadian Rhythms and Enriches MDA-MB-231 cells with Cancer Stem Cell like Population. (**A**) Schematic diagram showing the workflow for monitoring circadian rhythms in MDA-MB-231 treated with carboplatin to enrich for CSCs-like cells. MDA-MB-231 expressing *Bmal1*-*luc* and *Oct4-GFP* reporter are treated with carboplatin and circadian rhythms are monitored in real-time. (**B**)*Oct4*-GFP images from MDA-MB-231 treated with 50 ug/ml of carboplatin treatment over a 5-day period. Images taken with a 20x objective; scale bar is 50 μm. (**C**) Quantification of *Oct4*-GFP mean intensity per cell from Fig. 1B determined by Image J. Error bars represent SEM, (n=6-8 images from 3 independent coverslips, with 30-80 cells per image), pair-end T-test (* = p<0.01, ** = p<0.001, *** = p<0.0001, **** = p<0.00001). **(D**) FACS profiles of carboplatin-treated MDA-MB-231 at days 0, 1, and 5, with pseudocount plots based on CD24 and CD44 signals. Histograms display GFP and CD133 distributions. (**E**) Quantification of RT-qPCR showing the relative gene expression fold changes of *CK19, SNAI1,* and *ZEB1* during the time-course of 50 ug/mL carboplatin treatment of MDA-MB-231, normalized to 18s rRNA. Error bars represent SEM (n=4), pair-end t-test (*p<0.005, **p<5e^-4^, ***p<5e^-5^, ****p<5e^-6^). (**F**) Tumorspheres of untreated and 5-day carboplatin pretreated MDA-MB-231. Images taken with a 5x objective; scale bar is 200 μm. (**G**) Quantification of the surface area for tumorspheres shown in Fig. 1G. Error bars represent SEM (n=), pair-end t-test (**** =p<0.00001). (**H**) Bioluminescent circadian profiles of MDA-MB-231 treated with carboplatin (50ug/ml or 100ug/ml). Arrows indicate resynchronization points every 3 days. Bioluminescent plots show 3 independent circadian profiles from each cell line.

### Carboplatin treatment reprograms the transcriptome of TNBC cells

To further determine the gene networks underlying the loss of circadian rhythms and the enrichment of CSC-like cells, we analyzed the transcriptome of TNBC cells before and after carboplatin treatment. Principal Component Analysis (PCA) of RNA-seq demonstrated distinct clustering within groups, with a strong separation between TNBC and carboplatin-treated TNBC (**Fig. S1F**). Following carboplatin treatment, pathway analysis shows significant downregulation of genes involved in circadian clock networks regulated by CLOCK, BMAL1, and NPAS2, as well as those supporting extracellular matrix (ECM) organization (**Fig. S2A,** and **Supp Table S1**). A disrupted ECM can trigger mechanotransduction signals to induce EMT, which drives cancer cells to undergo dedifferentiation towards CSCs(35). The expression of ECM-related genes is controlled by the circadian clock(36). Consistently, key CSC promoting genes including *Sall4, Klf4, Zeb2,* and *Snai1* were upregulated in response to carboplatin (**Fig. S2B**).

Interestingly, carboplatin downregulated genes from the positive circadian-feedback limb (*Npas2* and *Bmal1*) and upregulated genes from the negative circadian-feedback limb (*Per1, Per2, Cry1,* and *Cry2*) (**Fig. S2**, **C** and **D**). Additionally, the BMAL1 activator, PPARγ, and the circadian synchronizer CREB1, were downregulated upon carboplatin treatment **(Fig. S2D** and **S3A**) (37, 38). This suggests that upregulation of the negative feedback limb triggers an imbalanced clockwork mechanism, which along with an inability to maintain synchronicity, leads to disrupted circadian rhythms in response to carboplatin treatment. In support of this conclusion, either *Bmal1* deficiency or *Per2* overexpression results in the loss of circadian rhythms(39, 40). Moreover, upregulated *Cry2* is linked to poor prognosis in colon cancer(41).

Beyond circadian misexpression, carboplatin triggered the upregulation of genes from the NF-kB signaling pathway (*RelA, RelB, Nfkb2, Chuk, Traf2, Traf4*, and *Traf6*) and the heat-shock response (*Hdf1*, *Hspa1a, Hspa1b, Hspa6, Dnaja1, Dnajb1*, and *Dnajb5*), both of which were shown to directly disrupt circadian rhythms(**Fig. S2, E** - **H,** and **Fig. S3A**)(42–44). Carboplatin upregulated both *RelA* and *RelB* from the canonical and non-canonical NF-kB pathways, respectively (**Fig. S2, E** and **F**). Interestingly, carboplatin treatment caused the upregulation of key genes involved in RNA-polymerase II (Pol II) promoter-proximal pausing including *Nelfa, Hexim1, RN7SK, Mepce, Supt5h*, and *Sirt6* (**Fig. S2**, **I** and **J**)(45, 46). Western blot analysis confirmed upregulation of these pausing factors (NELFB and HEXIM1), as well as the circadian repressors PER2 and CRY2 from the negative feedback limb, along with the CSC drivers b-Catenin and ALDH1A (**Fig. S3**, **B** and **C**)(47, 48).

Overall, the transcriptome of carboplatin-treated TNBC cells shows upregulated genes from the NF-kB pathway, heat-shock response, and b-catenin—all of which resemble the gene profile observed during formation and maintenance of iPSCs(49–51). As mentioned above, these pathways were shown to perturb the circadian clock, which is disrupted during reprogramming into iPSC(12).

### Transcriptional pausing mediates circadian disruption in carboplatin-dependent reprogramming of TNBC cells into a CSC-like state

To further determine the role of transcriptional pausing in this carboplatin-induced CSC paradigm, we performed Precision Run-On sequencing (PRO-seq) to identify Pol II pausing signatures underlying gene expression in carboplatin-treated TNBC cells. Principal Component Analysis (PCA) confirms that samples cluster tightly within each condition (untreated and carboplatin-treated TNBC cells) (**Fig. S3D**). Metagene analysis showed a significant increase of Pol II engaged in productive transcription elongation on upregulated genes in response to carboplatin (**Fig. 2A**). This indicates that upregulated genes in response to carboplatin are due to a Pol II pausing release. Concordantly, the pausing index (PI)—a metric to measure the degree to which Pol II stalls shortly after transcription initiation—is significantly diminished in upregulated genes in carboplatin-treated TNBC cells (**Fig. 2B, Fig. S9A,** and **Supplemental Table S2 and S3, S7, S8**). Additionally, these data show an increased PI on downregulated genes after carboplatin. We identified 542 upregulated genes with low PI that are involved in NF-kB signaling (*Traf6, Tlr4, Gsk3α, RelA, Rel,* and *Klf10*) and heat-shock response (*Hspa1b, Hspa2,* and *Dnaja1*), as well as core genes from the Pol II pausing mechanism itself (*Hexim1* and *Mepce*) (**Fig. 2 C, E** and **F**, and **Fig. S3E**). Among the 621 downregulated genes with high PI are direct regulators of circadian rhythms including *Npas2, Pparγ,* and *Rxrα* (**Fig. 2 D** and **G**). Individually, their deficient expression was shown to disrupt circadian rhythms(52–54). These data support a model whereby changes in transcriptional pausing contribute to circadian disruption in response to carboplatin. Mechanistically, the release of Pol II pausing on genes from the NF-kB pathway and the heat-shock response, along with the increased transcriptional pausing of key circadian regulators, may facilitate the disruption of daily rhythms. Consequently, this Pol II pausing mechanism silencing the clock may promote the reprogramming of carboplatin-treated TNBC cells towards a CSC-like state.

**Figure. 2.**
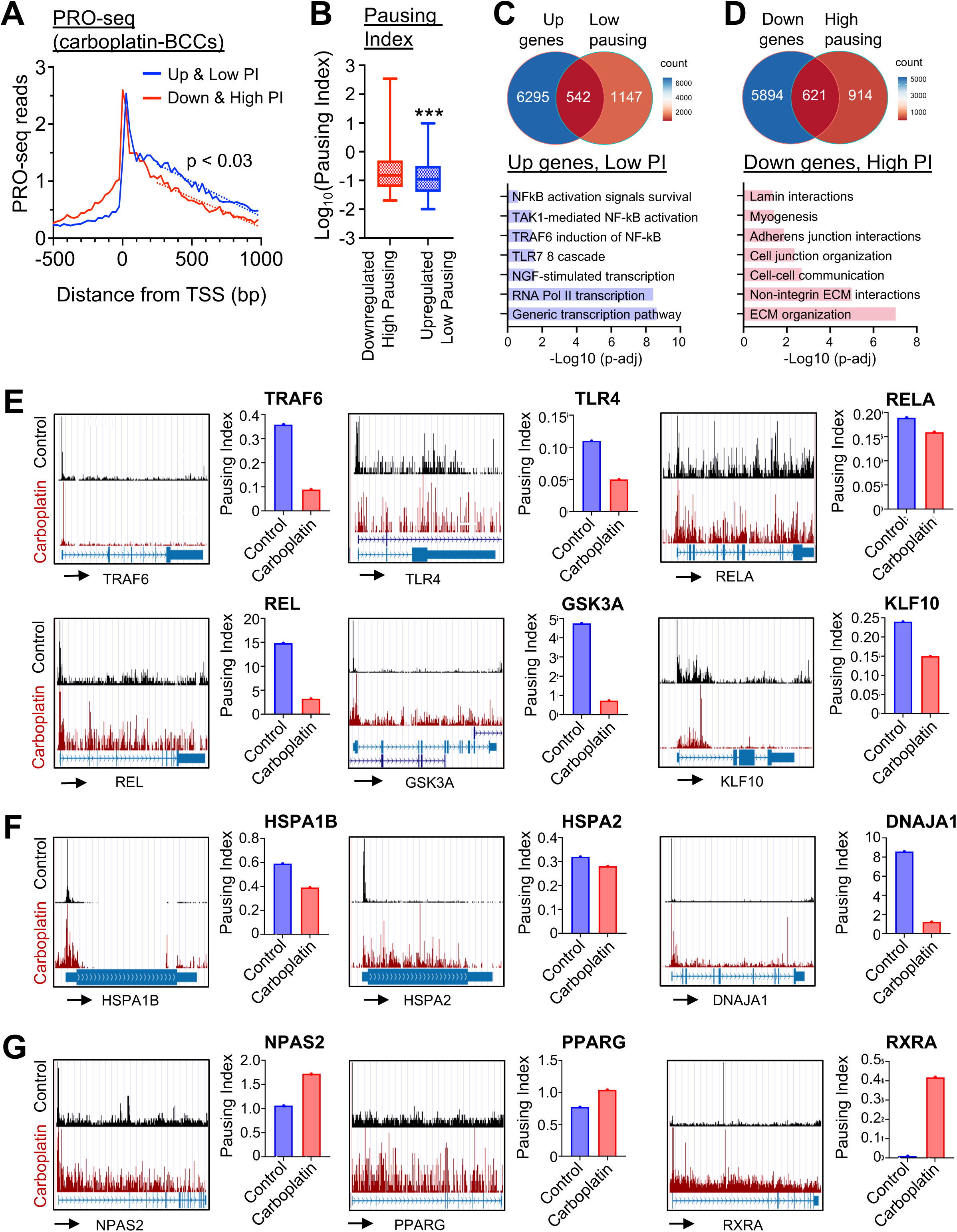
Carboplatin Treatment Results in Gene Expression and Transcriptional Pausing Changes in MDA-MB-231 Cells. (**A**) Aggregate gene profile from PRO-seq between MDA-MB-231 and 5 day carboplatin treated MDA-MB-231 showing upregulated genes with low pausing index and downregulated genes with high pausing index. (**B**) Box and whisker plot of the pausing index of upregulated and downregulated genes in CSCs. Whisker represents min/max value, unpaired T-test were used to calculate p-value (***p<0.0001). (**C**) Venn diagram showing 542 upregulated genes with low pausing in 5 day carboplatin treated MDA-MB-231 and Reactome pathway analysis of these genes. (**D**) Venn diagram showing 621 downregulated genes with high pausing in 5 day carboplatin treated MDA-MB-231 and Reactome pathway analysis of these genes. (**E-G**) UCSC genome browser images from MDA-MB-231 and 5 day carboplatin treated MDA-MB-231 showing genes involved in (**E**) NfkB activation, (**F**) heat-shock response, (**G**) Circadian gene expression.

### OCT4 drives expression of core circadian timekeeping genes in response to carboplatin

The data above indicate that carboplatin induces a transcriptomic reprogramming involving Pol II pausing to suppress or destabilize BMAL1 via NF-kB and heat-shock pathways. However, these data do not fully consolidate the carboplatin-mediated upregulation of genes from the negative circadian feedback limb including *Per* and *Cry*, whose expression is directly activated by BMAL1. The BMAL1-independent mechanisms promoting the expression of *Per* and *Cry* genes include transcription regulators from the heat-shock response (HSF1), hypoxia (HIF1a), glucocorticoid signaling (NR3C1), and cAMP signaling (CREB1). Among these, only heat shock protein HSF1 was upregulated following carboplatin treatment, which could partially explain the increase expression of *Per* genes via HSF1-binding to heat-shock elements (HSEs) in their promoter regions (**Fig. S2D,** and **S3A**)(55). Nevertheless, this mechanism does not explain the upregulation of CRY genes, which lack HSE at their promoters. Therefore, to decipher the mechanisms underlying carboplatin-dependent induction of PER and CRY genes, we implemented a circadian luciferase reporter assay to screen for circadian activators that are independent of BMAL1. We found the core pluripotency factors OCT4 and SOX2 can independently increase the expression of the circadian reporter (*Per2*-luc) in a dose-dependent manner (**Fig. 3A**). However, unlike CLOCK/BMAL1, the activity of OCT4 and SOX2 was not repressed by CRYs, which is needed for maintaining a functional clockwork mechanism. The activity of these pluripotency factors was not affected by CLOCK/BMAL1 (**Fig. 3B**). Additionally, the pluripotency factors KLF4 and NANOG, as well as the reprogramming factor MYC, upregulated the expression of *Per2*-luc in a dose-dependent manner (**Fig. S3F**).

**Figure. 3.**
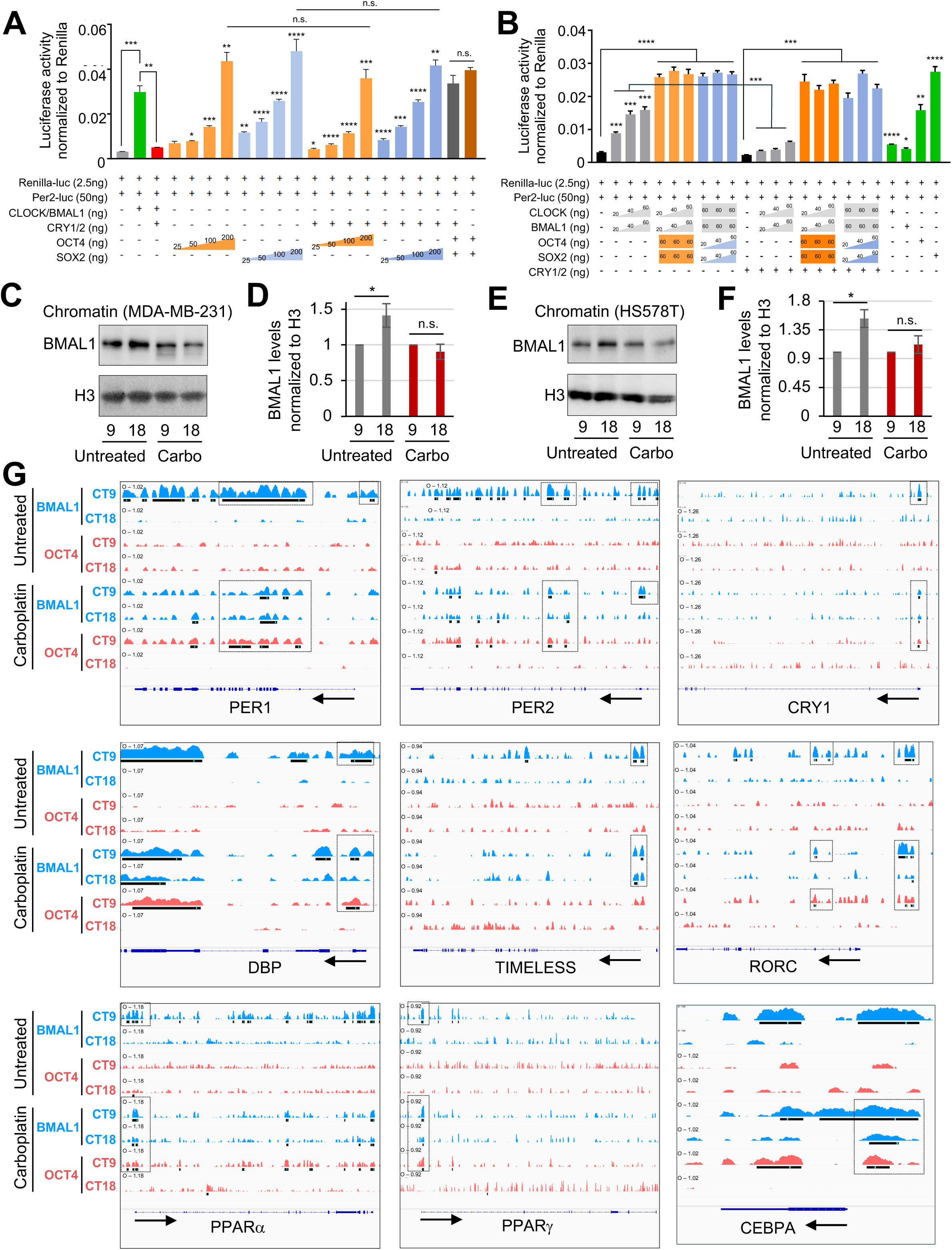
Pluripotency Factors and Carboplatin antagonize Circadian Rhythm on a Genomic Level. (**A-B**) Dual Luciferase assay in HEK293T cells using a Per2-Luc circadian reporter to examine the interaction between pluripotency and circadian genes and Renilla-Luc to normalize transfection efficiency. **(A)** Increasing amounts of OCT4 or SOX2 (25, 50, 100, 200 ng) were transfected with or without circadian repressors Cry1 and Cry2 (10 ng total). **(B)** Reciprocal titration experiments were performed: left, increasing CLOCK and BMAL1 (20, 40, 60 ng each) with constant OCT4 and SOX2 (60ng each), increasing OCT4 and SOX2 (20, 40, 60 ng each) with constant high CLOCK and BMAL1 (60ng each), right: all before mentioned conditions tested with Cry1/Cry2 (10ng total) to assess their repressive influence. Luciferase was normalized to Renilla; n = 4; error bars represent SEM, (*p<0.01, **p<0.001, ***p<0.0001, ****p<0.00001). (**C, E**) Western blot for BMAL1, PER2, and Histone H3 using chromatin extracts from MDA-MB-231 before and after 50ug/ml carboplatin after 5 days synchronized at timepoints 9 and 18 hours. (**D, F**) Quantification from Fig 3C and 3E respectively after normalization to Histone H3. Error bars represent SEM, (n=3-12), Pair-end t-test (*p<0.05, **p<0.01, ***p<0.001). (**G**) IGV browser images for BMAL1 and OCT4 ChIP-seq data from MDA-MB-231 (MDA-MB-231) untreated and treated with carboplatin for 5 days at time points 9hrs and 18hrs on circadian gene *loci*.

To further dissect the interplay between the circadian clock and pluripotency, we performed ChIP-seq analysis at two circadian time-points representing peak (CT9) and trough (CT18) for BMAL1 expression, using specific antibodies targeting BMAL1 and OCT4. Consistent to previous studies, we confirmed the rhythmic abundance of bulk chromatin-bound BMAL1 with a trough at circadian time 9 (CT9) and a peak at CT18, after synchronization with dexamethasone in two different TNBC cell lines, MDA-MB-231 and HS578T (**Fig. 3, C** to **F**)(56). Additionally, these Western blot analyses show that the rhythmic binding of BMAL1 to chromatin was blunted in carboplatin-treated TNBC cells supporting the loss of circadian rhythms in response to carboplatin. The ChIP-seq analysis show significance of peak calling at promoter proximal and intragenic regions of the circadian genes *Per1, Per2, Cry1, Dbp, Timeless, Rorc, Pparα* and *Pparγ* (57), *and Cebpa*(58) (**Fig. 3G**, and **Supplemental Table S4**). Using integrative genomic viewer (IGV), these ChIP-seq profiles show the rhythmic binding of BMAL1 to its circadian target genes with a CT9 peak, as previously shown(59). This rhythmic BMAL1 binding was disrupted after carboplatin treatment showing blunted binding at both CT9 and CT18. Interestingly, except for *Timeless*, we found OCT4 binding to circadian genes with a significant peak calling at CT9 only after carboplatin treatment. For *Dbp* gene, the binding of BMAL1 at promotor proximal sites remains with a rhythmic peak at CT9, but with the addition of OCT4 binding at promoter proximal sites after carboplatin treatment.

Collectively, these ChIP-seq data show the disruption of circadian rhythms in CSC-like cells, generated from carboplatin-treated TNBC cells, by a molecular mechanism that depends on a non-rhythmic BMAL1 binding combined with OCT4 binding at CT9. As shown in **Figure 3, A** and **B**, OCT4 promotes expression of the circadian reporter *Per2*-luc. Constitutive expression of PERs was shown to disrupt circadian rhythms(40). Thus, OCT4 binding to CT9 may be a part of a transcriptional program whereby the pluripotency network disrupts the circadian clockwork mechanism. Intriguingly, while BMAL1 chromatin binding is blunted after carboplatin treatment, we found significantly high levels of chromatin bound PER2 at CT9 (**Fig. S3, G** and **H**). The rhythmic chromatin-bound PER2 with a circadian peak at CT18 was completely disrupted after carboplatin treatment, which may account for the loss of circadian rhythms. Mechanistically, these data suggest that loss of circadian cycles is achieved by an altered recruitment of BMAL1 and OCT4 to circadian target genes upon carboplatin treatment.

### OCT4 deficiency rescues circadian disruption and reduces the enrichment of CSCs

Since OCT4 can drive expression of key circadian timekeeping genes such as *Per* and *Cry* (**Fig. 3**), we evaluated its ability to affect circadian rhythms. We found that TNBC cells stably expressing an shRNA targeting *Oct4* (*shOct4*-cells) exhibited circadian rhythms of greater amplitude compared to TNBC cells stably expressing an shRNA control (shControl, scrambled shRNA sequence) (**Fig. 4A**, and **Fig. S4**, **A** and **B**). Notably, unlike shControl cells, *shOct4*-cells maintained circadian rhythms of 24-hour periodicity under carboplatin treatment (**Fig. 4B**, and **Fig. S4C**). Mathematical modeling to calculate period length under carboplatin treatment showed that shControl-cells displayed no circadian oscillation, while *shOct4*-cells exhibited a nearly 24-hour rhythms (**Fig. S4C**). This result indicates that OCT4 deficiency can rescue the loss of circadian rhythms in TNBC cells treated with carboplatin. Earlier, it was shown that overexpression of wild-type Clock gene can rescue the loss of circadian rhythms in mouse models(60). Concordantly, we observed that overexpression of the *Clock* gene (CLOCKoe) rescued circadian rhythms in carboplatin-treated TNBC cells (**Fig. 4B**).

**Figure. 4.**
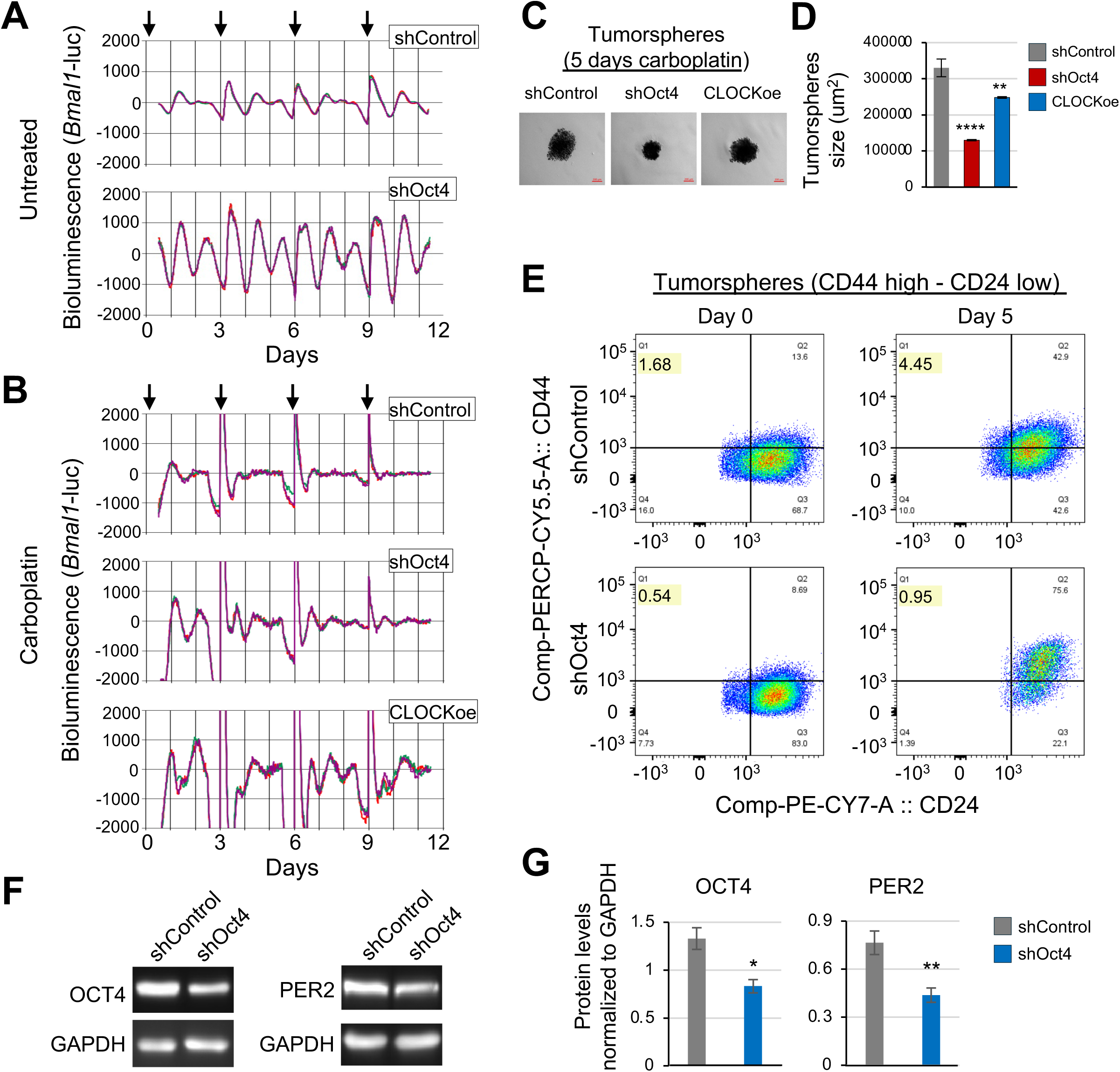
OCT4 Protein Deficiency Strengthens Circadian Rhythms and Decreases Cancer Stem Cell like Population. (**A**) Bioluminescent circadian profiles of MDA-MB-231 stably expressing shRNA targetting *Oct4* in comparison to control. Arrows indicate resynchronization points. Bioluminescent plots show 3 independent circadian profiles from each treatment. (**B**) Bioluminescent circadian profiles of MDA-MB-231 stably knockdown for *Oct4*, or stably overexpressing *CLOCK* treated with 50 ug/ml of carboplatin. Carboplatin treatment was removed on day 6. Arrows indicate resynchronization points. Bioluminescent plots show 3 independent circadian profiles for each condition. (**C**) Representative images of 3D tumorsphere assay using MDA-MB-231 deficient for *Oct4*, or overexpressing *Clock*. 5×10^3^ cells were cultured in 50 ug/ml of carboplatin for 5 days. Phase contrast images 5x objective (Scale bar =200 μm). (**D**) Quantification of the surface area for each tumorsphere shown in Fig. 4C. Error bars represent SEM, pair-end t-test (n=8), (*p< 0.01, ** p<0.00001). (**E**) Flow cytometry for CD44 and CD24 stained MDA-MB-231 knockdown for *Oct4* and control before and after carboplatin treatment. (**F**) Western blot for OCT4, PER2, and GAPDH (loading control) using whole cell extracts from Control and *Oct4* deficient MDA-MB-231. (**G**) Quantification from Fig. 4F after normalization to GAPDH. Error bars represent SEM, (n=3), pair-end T-test (*p<0.05, **p<0.01).

Consistent with the idea of an antagonistic interplay between the circadian clock and the pluripotency network to facilitate the formation of CSCs, we found that under carboplatin treatment both *shOct4*-cells and CLOCKoe-cells generated significantly smaller tumorspheres compared to shControl-cells (**Fig. 4, C** and **D**). Through flow cytometry analysis for the canonical CSC surface markers CD44 and CD24, we observed that tumorspheres generated under carboplatin had a ∼3-fold increase in the number of cells that are CD44^high^/CD24^low^, which is recognized as a *bona fide* CSC signature in TNBC and other cancer types(61–63). Paralleling its circadian restoration under carboplatin, this CSC signature was diminished in shOct4-cells by ∼3-fold before carboplatin and ∼5-fold after carboplatin treatment (**Fig. 4E**). Western blot analysis shows the levels of PER2 were significantly decreased in shOct4-cells (**Fig. 4, F** and **G**). This is consistent with the luciferase reporter assay and ChIP-seq analyses, indicating that OCT4 promotes *Per2* expression (**Fig. 3**). Apart from their significantly smaller size, tumorspheres generated from sh*Oct4*-cells showed a ∼50% reduction in formation efficiency compared to shControl-cells (**Fig. S5, A** and **B**). There were no significant differences of cell cycle progression or cell viability between sh*Oct4*-cells and shControl-cells, as confirmed by flow cytometry and XTT assay (**Fig. S5, C** and **D**). One of the hallmark characteristics of CSCs is their ability to migrate through extracellular matrix (ECM) within the tumor microenvironment to promote tumor growth and invasion of other tissues(64). Thus, to characterize the invasive ability of sh*Oct4*-cells, which form smaller tumorspheres and less efficient than shControl-cells, we implemented a matrigel-dependent invasion assay(65). We found that *shOct4*-cells have significantly lower invasion capacity (∼50% less) compared to shControl-cells (**Fig. S5, E** and **F**).

Collectively, these data support the notion that circadian disruption under carboplatin treatment contributes to the enrichment of CSCs. Notably, partial restoration of circadian rhythms by knocking down Oct4 attenuated CSC enrichment in both untreated and carboplatin treated TNBC cells. Thus, these findings further sustain the idea of an antagonistic interplay between the circadian clock and the pluripotency network whereby OCT4 can disable the circadian clock to promote the formation of CSCs.

### Oct4 deficiency restored the expression of genes involved in circadian rhythms and transcriptional pausing following carboplatin treatment

To further determine how OCT4 contributes to circadian disruption and transcriptomic reprogramming, we performed RNA-seq on shOct4-cells following carboplatin treatment. Post carboplatin treatment, *Oct4* deficient cells expressed significantly higher levels of timekeeping genes from the positive circadian feedback limb including *Bmal1, Clock,* and *Npas2,* along with *Ogt* encoding for O-GlcNac transferase shown to stabilize CLOCK and BMAL1 (**Fig. S6 A, C, D, and Supp Table S5**)(66). Conversely, genes supporting the negative circadian feedback limb such as *Cry2* and *Sirt6*(67) were downregulated in *shOct4*-cells (**Fig. S6D**). Additionally, the downregulation of ECM-related genes we observed in carboplatin-treated TNBC cells (**Fig. S2A**) was restored in *shOct4*-cells (**Fig. S6A**). As mentioned above, downregulation of ECM genes can facilitate EMT(35). Moreover, the expression of CSC-promoting genes involved in dedifferentiation and EMT including *Aldh1a3, Ck19* (*Krt19*), *Snai1* and *Snai3* were significantly downregulated in *shOct4*-cells treated with carboplatin (**Fig. S6B**). Notably, the upregulated expression of genes encoding for transcriptional pausing factors (*Hexim1, Mepce*, and *Nelfa*) that we observed in carboplatin treated TNBC cells (**Fig. S2, I** and **J**), was restored in carboplatin-treated *shOct4*-cells (**Fig. S6 E** and **F**). Furthermore, the expression of genes encoding for the Pol II pausing releasing factors CCNT1 and CCNT2 was upregulated in carboplatin-treated *shOct4*-cells. Interestingly, carboplatin-treated *shOct4*-cells have upregulated levels of genes associated with the prevention of EMT in TNBC including *Cdh1, Tjp1* (*ZO-1*), and *Lama2* (**Fig. S6 G** and **H**)(68–70).

We evaluated the tumorigenic potential of *shOct4*-cells by orthotopic injections in the mammary fat pad of immunodeficient mice. We observed a trend towards a smaller tumor size generated from *shOct4*-cells compared to shControl-cells (**Fig. S8, A** to **D**). Importantly, we found a significant downregulation of the CSC markers CD44, KRT7 and KRT17 (**Fig. S8, E** and **F**).

Overall, these data indicate that pluripotency factors such as OCT4 play a central role in coordinating the transcriptional reprogramming of TNBC cells during carboplatin treatment. OCT4 deficiency restores the expression of key circadian regulators and transcriptional pausing factors, as well as genes involved in maintaining epithelial cell identity while suppressing the expression of CSC promoting genes. These findings further support a model in which OCT4-driven transcriptional reprogramming contributes to circadian disruption and the formation of CSCs.

### BMAL1 deficiency or OCT4 overexpression promote the enrichment of CSCs

To further characterize the incompatibility between circadian rhythms and CSCs, we generated TNBC cells stably expressing an shRNA targeting *Bmal1* (*shBmal1*), as well as TNBC cells stably expressing an *Oct4* overexpression construct (OCT4oe) (**Fig. 5A**). Both *shBmal1*-cells and OCT4oe-cells exhibit disrupted circadian profiles with dampened circadian amplitude (**Fig. 5, B** and **C**). Concordantly, both cell lines could generate significantly larger tumorspheres compared to control cells (**Fig. 5, D** and **E**). Apart from the larger size, *shBmal1*-cells generated tumorspheres 33% more efficiently than shControl-cells (**Fig. S5, A** and **B**). Additionally, the cell cycle progression and viability of *shBmal1*-cells did not differ from shControl-cells (**Fig. S5, C** and **D**). By flow cytometry, we confirmed that *shBmal1*-cells have an over 8-fold higher number of CD44^high^/CD24^low^ CSCs compared to shControl-cells, and generated tumorspheres with a ∼5-fold increase in CD44^high^/CD24^low^ CSCs compared to shControl-cells (**Fig. 5F**). As mentioned above, a main functional feature of CSCs is their ability to migrate through ECM to promote tumor growth and invasion(64). In accordance with their higher increased number of CD44^high^/CD24^low^ CSCs, *shBmal1*-cells showed a ∼4 fold more invasion capacity compared to shControl-cells (**Fig. S5, E** and **F**). Western blot analysis showed that *shBmal1*- and OCT4oe-cells have upregulated PER and CRY proteins (**Fig. 5, G** to **J**). Furthermore, apart from a significant downregulation of CLOCK protein, *shBmal1*-cells have upregulated levels of KLF4 protein (**Fig. 5G**, and **S5, G** and **H**). Overall, these data support our premise that disruption of circadian rhythms, by genetic depletion of Bmal1 or carboplatin treatment, leads to the enrichment of CSC-like cells.

**Figure. 5.**
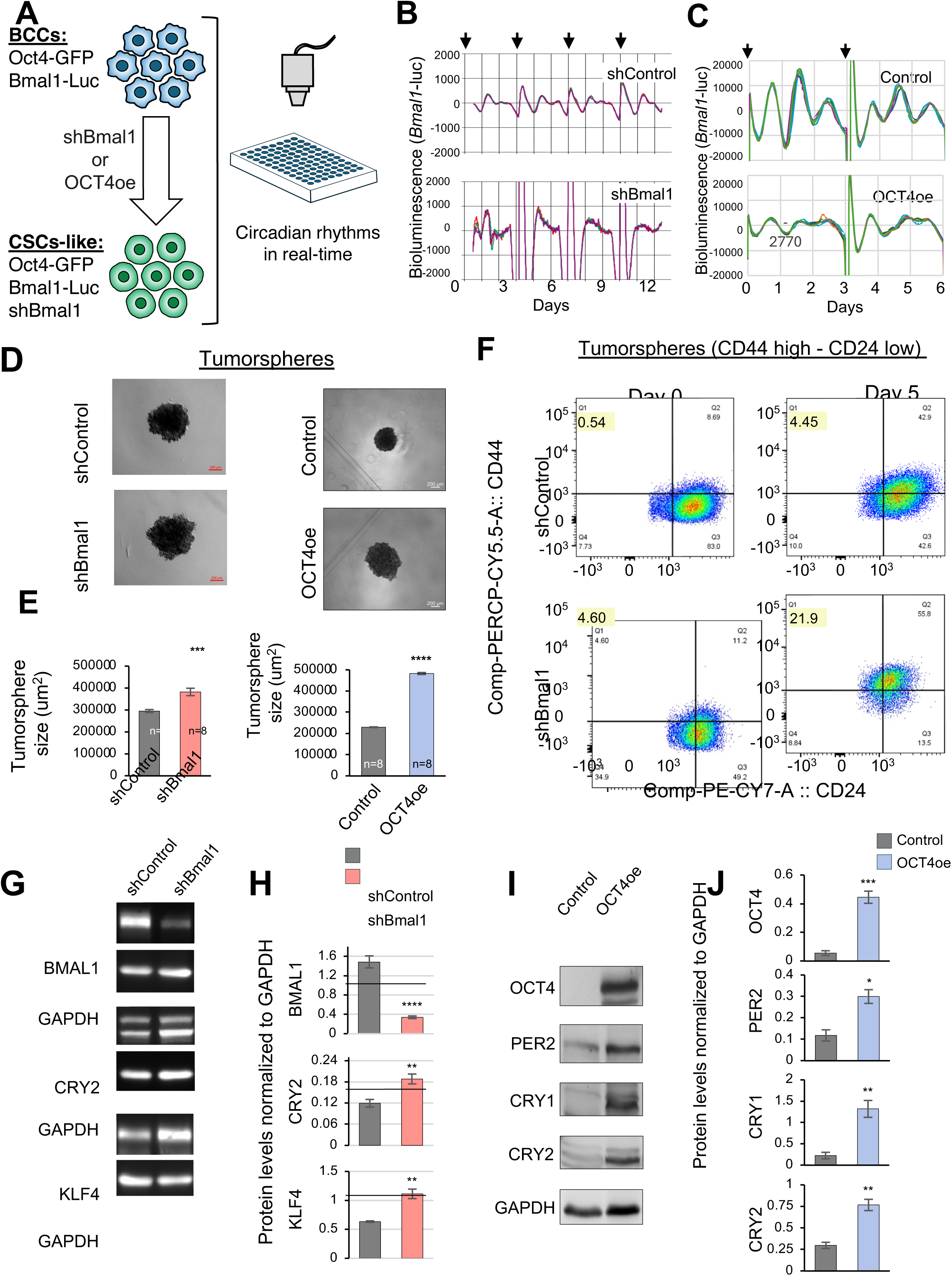
Bmal1 Deficiency and OCT4 Overexpression promotes Cancer Stem Cell like Generation. (**A**) Schematic diagram showing the workflow for monitoring circadian rhythms in MDA-MB-231 deficient in circadian rhythms to enrich for CSCs-like cells. MDA-MB-231 expressing *Bmal1*-*luc* and *Oct4-GFP* reporter are transduced with a lentiviral shRNA to knockdown *Bmal1* or OCT4 overexpression and circadian rhythms are monitored in real-time. (**B**) Bioluminescent circadian profiles of MDA-MB-231 stably expressing shRNA knockdown for *Bmal1* in comparison to control. Arrows indicate resynchronization points. Bioluminescent plots show 3 independent circadian profiles from each treatment. (**C**) Bioluminescent circadian profiles of MDA-MB-231 stably overexpressing OCT4 in comparison to control. Arrows indicate resynchronization points. Bioluminescent plots show 6 independent circadian profiles from each treatment. (**D**) Tumorsphere formation assay using *Bmal1* knockdown or OCT4 overexpression. Phase contrast images taken with 5x objective. (Scale bar=200Dm). (**E**) Quantification of the surface area for tumorspheres shown in Fig. 5D. Error bars represent SEM, (n=8), Pair-end t-test (*** p<0.001, **** p<0.0001). (**F**) Flow cytometry for CD44 and CD24 stained MDA-MB-231 knockdown for *Bmal1* and control between day 0 and day 5. (**G**) Western blot for BMAL1, CRY2, KLF4 and GAPDH (loading control) using whole cell extracts from Control and *Bmal1* deficient MDA-MB-231. (**H**) Quantification from Fig. 4G after normalization to GAPDH. Error bars represent SEM, (n=3), pair-end T-test (**p<0.01, ****p<0.0001). (**I**) Western blot for OCT4, PER2, CRY1, CRY2 and GAPDH (loading control) using whole cell extracts from MDA-MB-231 overexpressing OCT4. (**J**) Quantification from Fig. 5I after normalization to GAPDH. Error bars represent SEM, (n=3-4), pair-end T-test (*p<0.05, **p<0.005, ***p,0<0005).

### Circadian disruption activates the heat-shock response and NF-κB signaling pathway

To investigate the mechanisms underlying how circadian disruption induces the enrichment of CSCs, we performed RNA-seq on *shBmal1*-cells. Pathway analysis revealed a strikingly similar transcriptome to the one observed in carboplatin-treated TNBC cells. *shBmal1* deficiency results in the upregulation of the NF-kB signaling pathway (*Tnfsf14, Tnfsf8, Tnfsf15, Cd70, Nfkb2,* and *RelB*) and the heat-shock response (*Hspa1b, Hspa1a, Hspa, Hspa6, Hspb8, Dnaja1, Dnajb1*, and *Dnajb5*), as well as CSC-promoting genes including *b-catenin* (*Ctnnb1*), *Klf4, Cd44, Ck17* (*Krt17*), and *Ck7* (*Krt7*) (**Fig. S7 A-G, and Supp Table S6**). Interestingly, the genes from the negative circadian feedback limb including *Per1, Per2* and *Cry2* are upregulated in *shBmal1*-cells (**Fig. S7C**). Although, this may seem paradoxical since these genes are transcriptionally activated by BMAL1, we show their expression is upregulated by pluripotency factors such as KLF4, SOX2 and OCT4 (**Fig. 3**, and **Fig. S3F**). Alternatively, *Per* genes contain D-box elements in their promoters, which serve as binding sites for the transcriptional activator DBP(71), which is upregulated in *shBmal1*-cells (**Fig. S7C**).

We confirmed the tumorigenic potential of *shBmal1*-cells via orthotopic injections in the mammary fat pad of immunodeficient mice. The tumor size did not significantly differ upon injection of either *shBmal1*-cells or shControl-cells (**Fig. S8, A** to **D**). However, we found upregulated levels of the CSC markers CD44, KRT7 and KRT17 (**Fig. S8, E** and **F**). Taken together, these data indicate that disruption of circadian rhythms, via *shBmal1* or carboplatin, is sufficient to trigger a transcriptomic reprogramming that is conducive to the formation of CSCs.

### Tumors from breast cancer patients exhibit blunted circadian rhythms

Our data show that loss of circadian rhythms facilitates the formation of CSCs, yet the importance of the circadian clock in normal human breast tissues and how it is disengaged in breast cancer remain limited. Thus, to determine the importance of the circadian clock in cancer patients, we analyzed publicly available RNA-seq transcriptome from The Cancer Genome Atlas (TCGA) database to survey circadian gene expression patterns in breast cancer tumors compared to surrounding normal breast tissues. We used Pearson’s correlations analysis to investigate the correlations between genes involved in circadian rhythms, transcriptional pausing, NF-κB signaling, and pluripotency networks between normal tissue and breast cancer tumors (42) (**Fig. 6 A-E**, and **Fig. S9 B-G**). Notably, these correlations were significantly dampened in tumor tissues from the same patients (**Fig. 6, A** and **B**, and **Fig. S9, B** and **C**). For a better understanding of these bidirectional correlations, we analyzed the relationship among the circadian clock itself and its interplay with pluripotency or transcriptional pausing. We found the self-regulation of core circadian time keeping genes expressed in normal tissues and was significantly bunted in the primary tumor (**Fig. 6C**). Furthermore, the bidirectional correlations between circadian rhythms and the core pluripotency gene network (**Fig. 6D**), as well as circadian rhythms and transcriptional pausing (**Fig 6E**) were lost in primary tumor samples (**Fig. 6D-E**). Interestingly, in normal tissue the circadian and transcriptional pausing pathways show a distinct bidirectional correlation pattern, as indicated by strong positive co-clustering - (area in the black square) (**Fig. 6E**). However, in the primary tumor tissue, this inter-module coupling is lost – while correlations within each pathway (circadian clock or transcriptional pausing) are still present, albeit more diffuse (**Fig. 6E**). Overall, these TCGA data analyses further support the interplay between the circadian clock, pluripotency, and transcriptional pausing in normal breast tissues, which becomes disrupted or uncoupled in primary breast tumors.

**Figure. 6.**
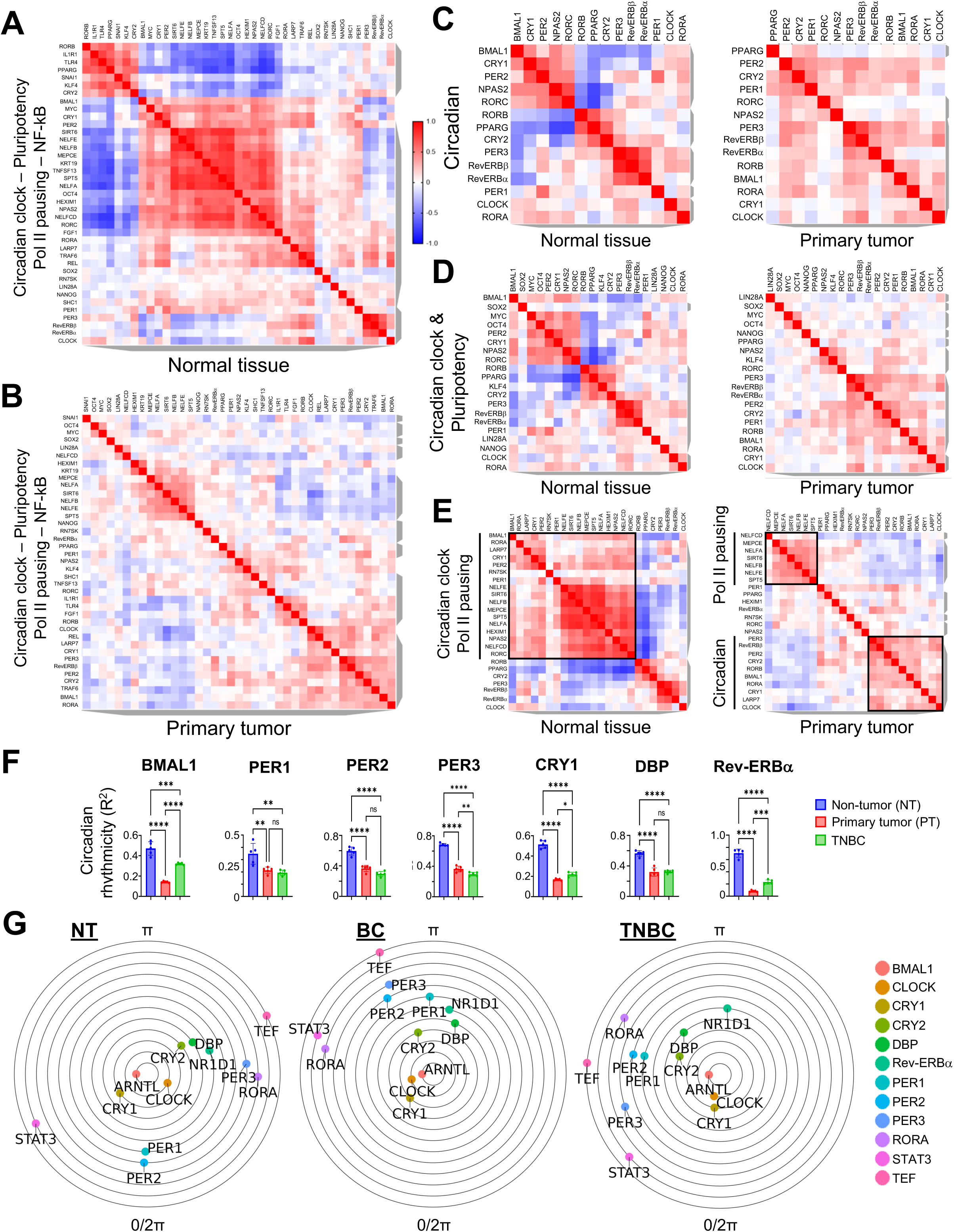
Analysis of TCGA Datasets Reveal Loss of Gene Expression Correlations Between Circadian Clock, Pluripotency, and Transcriptional Pausing Pathways, Along with Disruption of the Inverse Correlations Within the Circadian Regulatory Network. **(A-B)** Spearman correlation heatmap showing bidirectional correlations between genes involved in circadian rhythms, pluripotency, transcriptional pausing, and NF-kB pathways in normal tissues **(A)** and primary tumors **(B)** from breast cancer patients in TCGA dataset. (**C-E**) Spearman correlation heatmaps showing bidirectional correlation between gene in circadian clock pathway **(C)**, between core circadian clock genes and core pluripotency genes **(D)**, and between core circadian clock genes and core transcriptional pausing genes, **(E)** in normal tissues versus primary tumors from breast cancer patients in TCGA. For all heatmaps in this Figure, the color of each cell corresponds to the Z-score value. (**F**) CIRCust based analysis of circadian rhythmicity on core circadian gene between normal tissue and primary tumor and TNBC. (**G**) CIRCust peak phase estimates of circadian genes in Normal tissue, Primary tumor, and TNBC analyzed by CIRCust.

To further evaluate our TCGA analysis in the context of circadian rhythmicity, we applied CIRCular-robUST (CIRCUST), a recently developed method for analyzing rhythms of gene expression using data sets lacking multiple circadian timepoints(72). Using this method, we confirmed circadian rhythmicity of core circadian timekeeping genes in normal tissues (**Fig. 6, F** and **G**, and **Fig. S9H,** and **Fig. S10A,** and **supplemental Table S7**), which were diminished in both primary tumors from multiple breast cancer patients as well as primary tumors from TNBC patients (**Fig. 6, F** and **G**, and **Fig. S10, B** and **C,** and **supplemental Table S7**). Overall, our CIRCUST analyses indicate that circadian gene rhythmicity is normal in healthy tissues but disrupted in primary tumor tissues from breast cancer patients.

### Impaired relationship between the circadian clock and transcriptional pausing correlates with low survival of cancer patients

To further determine the impact of the core circadian timekeeping genes on cancer patient outcomes, we analyzed ∼1000 breast cancer patients within TCGA. We found that patients with low *Clock* expression exhibit a significantly low overall survival outcome (**Fig. S10D**). To assess if this poor survival correlation is conserved in multiple types of cancers, we analyzed a larger population of cancer patients (∼10,000 patients) from a combined TCGA TARGET GTEx data base encompassing RNA-seq data from multiple cancer types including: adrenal gland, bile duct, bladder, brain, breast, cervix, colon, endometrium, esophagus, eye, head and neck, kidney, lining of body cavities, liver, lung, lymphatic tissue, ovary, pancreas, paraganglia, prostate, rectum, skin, soft tissue bone, stomach, testis, thymus, thyroid gland, and uterus. We found that correlation between low *Clock* expression and poor patient outcome gained significance across multiple cancer types (**Fig. S10E**). Apart from low levels of *Clock* expression, the poor outcome from this larger population of patients across multiple cancer types, was extended to lower expression of *Npas2* and *Rorc* (**Fig. S10, F** and **G**). The correlation between low *Bmal1* expression and poor patient outcome was also observed in multiple cancer types including breast, liver, eye, thymus, thyroid and uterus (**Fig. S11A**).

To further clarify whether the interrelationship among circadian genes based on their coordinated expression contribute to patient outcome, we use PCA-based feature selection method on TCGA dataset. We separated patients into “high correlation” and “low correlation” groups based on their contribution towards major axes of gene expression pattern in PCA analysis. We found that a disrupted relationship between transcriptional pausing genes and the circadian genes *Clock*, *Bmal1* and *Npas2* correlated with low survivability in the larger population of cancer patients described above (**Fig. S11B**). Furthermore, TCGA data sets from cancer patients treated with platin-based chemotherapeutics also show poor survivability under low *Clock* or *Rorc* expression (**Fig. S11, C** and **D**).

These analyses indicate that loss of essential circadian clock components along with a disrupted relationship between transcriptional pausing and the circadian clock across multiple human cancers may be a universal molecular signature affecting patient survival (**Fig. S11E**). Mechanistically, under carboplatin treatment, the core pluripotency factor OCT4 is recruited to constitutively activate the expression of timekeeping genes and disrupt circadian rhythms (**Fig. S11F**). Additionally, genes encoding for BMAL1 repressors such as NF-kB, as well as genes promoting CSC-like properties (EMT and heat-shock response)(73) exhibit less Pol II pausing under carboplatin treatment. Meanwhile genes supporting the positive circadian feedback limb (*Npas2, Pparγ* and *Rxrα*) exhibit increased Pol II pausing in response to carboplatin(54, 57). Notably, the circadian clockwork, the core pluripotency network, and transcriptional pausing are part of an integrated molecular module (**Fig. S12**).

### Exosomes from mesenchymal stem cells disrupt circadian rhythms in TNBC cells

It has been recently shown that exosomes released from human MSCs in the bone marrow instruct TNBC cells to dedifferentiate towards CSCs (74). We purified MSC-derived exosomes and characterized them by Nanosight, which uses nanoparticle tracking analysis to determine the size and concentration of exosomes. We used exosomes with an average size of 100nm in diameter (**Fig. S13, A** to **C**). We used flow cytometry to confirm the enrichment of CSCs from TNBC cells (MDA-MB-231 stably expressing the stem cell reporter *Oct4*-GFP along with shControl or sh*Oct4*) after exposure to MSC-exosomes (74). Upon MSC-exosome treatment, we observed a gradual increase of GFP+ cells over 5 days of treatment (**Fig. S13D**). To further confirm this result, we used flow cytometry for CD44 and CD24 to analyze the enrichment of CD44^high^/CD24^low^ CSCs from TNBC cells exposed to MSC-derived exosomes. Exposure to MSC-derived exosomes for as little as 3 days resulted in a ∼2-fold enrichment of CD44^high^/CD24^low^ CSCs, which was shown to be sufficient to shift tumor physiology toward increased aggressiveness in TNBC (**Fig. S13E**)(75). Next, we used this approach to determine whether *Oct4* deficiency can partially rescue the enrichment of CSCs in TNBC cells treated with MSC-derived exosomes. Notably, shControl-cells exposed to MSC-exosomes triggered a ∼3.5-fold enrichment of CD44^high^/CD24^low^ CSCs, which was decreased to ∼2-fold in shOct4-cells (**Fig. S13F**). These data confirmed that MSC-derived exosomes can induce the enrichment of CD44^high^/CD24^low^ CSCs, as previously shown(74).

To monitor circadian rhythms, we used our TNBC cells with a stably integrated circadian luciferase reporter (*Bmal1*-luc). Upon exposure to MSC-derived exosomes, circadian rhythms from two TNBC cell lines MDA-MB-231 and HS578T, were recorded in real time by Kronos HT, as described above (**Fig. S14A**). Circadian rhythms were disrupted after 5 to 6 days of treatment with MSC-derived exosomes (**Fig. S14, B** and **C**). Moving average of circadian oscillations were calculated at each synchronization time (**Fig. S14D**). These data suggest that the cargo of MSC-derived exosomes, which can efficiently promote dedifferentiation of TNBC cells into CSCs (74), can disrupt circadian rhythms. Consistent with the enrichment of CSCs in response to MSC-exosome exposure, TNBC cells generated larger tumorspheres (**Fig. S14, E** to **F**).

Analysis of the single cell RNA-seq (scRNA-seq) data from Sandiford et al., (74) shows the upregulation of multiple stem cell promoting pathways in TNBC cells exposed to MSC-derived exosomes (**Fig. S14G**). These stem cell promoting pathways include WNT(76), ErbB1 (also known as EGFR)(77), NF-kB(78), ALK (Activin A receptors)(79), and IL1(80). Both IL1 and NF-kB were shown to promote breast cancer progression and therapeutic resistance(80). As mentioned above, apart from its role in cancer stem cell maintenance, NF-kB pathway can repress CLOCK:BMAL1 activity (42). These cells also show an enrichment of *Per1*. Overexpression of *Per* genes disrupt circadian rhythms as they can constitutively repress CLOCK:BMAL1 transcriptional activity (40). Thus, MSC-derived exosomes reprogram the transcriptome of TNBC cells towards the disruption of circadian rhythms and the enrichment of CSCs.

Overall, these data show that circadian rhythms are disrupted in TNBC cells by carboplatin, *Bmal1* deficiency, or MSC-derived exosomes. All these paradigms resulted in transcriptomic reprogramming that is conducive to the upregulation of CSC-promoting pathways Including WNT, NF-kB, and EGFR. Notably, under conditions of *Oct4* deficiency, which restores circadian rhythms (**Fig. 4**, and **Fig. S6**), TNBC cells exposed to MSC-derived exosomes generated ∼2-fold less CD44^high^/CD24^low^ CSCs compared to shControl-cells (**Fig. S13F**). Additionally, contrary to shControl-cells, sh*Oct4*-cells exhibit robust circadian rhythms of higher amplitude upon exposure to MSC-derived exosomes (**Fig. S15**).

## DISCUSSION

Our findings indicate that loss of circadian rhythms by multiple experimental paradigms including the chemotherapeutic agent carboplatin, *Bmal1* knockdown, or MSC-derived exosomes, can facilitate the formation of CSCs. Mechanistically, using a circadian reporter (*Per2*-luc) assay, we show that core pluripotency factors including OCT4, SOX2, KLF4, and NANOG can activate circadian gene expression, Moreover, unlike CLOCK:BMAL1, activation of the circadian reporter by these pluripotency factors cannot be repressed by CRYs. The main cog of the circadian clockwork mechanism is composed of CLOCK:BMAL1-mediated transcriptional activation of circadian genes (positive feedback limb), which is repressed by PER:CRY complexes (negative feedback limb) to maintain 24-hour periodicity (81, 82). Thus, constitutive activation of circadian timekeeping genes by pluripotency factors may lead to circadian disruption. Concordantly, OCT4 overexpression results in circadian attenuation with dampened amplitude that coincides with the upregulation of the circadian negative feedback limb (PER and CRY proteins) (**Fig. 5, C, I and J**). In support of this notion, constitutive expression of PER2 was shown to disrupt circadian rhythms(40). Remarkably, the loss of circadian rhythms under carboplatin treatment was partially restored in *Oct4*-deficient TNBC cells (*shOct4*-cells) (**Fig. 4B**, and **Fig. S4C**).

Our transcriptomic analysis of TNBC cells treated with carboplatin show a reprogramming of gene expression that includes the upregulation of *Per* and *Cry* (circadian negative limb) along with the downregulation of *Npas2* and *Bmal1* (circadian positive limb). This suggests that circadian rhythm disruption is driven by the dysregulation of clockwork components, specifically through the downregulation of the positive limb and the upregulation of the negative limb. This reprogramming also shows the upregulation of genes involved in CSC-promoting pathways including EMT (*Sall4, Zeb2*, and Snai1), NF-kB (*RelA, RelB, Nfkb2, Chuk, Traf2, Traf4,* and *Traf6*), and heat-shock response (*Hspa1a, Hspa1b,* and *Hspa6*) (**Fig. S2**)(78, 83–85). Importantly, upregulation of either NF-kB pathway or the heat-shock response are associated with circadian disruption(42–44). These data support the idea of an antagonist interplay between CSC-promoting pathways and the circadian clock. Concordantly, by flow cytometry we observed that carboplatin treatment of TNBC cells stably expressing the circadian reporter Oct4-GFP, resulted in the enrichment of GFP+ cells, CD133+ cells, and CD44^high^/CD24^low^ CSCs (**Fig. 1D**). Additionally, carboplatin-treated TNBC cells, which lost their circadian rhythms, generated significantly larger tumorspheres and with higher efficiency compared to the untreated TNBC cells (**Fig. 1, F** and **G**, and **Fig. S5, A** and **B**). Unexpectedly, we observed an upregulation of genes encoding for core transcriptional pausing factors including, *Nelfa, Hexim1, Mepce,* and *Supt5h* (**Fig. S2, I** and **J**). Thus, to further understand the role of transcriptional pausing we performed PRO-seq analysis on carboplatin-treated TNBC cells. We found a significant decrease in Pol II pausing of key genes from the NF-kB pathway and the heat-shock response, both of which can promote and maintain CSCs as well as disrupt circadian rhythms (**Fig. 2**). As mentioned above, these pathways are upregulated in carboplatin-treated TNBC cells, thereby supporting the notion of an antagonistic relationship between circadian disruption and the enrichment of CSCs. Concomitantly, key timekeeping genes from the circadian positive limb including Npas2 and Pparγ exhibit increased Pol II pausing that correlated with their downregulation in carboplatin-treated TNBC cells. Remarkably, our transcriptomic analysis shows that the upregulation of CSC-promoting genes and the downregulation of key circadian timekeeping genes from the positive feedback limb including Bmal1 and *Npas2*, were restored in *Oct4*-deficient TNBC cells (*shOct4*-cells) treated with carboplatin (**Fig. S6**). Concordantly, *shOct4*-cells generated significantly smaller tumorspheres and with less efficiency compared to shControl-cells (**Fig. 4, C** and **D**, and **Fig. S5, A** and **B**). Additionally, *shOct4*-cells exhibited a significantly less invasive capacity and generated smaller solid tumors in immunodeficient mice compared to shControl-cells (**Fig. S5, E** and **F,** and **Fig. S8**).

We found that OCT4 can activate the expression of the *Per2*-luc reporter even in the presence of CRY proteins (**Fig. 3, A** and **B**). Consequently, OCT4 overexpression disrupted circadian rhythms (**Fig. 5C**) by upregulation of the negative feedback limb (PER and CRY proteins) (**Fig. 5, I** and **J**). Our ChIP-seq analysis provided us with a more defined mechanism involved in the loss of circadian rhythms in CSCs generated from carboplatin-treated TNBC cells (**Fig. 3G**). This mechanism involves two major changes that include, 1) loss of BMAL1’s cyclic binding pattern to its target genes, and 2) the binding of OCT4 to BMAL1 targeted genes. Thus, we propose a self-destructing circadian clock program relying on a constitutive BMAL1 binding pattern throughout the circadian cycle along with rhythmic OCT4 binding to disrupt the expression of circadian timekeeping genes in CSCs.

Further supporting the notion that circadian disruption facilitates the formation of CSCs, we show that *Bmal1*-deficient TNBC cells (shBmal1-cells) exhibit ∼9-fold increase of CD44^high^/CD24^low^ CSCs (**Fig. 5F**). One of the hallmarks of CSCs is their invasive capacity(31), and *shBmal1*-cells, lacking circadian rhythms, show a significant increase invasion through Matrigel compared to shControl-cells (**Fig. S5, E** and **F**). Furthermore, *shBmal1*-cells generated significantly larger tumorspheres and more efficiently than shControl-cells (**Fig.5, D** and **E**, and **Fig. S5, A** and **B**). Additionally, solid tumors from orthotopic injections of *shBmal1*-cells in immunodeficient mice expressed significantly higher levels of the CSC markers CD44, KRT7 and KRT17 (**Fig. S8, E** and **F**). Similar to carboplatin-treated TNBC cells, the transcriptome of shBmal1-cells showed an upregulation of CSC-promoting genes (CD44, KRT7 and KRT17), as well as the NF-kB pathway and the heat-shock response (**Fig. S7**).

Overall, these data suggest that formation of CSCs relies on multiple transcriptional mechanisms including, a) constitutive, rather than cyclic, binding of BMAL1 to circadian genes, b) recruitment of core pluripotency factors such as OCT4 to disrupt circadian expression, and c) dysregulated transcriptional pausing to disable the circadian clock and facilitate the expression of CSC-promoting genes. Because CSCs are recognized as tumor initiating and propagating cells capable of generating metastasis (1–3), hampering their formation by restoring circadian rhythms during chemotherapy opens new avenues for developing new cancer treatments.

Patient data from TCGA further validated the interconnection between the circadian clock and transcriptional pausing, which is disengaged in primary tumors from nearly 1000 breast cancer patients (**Fig. 6 A-E**). The uncoupling of such relationship in breast tumors supports the concept that a functional circadian clock requires a balanced pausing and unpausing of its negative and positive effectors, respectively. This underlines how an intriguing level of cellular plasticity depends on a transcriptional pausing mechanism, which alters circadian rhythms to control the transition of mature TNBC cells into CSCs within breast tumors. Furthermore, the interplay between the circadian clock and the pluripotency network including OCT4, intensifies such level of plasticity. Remarkably, analysis of nearly 10,000 patients across multiple cancer cohorts in the TCGA TARGET GTEx dataset revealed a significantly association between reduced expression of circadian clock genes *Clock*, *Bmal1* and *Npas2* and worsened clinical outcome, suggesting that disruption of the circadian clock may contribute to cancer progression (**Fig. S10**). Moreover, the regulatory interconnection between transcriptional pausing and the circadian clock we elucidated throughout this research is disrupted in patients from multiple cancer types and correlates with low survival (**Fig. S11B**). This suggests a more universal role for the disruption of circadian-transcriptional pausing partnership across diverse cancer types. In support of this view, we found a cell type derived from human osteosarcoma (U2OS) that lost their circadian rhythms upon carboplatin-mediated reprogramming towards CSCs (**Fig. S1**, **D** and **E**). Although chemotherapeutic drugs are effective on targeting cancer cells, they are also effectively promoting the development of CSCs, which have the capacity of invading distal tissues and initiate metastatic lesions(8). We envision that new treatments capable of restoring circadian cycles will avert the formation of CSCs. However, restoration of circadian rhythms to abolish CSCs might not apply to all cancer types since CLOCK and BMAL1 were shown to be required for the optimal growth of glioblastoma stem cells(86) and acute myeloid leukemia stem cells(87).

We revealed that exosomes derived from MSCs, which were shown to promote the enrichment of CSCs (74), can disrupt circadian rhythms in TNBC cells (**Fig. S14, B** and **C**). Our analysis from single cell RNA-seq (74) show un upregulation of the CSC-promoting pathways WNT and NF-kB (32, 88), which are interconnected with the circadian clock(42, 89). Importantly, TNBC patients undergoing chemotherapeutic treatment could be negatively affected by both MSC-derived exosomes and chemotherapeutic drugs, which can trigger the loss of circadian cycles and promote the development of CSCs. Compared to other breast cancer types, TNBC has a higher risk of early recurrence and metastasis. Cancer recurrence happens in about 50% of early stage TNBC patients with an almost 40% mortality within the first 5 years of treatment(90). This poor prognosis could be enabled by dual effects from MSC-derived exosomes combined with chemotherapy, both of which facilitate the loss of circadian rhythms to promote the formation of CSCs capable of augmenting recurrence in TNBC patients. However, how the loss of circadian rhythms impacts the poor prognosis of TNBC has remained unexplored. The data in this manuscript shed light into the molecular mechanisms underlying the loss of circadian rhythms in TNBC cells in response to the chemotherapeutic agent carboplatin, which promotes the enrichment of CSCs. Thus, our findings could lead to the development of new therapies that incorporate restoration of circadian rhythms to avert formation and maintenance of CSCs and consequently improve the prognosis of TNBC patients.

Overall, our molecular findings indicate that transcriptional pausing in combination with the pluripotency network can govern circadian rhythms, which in turn act as a key regulator for the formation of CSCs. Concordantly, the status of such interplay between the circadian clock, transcriptional pausing, and pluripotency becomes a molecular signature that distinguishes normal tissues from primary tumors affecting survivability of cancer patients. Thus, restoration of this interdependent circadian framework could prevent the formation of CSCs in TNBC and possibly other cancer types, thereby limiting downstream consequences such as therapeutic resistance and relapse.

## METHODS

### Cell Culture

Triple negative MDA MB-231 (BCCs) with a transgenic *Oct4*-GFP reporter were obtain from Dr. Pranela Rameshwar’s laboratory. BCCs or HS578T (ATCC: HTB-126) were grown on DMEM [DMEM-high glucose, 4500 mg/L and sodium bicarbonate, 10% FBS, 1x MEM Non-essential Amino Acid Solution, 10 mM HEPES solution, 1 mM Sodium Pyruvate solution, 2 mM L-Glutamine solution, antibiotics (100 I.U./ml-Penicillin, 100 µg/ml-Streptomycin), 100 µg/ml Primocin]. For integration of the *Bmal1*-luciferase (*Bmal1*-luc) (Addgene: 68833) or Per2-luciferase (Per2-luc) (Addgene: 212035) reporter, BCCs/HS578T were infected with filtered viral supernatant from HEK293T cells transfected with the reporter plasmid and ViraPower (Invitrogen Cat: K497500) lentiviral packaging system. Following transduction, BCCs were selected with blasticidin (Gibco Cat: A11139-03) (5000x, 2ug/ml) for 3-5 days. OneGlo (Promega, Cat: E6120) bioluminescence assay was used to confirm integration of the *Bmal1*-luc/Per2-luc reporter in BCCs. For carboplatin treatment, BCCs were treated with Carboplatin (Sigma-Aldrich Cat: C2538) at 50 ug/ml or 100 ug/ml resuspended in 5% glucose solution. Generation of shRNA knockdowns and CLOCK over expression (GeneCopoeia) and OCT4 over expression (Addgene: 16579)(91) were done by infecting BCCs-*Bmal1*-Luc cells with lentiviral particles. Transduced cells were then selected with Puromycin (Sigma-Aldrich Cat: P9620) for 3 days. For circadian synchronization, cells were treated with 100 nM dexamethasone for 2 hours at 37°C or by media change (92).

### Cell viability assay

Each cell line was seeded onto 96 well plate (n=4) at a concentration of 1 or 2×10^3^ cells. XTT assay (ATTC, Cat: 30-1011K) was done every day according to the manufacturer’s specifications within a period of 4 days. A plate reader was used to measure absorbance values of the colorimetric assay (475nm and 660nm) and background values were subtracted from experimental values.

### Exosome Preparation, Isolation, and Quantification

Immortalized human mesenchymal stem cells (hTert MSCs) were used to collect and isolate exosomes. To isolate exosomes, 1*10^6^ hTert MSCs were seeded on multiple 10cm dishes with exosome free FBS. Media was collected and replenished every 2-3 days for a week(74). The media was subjected to serial differential centrifugation(74, 93). Briefly, the first spin was done at 2000xg for 20 minutes to get rid of dead cells and large debris, the second spin was done at 10,000xg 30-45 minutes to get rid of large vesicles, Subsequently, the third spin was done at 100,000xg for 90 minutes on a Beckman preparative ultracentrifuge (Type 50.2 TI Fixed angle Rotor) to spin down the exosomes, and a final spin was done at 130,000xg for 90 min on a tabletop Beckman preparative ultra centrifuge (TLA 100.3 Fixed angle rotor) after to pellet the exosomes after washing with PBS. Exosomes were resuspended in 300ul of PBS and quantified via nanoparticle tracking analysis using the Nanosight LM10 system at a dilution factor of 1:1000.

### Western Blotting

Whole cell extracts were obtained by lysing cells in an SDS-based buffer (2% SDS, 150 mM NaCl, 50 mM Tris-HCl pH 7.0, 5 mM EDTA plus protease inhibitors) followed by sonication. Nuclei for nuclear and chromatin extracts were isolated by resuspending cell pellets in Cellular Lysis Buffer (10 mM HEPES, 10 mM KCl, 1% IGEPAL plus protease inhibitors) and incubating for 20 minutes on ice followed by centrifugation at 21,000 x g. The supernatant was saved as the cytoplasmic fraction, and the pellet (isolated nuclei) were lysed in SDS based buffer, mentioned above with protease inhibitors, followed by sonication for nuclear extracts. Chromatin extracts were obtained by lysing nuclei with 0.2M HCl for 20 minutes on ice followed by centrifugation at 21,000 xg for 10 minutes at 4°C. The supernatant containing soluble chromatin bound proteins was collected and neutralized with an equal volume of Tris-HCl pH 8. Quantification of protein concentration was done using Pierce BCA Protein Assay Kit (Cat: 23225). Briefly, 10-30 ug of protein were run on 8%-12% polyacrylamide gels or 4-20% gradient gels (Bio-Rad) and transferred on to either polyvinylidene difluoride (PVDF) or Nitrocellulose membranes. Membranes were blocked using Membrane Blocking Solution (Invitrogen Cat: 000105) or Everyblot Blocking Buffer (Bio-Rad Cat: 12010020) for 1 hour and probed with primary antibodies overnight. Proteins were visualized using horseradish-peroxidase conjugated secondary antibodies (Bio-Rad) and SuperSignal West Femto Maximum Sensitivity Substrate (Thermo Scientific, Cat: 34096) or StarBright Blue 700/520 fluorescently conjugated secondary antibodies (Bio-Rad) on the ChemiDoc MP analyzer from Bio-Rad. Protein levels quantification was done by measuring band intensity using the Bio-rad ImageLab software and normalizing to GAPDH or Histone H3 loading controls. The following antibodies were used: ALDHA1 (CST, 54135S, 1:1000), b-CATENIN (BD Biosciences, 610154, 1:1000), BMAL1 (CST, 14020S, 1:1000), CLOCK (CST, 5157S, 1:1000), COBRA1/NELF-B (CST, 14894S, 1:1000), CRY1 (ABClonal, A13662, 1:1000), CRY2 (Proteintech, 13997-1-AP, 1:1000), HEXIM1 (CST, 12604S, 1:1000), KLF4 (CST, 12173S, 1:1000), OCT3/4 (SantaCruz, sc-5279, 1:1000), Oct4 (Proteintech, 11263-1-AP, 1:1000), PER2 (ABClonal, A5107, 1:1000), GAPDH (Proteintech, 60004-1-Ig, 1:10000), hFAB Rhodamine GAPDH (Bio-Rad, 12004168, 1:5000), GAPDH (BioVision, 3777R, 1:3000), and Histone H3 (CST, 14269S or 4499, 1:10000).

### RT-qPCR

Total RNA was isolated using TriPure Isolation Reagent (Roche Cat: 11667165001) according to manufacturer specifications. The RNA pellet was resuspended in DEPC water and quantified by Nanodrop. cDNA was made according to manufacturer’s instructions using LunaScript RT SuperMix (NEB Cat M3010) starting with 1 mg of RNA. Resulting cDNA was diluted 10-fold and 1ml was used for each reaction. qPCR was performed on the CFX384 Touch Real-Time PCR Detection System from BioRad using Luna Universal qPCR Master Mix (NEB Cat: M3003X). The following qPCR cycles settings were used, initial denaturing step at 95°C for 3 minutes, followed by 40 amplification cycles at 95°C for 30 seconds, 60°C for 30 seconds, and a melting curve after the 40 amplification cycles were completed. Real time Ct values for each target gene were normalized to *Gapdh* to obtain DCt values whereby fold change was calculated as equal to 2 - DCt. The following primers for *OCT3/4* 5’-CTTGCTGCAGAAGTGGGTGGAGGAA-3’ (Forward) and 5’-CTGCAGTGTGGGTTTCGGGCA-3’ (reverse) and *GAPDH* 5’-GCCTCAAGATCATCATCAGCAATGCCT-3’ (Forward), 5’-TGTGGTCATGAGTCCTTCCACGAT3’ (Reverse) were used (94), *SNAI1* 5’-CTCTAATCCAGAGTTTACCTTC-3’ (Forward) and 5’-GACAGAGTCCCAGATGAG-3’ (reverse), *KRT19* 5’-GCGAGCTAGAGGTGAAGATC-3’ (Forward) and 5’-CGGAAGTCATCTGCAGCCA-3’ (Reverse), *ZEB1* 5′-TTCACAGTGGAGAGAAGCCA-3’ (Forward) and 5′-TTCACAGTGGAGAGAAGCCA-3’ (Reverse), *ABCG2* 5′-GCCACAGAGATCATAGAGCCT-3′ (Forward) and 5′-TCACCCCCGGAAAGTTGATG-3′ (Reverse), *NANOG* 5′-GAGAAGAGTGTCGCAAAAAAGGA-3′ (Forward) and 5′-TGAGGTTCAGGATGTTGGAGAGT-3’ (Reverse), *CD44* 5’-CGCAGATCGATTTGAATATAACC-3’ (Forward) and 5’-CCGATGCTCAGAGCTTTCTC-3’ (Reverse), *Per2* 5’-CTG TCA CCA CCA TAG AAA GG-3’ (Forward) and 5’-CAA GGA GGC TGG TTC TTA TAG-3’ (Reverse), *Cry2* 5’-GTG GTA GTA CCC TAG CTC AA-3’ (Forward) and 5’-GCT GCC TCT GAA ACT CTA TG-3’ (Reverse), *NR1D1* 5’-ATC CTC CTC CTC CTT CTA TAA C-3’ (Forward) and 5’-GCT TGG TAA TGT TGC TTG TG-3’ (Reverse), *RORA* 5’-GAC AGA CCA TCG CTT TAG TT-3’ (Forward) and 5’-CTA AGC TTC TCT GGC CTT ATG-3’ (Reverse), *CLOCK* 5’-GTG ATG GTG AGG TGC TAA A-3’ (Forward) and 5’-GAG CAA AGC TAG AGA CAG ATA G-3’ (Reverse), *BMAL1* 5’-GTT TCT CGA CAC GCA ATA GA-3’ (Forward) and 5’-CCT AGA AGT TCC TGT GGT AGA-3’ (Reverse).

### Dual-Luciferase Assay for Transient Transfections

HEK293T cells were seeded at 2×10^4^ cells per 96 well in a quadruplicate and incubated overnight. The next day cells were transfected with 50 ng of *Per2-*luc (Addgene:110059) (95) and 2.5 ng of Renilla-Luciferase (Promega), with or without cDNAs for circadian activators (CLOCK (Addgene: 47334) (96) and BMAL1 (Addgene: 31367) (97)), circadian repressors (CRY1 (Addgene: 25843) and CRY2 (Addgene: 25842) (98)), or pluripotency genes (OCT4 (Addgene: 26816), SOX2 (Addgene: 26817), KLF4 (Addgene: 26815), MYC (Addgene: 26818) (99), NANOG (Addgene: 28221) (100)) using 1 mg/mL PEI MAX (Polysciences, Cat: 24765) at 1:2 DNA:PEI ratio. Twenty-four hours after transfection, cells were lysed followed by luciferase assay using Dual-luciferase Reporter Assay (Promega Cat: E1960) according to the manufacturer’s instructions.

### Real-time Monitoring of Circadian Rhythms

A day prior to real-time bioluminescent monitoring using KronosHT (Atto), cells were seeded on white optical 96 well or 24 well plates at 4×10^5^ cells and cultured on DMEM. The following day, cells were washed with PBS, and cultured with Circadian Media [DMEM without phenol red, 10% FBS, 3.5 g/L glucose, 350 mg/L Sodium Bicarbonate, 1x MEM Non-essential Amino Acid Solution, 10 mM HEPES solution, 1 mM Sodium Pyruvate solution, 2 mM L-Glutamine solution, antibiotics (100 I.U./ml Penicillin,100 µg/ml Streptomycin), 100 µg/ml Primocin] with Luciferin (Promega, Cat: L2916) (500x, 0.1mM) (101). Circadian rhythms were monitored over a 6-12-day period, recording bioluminescence for 5 seconds in 16-minute intervals per each well for the 96 well or 10 seconds in 6-minute intervals per well for the 24 well. To resynchronize the cells, dexamethasone treatment, or media changes, were done every 3 days until completion of the experiment. The data were detrended using the KronosHT software, following a +/− 12hr width, and smoothed following a +/−1 point moving average and median filter. Detrended data was exported, and graphical representations were made using KronoAnalyzer (Atto) or Excel.

### Mathematical analysis of circadian rhythms from real-time profiles

To capture the circadian rhythm 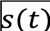 in the data a model of the form

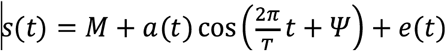

is fitted to the data as described in Indic & Brown (102). 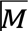 is the MESOR, 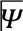 is the acrophase, 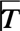 is the period and 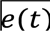 is the error term with 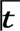in hour.

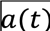 captures the change in amplitude observed in the data which is expressed as 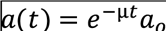 where µ representing the change in amplitude per hour and 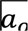 as the initial amplitude. With 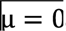, the equation becomes the cosinor-based model that has been widely used for finding circadian rhythm in chronobiology (103).

If 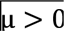, the rhythm exhibits a decay and if 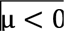 the rhythm exhibits a growth. The parameters, 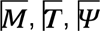, 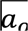 and 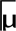 are estimated using a nonlinear optimization procedure.

### 3D Tumorsphere Formation

Cells were seeded into U-shaped 96 well plates at a concentration of 5×10^3^ in 100 ul of media for 5 days. Images of tumorsphere morphology were taken daily over the 5-day period. The size of the tumorpheres was assessed by measuring the surface area of the images using Zeiss microscopy software Zen Blue.

### Tumor formation efficiency assay

Single cell BCCs (1×10^3^) were seeded in a 6 well of an ultra-low attachment plate (Corning) in 3D Tumorsphere Medium XF (PromoCell) supplemented with 20 ng/mL bFGF and EGF (PeproTech), the media was made semisolid with addition of 0.5% methylcellulose to prevent cell aggregation and incubated for 12 days. After 12 days, the semisolid media was diluted cells gathered to the middle of the well and pictures were taken for tumor-sphere count. Tumorsphere Formation Efficiency % was calculated as (# of tumorspheres formed/initial cells seed) x 100%.

### Coverslip Preparation and Treatment

Coverslips were coated with diluted Matrigel (1:300) (Corning, Cat: 354277) for 1 hour in a 24 well plate. Cells were seeded at a density of 3.5×10^4^ cells per well. The following day after attachment, the cells were treated with 100ug/ml Carboplatin for 5 days. Seeded coverslips were fixed with 4% Formaldehyde in PBS for 12 minutes and then washed twice in 0.1% PBS with Triton-X. Fixed coverslips were mounted on with Prolong-Gold Antifade Mountant with DNA Stain DAPI (Invitrogen, P36931) on to slides.

### Invasion Assay

Corning 24-well 5µm transwell inserts (Corning Cat: 3421) were coated with 300ug/ml of Corning Matrigel (Corning Cat: 354234) diluted in sterile Coating Buffer (0.01M Tris (pH 8.0), 0.7% NaCl) for 2 hours at 37C. Residual coating buffer was aspirated, and single cells were resuspended in FBS free DMEM and seeded in the inserts at a density of 2×10^4^ cells per insert. The well received 750µl of DMEM with FBS as chemoattractant. Plates were incubated at 37C for 3 days, then fixed in 4% formaldehyde in PBS for 10 minutes and then washed twice with PBS. Fixed transwells were stained with 3ug/ml Hoechest (Invitrogen Cat: H3570) in 0.05% PBS with Triton-X for 10 minutes and washed twice with 0.05% PBS with Triton-X. Transwells were imaged with a 5x objective lens after staining and then swabbed with cotton applicator swabs. Transwells were washed twice again with 0.05% PBS with Triton-X and imaged again. Images were quantified and percentage of invading cells was determined by dividing the quantity of cells after swabbing by the quantity of cells before swabbing.

### Imaging and Quantification

Images were taken on a Zeiss Axio Observer.Z1 / 7 using a 5x or 20x objective lens. GFP filter (41% laser intensity, 1.57s exposure), and DAPI filter (50% laser intensity, 500ms exposure) were used. All image quantification was done using ImageJ. A threshold was applied to the image, followed by processing by applying median (radius =3) and watershed. The analyze particle tool (size 100 – inf: pixel units, circularity 0.25 – 1.00) was used to quantify the mean fluorescent intensity or quantity cells of each image.

### Flow Cytometry

To identify cancer stem cell, cells were stained for the cell surface markers CD44, CD24, and CD133. 2×10^5^ cells were resuspended in FACS Sorting Buffer (0.9% FBS, 1mM EDTA in PBS) and blocked with FC antibody (BD Biosciences, Cat: 564219) for 10 minutes on ice. Cells were centrifuged at 200xg for 5 minutes at 4C°. Supernatant was removed and cells were resuspended in FACS Sorting Buffer with CD44-PerCP-Cy 5.5 conjugated (Cell Signaling Technologies, Cat: 560531) CD24-PE-Cy7 conjugated (BD Pharmingen, Cat: 561646) and CD133-BV421 conjugated (BioLegend, Cat: 372807) antibodies (1:1200) and incubated on ice for 30-40 minutes. Cells were centrifuged again and resuspended in FACS Sorting Buffer and profiled using the BD LSRFortessa X-20 at the NJMS Flow Cytometry and Immunology Core Laboratory. For cell cycle analysis, live-cells were stained with 2.5 ug/ml of Hoechest 33342 (Cat# 62249, ThermoScientific) for 30 minutes. Cells were then pelleted, resuspended in FACS buffer with Hoechst and profiled same as above.

### RNA-seq

Total RNA was extracted using RNeasy Mini Kit (Qiagen, Cat: 74104) following manufacture instructions. RNA was quantified using nanodrop prior sequencing. RNA-seq was performed at PrimBio Research Institute through Ion Torrent and at NJMS genomic center through illumina. For RNA-seq performed at PrimeBio, Mammalian Ribo-off rRNA removal kit was used to remove rRNA from approximately 1ug of total RNA following the manufactures recommended protocol. Library preparation is done with Ion Total RNA-seq Kit v2 from Life Technologies with manufacturers recommended protocol. For RNA-seq performed at NJMS, libraries were prepared using NEBNext Ultra II DNA Library prep Kit for Illumina (New England Biolabs, Catalog number E7645) according to manufacturer protocol. The Adaptor Ligation reaction was performed with the Illumina Adaptor diluted 10-fold (1:10). Adaptor-ligated DNA was cleaned-up without size selection (DNA input ≤ 50 ng) with AMPure XP beads (Beckman Coulterm Inc. Catalog number A63881). PCR Enrichment of Adaptor ligated DNA was carried out using NEBNext Multiplex Oligos for Illumina (Dual Index Primers Set 1) (New England Biolabs, Catalog number E7600S) with 7 cycles of Denaturation and Annealing/Extension. Library fragment size distribution was checked with D1000 ScreenTape assay for TapeStation systems (Agilent), and library concentration was determined using Qubit 1X dsDNA high sensitivity (Invitrogen).

### RNA-seq analysis

Bulk RNA-seq data from NJMS genomic center on Carboplatin treated and untreated shKD MDA-MB-231 samples were analyzed through Nf-Core(104) RNA-seq pipeline with singularity. (r 3.12.0, remove_ribo_rna, --profile_singularity). Briefly, raw reads were trimmed with Trimgalore! and rRNA were filtered with sortMeRNA. Fragments were aligned to hg38 reference sequence through STAR, and quantification were done through Salmon. Count matrix from nf-core RNA-seq pipeline was then processed through Nf-Core Differentialabundance pipeline. (--filter_min_abundance 10, --filter_min_samples 6, --differential_min_fold_change 1.5, --differential_max_pval 0.05, r 1.2.0). Pathway analysis was done through GSEA or Enrichr, and 5 pathways of interest are selected to construct bar graph. Bar graph for Pathway analysis visualization and volcano plots were constructed through prism9. Pseudo read count were generated from Morpheus for exosome RNA-seq, and pathway analysis were done through Enrichr.

### CIRCust timepoint assessment

The circadian activity of Bulk RNA-seq samples was assessed using CIRCust(72). To validate their rhythmic activity, time-stamped MCF10A samples from a previous study(105) were analyzed with CIRCust. The Circular Principal Component Analysis (CPCA) was then employed on the MCF10A, and the score matrix derived from CPCA was used to locate time-stamped sample’s location on a unit circle.

Assessment of the Circadian activity of breast cancer patient sample in TCGA were done through CIRCust(72). Briefly, CIRCust(72) excluded low-expression genes after normalization. Circular Principal Component Analysis (CPCA) was then performed using a CIRCust defined list of core circadian genes. The loading matrix derived from the CPCA analysis was then employed to construct a unit circle, onto which potential epigenes were projected to determine their respective phases. FMM model is then used to reorder the expression data, followed by a R2-based goodness-of-fit check to assess the fitness of the FMM model for each gene.

### Cancer genome ATLAS analysis

The TCGA TARGET GTEX reanalyze data were downloaded from the USCS Xena Portal and analyzed in RStudio(106). The FPKM normalized dataset was used to extract all adult breast cancer patients to create a large cohort (n = 1091). A smaller subset was created for all adult breast cancer patients with both primary tumor and normal tissue sample. PCA was used to calculate covariance between genes of interest in the large and small patient cohorts, and patient were separated into high-covariance and low-covariance subset based on their contribution to PCA. High and low correlation (covariance) groups were identified using a PCA-based feature selection approach focused on pathway of interest. Principal component analysis (PCA) was performed on the full patient expression dataset to capture dominant patterns of co-variation among target genes. The first three principal components (PC1–PC3), which explained the largest proportion of variance, were selected for analysis. Each sample’s position (projection) in the reduced PCA space was used to quantify its overall contribution to the gene-gene correlation structure within the pathway of interest. Samples with higher projections—indicating stronger alignment with the major axes of gene expression correlation—were classified as the high-correlation group, while those with lower projections were assigned to the low-correlation group. Essentially, samples with high projections (reflecting stronger coordinated expression of the target genes) were defined as “high correlation,” while those with weaker projection patterns were defined as “low correlation”.

Expression levels of genes of interest were used to stratify adult cancer patients (∼10,000 patients) from the TCGA TARGET GTEx dataset available through UCSC Xena. Patient survival data from the TCGA database were used to assess survival differences among patient subgroups using either GraphPad Prism 9 or the UCSC Xena Browser. Bar graphs representing circadian gene activity were generated using GraphPad Prism 9. Spearman correlation matrices were clustered using Clustergrammer2 with Euclidean distance and complete linkage clustering. Patient treatment information was retrieved using the GDCquery function from the TCGAbiolinks R package(107–109). Patients treated with platinum-based drugs were selected and stratified according to the expression level of the gene of interest.

### Chromatin Immunoprecipitation (ChIP)

Dexamethosone synchronized untreated and carboplatin treated (50 ug/ml for 5 days) BCCs were crosslinked with 1% formaldehyde in PBS with Ca2+/Mg2+ for 10 minutes at room temperature followed by quenching with glycine 0.125 M (final concentration) for 5 minutes on a shaker at timepoints 9hrs and 18hrs. Crosslinked cells were washed 3x with ice cold PBS and scraped off the plate in PBS to be centrifuged at 200 x g for 10 minutes at 4°C. Following centrifugation, crosslinked cell pellets were lysed in Cellular Lysis Buffer (5 mM PIPES. pH 8.0, 85 mM KCl, 0.5% IGEPAL CA-630) with protease inhibitors (Pierce Cat: A32955) on ice for 5 minutes followed by short centrifugation at 200 x g for 2 minutes. The supernatant was removed, and the remaining pellet was resuspended in Nuclear Lysis Buffer (50 mM Tris-HCl pH 8.0, 10 mM EDTA pH 8.0, 0.2% SDS) with protease inhibitors followed by sonication using Qsonica Q800R Sonicator for 20 minutes (30% Amplitude, 20 sec ON, 20 sec OFF, at 4°C). After sonication, samples were centrifuged on a tabletop centrifuge at 21,000 x g to remove cell debris, and 5 ul of chromatin was set aside to confirm sonication efficiency. The sonication test was performed by adding Elution Buffer (50 mM NaHCO_3_, 140 mM NaCl, 1% SDS) plus Proteinase K (Promega Cat: MC5005) and incubated overnight at 65°C. RNAse A (Qiagen Cat: 1007885) was added to each chromatin input and incubated for 30 minutes at 37°C. Input samples were subjected to phenol-chloroform extraction and sonicated DNA size was confirmed by 1% agarose gel electrophoresis. DNA fragments were below 600 bp.

For immunoprecipitations (IPs), antibodies for BMAL1 (Proteintech, Cat: 14268-1-AP), OCT4 (SantaCruz, Cat: sc-5279x) or Rabbit IgG (Invitrogen, Cat 026102) was conjugated to magnetic dynabeads Protein A or Protein G (Invitrogen Cat: 10006D/10007D) in Dilution IP Buffer (16.7 mM Tris-HCl pH 8.0, 1.2 mM ETDA pH 8.0, 167 mM NaCl, 0.1% SDS, 0.24% Triton-×100) in the presence of protease inhibitors for 6 hours at 4°C. Conjugated beads were washed twice with Dilution IP Buffer and resuspended on the same buffer. Chromatin was precleared for 1 hour twice with Protein A Agarose beads (Cell Signaling Technology, Cat: 9863S) prior to the IP. Approximately 20ug of precleared chromatin was added per each IP and incubated at 4°C with end-to-end rotation overnight. An aliquot of 10% of non-precleared material was set aside as input per each IP. Next day, conjugated beads were washed with cold Dilution IP Buffer twice, once with room temperature TSE Buffer (20 mM Tris-HCl pH 8.0, 2 mM EDTA pH 8.0, 500 mM NaCl, 1% Triton-×100, 0.01% SDS), once with room temperature LiCl Buffer (100 mM Tris-HCl pH8.0, 500 mM LiCl, 1% sodium deoxycholic acid, 1% IGEPAL CA-630), and twice with room temperature TE Buffer (10 mM Tris-HCl pH 8.0, 1 mM EDTA pH 8.0). Samples were then incubated in Elution Buffer containing RNAse A at 37°C for 30 minutes followed by incubation with Proteinase-K at 56°C for 1 hour. Inputs were treated the same as IPs after completion of the washing steps. Conjugated magnetic beads were removed and the elute was de-crosslinked by incubation at 65°C overnight on a shaker. Precipitated DNA was purified using Qiagen PCR Purification Kit (Qiagen Cat: 28106)and eluted in Elution Buffer. Purified DNA was quantified using Qubit and sent for sequencing.

### ChIP-seq analysis

Sequencing of purified ChIPs were done by Novogene. Raw sequencing data from Novogene were processed using the nf-core ChIP-seq pipeline on the Rutgers Amarel High Performance Computing (HPC) cluster with Singularity. Briefly, reads were trimmed using TrimGalore! and aligned to the GRCh38 reference genome with BWA. PCR duplicates were marked using Picard, and read filtering was performed with SAMtools and BedTools. Peak calling and signal track generation were performed using MACS3. The callpeak command with the --bdg flag was used to generate target and lambda bedgraph files, which were then processed with bdgpeakcall for peak identification and bdgcmp with the --logFE flag to generate fold-change signal tracks in BigWig format.

### PRO-seq

Untreated and carboplatin treated BCCs were collected and washed with PBS. Subsequently, cells were resuspended in Buffer W (10 mM Tris-HCl pH 8.0, 10% glycerol, 250 mM sucrose, 10 mM KCl, 5 mM MgCl2, 0.5 mM DTT, protease inhibitors cocktail) at a cell density of 2 × 10^6^ cells/mL. Then, 9x volume of Buffer P (10 mM Tris-HCl pH 8.0, 10% glycerol, 250 mM sucrose, 10 mM KCl, 5 mM MgCl2, 0.5 mM DTT, 0.1% Igepal, protease inhibitors cocktail) was added to each sample. These cell suspensions were homogenized and incubated for up to 2 min on ice. Cells were then centrifuged at 800 x g for 5 min and resuspended in Buffer F [50 mM Tris-HCl pH 8.0, 40% glycerol, 5 mM MgCl2, 0.5 mM DTT, 4 u/mL RNase inhibitor (SUPERaseIN, Ambion)] at a density of 2×10^6^ cells/100 μL. Permeabilized cells were resuspended in Buffer F and then passed through a cell strainer to assure a single cell suspension, and immediately frozen in liquid nitrogen. Permeabilized cells were stored in −80°C until ready for PRO-seq experiments. PRO-seq experiments and analyzes were done at the Nascent Transcriptomics Core at Harvard Medical School. Aliquots of frozen (−80°C) permeabilized cells were thawed on ice and pipetted gently to fully resuspend. Aliquots were removed and permeabilized cells were counted using a Luna II, Logos Biosystems instrument. For each sample, 1 million permeabilized cells were used for nuclear run-on, with 50,000 permeabilized Drosophila S2 cells added to each sample for normalization. Nuclear run on assays and library preparation were performed essentially as described in Reimer et al.(110) [with modifications noted: 2X nuclear run-on buffer consisted of (10 mM Tris (pH 8), 10 mM MgCl2, 1 mM DTT, 300mM KCl, 20uM/ea biotin-11-NTPs (Perkin Elmer), 0.8U/uL SuperaseIN (Thermo), 1% sarkosyl). Run-on reactions were performed at 37°C. Adenylated 3&#39; adapter was prepared using the 5&#39; DNA adenylation kit (NEB) and ligated using T4 RNA ligase 2, truncated KQ (NEB, per manufacturer’s instructions with 15% PEG-8000 final) and incubated at 16°C overnight. 180uL of betaine buffer (1.42g of betaine brought to 10mL) was mixed with ligations and incubated 5 min at 65°C and 2 min on ice prior to addition of streptavidin beads. After T4 polynucleotide kinase (NEB) treatment, beads were washed once each with high salt, low salt, and 0.25X T4 RNA ligase buffer (NEB) and resuspended in 5&#39; adapter mix (10 pmol 5&#39; adapter, 30 pmol blocking oligo, water). 5&#39; adapter ligation was per Reimer but with 15% PEG-8000 final. Eluted cDNA was amplified 5-cycles (NEBNext Ultra II Q5 master mix (NEB) with Illumina TruSeq PCR primers RP-1 and RPI-X) following the manufacturer&#39;’s suggested cycling protocol for library construction. A portion of preCR was serially diluted and for test amplification to determine optimal amplification of final libraries. Pooled libraries were sequenced using the llumina NovaSeq platform.

### PRO-seq data analysis

All custom scripts described herein are available on the AdelmanLab GitHub (https://github.com/AdelmanLab/NIH_scripts). Read pairs were trimmed using cutadapt 1.14 to remove adapter sequences (-O 1 --match-read-wildcards -m {20}). An additional nucleotide was removed from the end of read 1 (R1), using seqtk trimfq (https://github.com/lh3/seqtk), to preserve a single mate orientation during alignment. The paired end reads were then mapped to a combined genome index, including both the spike (dm6) and primary (hg38) genomes, using bowtie2 [10.1038/nmeth.1923]. Properly paired reads were retained. These read pairs were then separated based on the genome (i.e. spike-in vs primary) to which they mapped. Reads mapping to the reference genome were separated according to whether they were R1 or R2, sorted via samtools 1.3.1 (-n), and subsequently converted to bedGraph format using a custom script (bowtie2stdBedGraph.pl). We note that this script counts each read once at the exact 3’ end of the nascent RNA. Because R1 in PRO-seq reveals the position of the RNA 3’ end, the “+” and “-” strands were swapped to generate bedGraphs representing 3’ end positions at single nucleotide resolution. Combined bedGraphs were generated by summing counts per nucleotide across replicates for each condition.

Annotated transcription start sites were obtained from human (GRCh38.99) GTFs from Ensembl. After removing transcripts with {immunoglobulin, Mt, Mt_tRNA, rRNA} biotypes, PRO-seq signal in each sample was calculated in the window from the annotated TSS to +150 nt downstream, using a custom script, make_heatmap.pl. This script counts each read one time, at the exact 3’ end location of the nascent RNA. Given good agreement between replicates and similar return of spike-in reads within conditions, bedGraphs were merged within conditions, and spike normalized, to generate bigWig files binned at 10bp.

### Refinement of gene annotation (GGA) using PRO-seq and RNA-seq

Paired-end RNA-seq reads were mapped to the hg38 reference genome via HISAT2 v2.2.1 (--known-splicesite-infile). To select gene-level features for differential expression analysis, and for pairing with PRO-seq data, we assigned a single, dominant TSS and transcription end site (TES) to each active gene. This was accomplished using a custom script, get_gene_annotations.sh (available at https://github.com/AdelmanLab/GetGeneAnnotation_GGA), which uses RNA-seq read abundance and PRO-seq R2 reads (RNA 5’ ends) to identify dominant TSSs, and RNA-seq profiles to define most commonly used TESs. RNA-seq and PRO-seq data from all conditions were used for this analysis, to comprehensively capture gene activity in these samples.

### Differential expression analysis

Reads were summed within the TSS to TES window for each active gene using the using the “make_heatmap” script (https://github.com/AdelmanLab/NIH_scripts), which counts each read one time, at the exact 3’ end location of the nascent RNA. DEseq2, using the Wald test, was used to determine statistically significant differentially expressed genes. Unless otherwise noted, the default size factors determined by DEseq2 were used.

### Metagene analysis

Average metagene plots were generated by summing reads within bins centered at each indicated position with respect to the TSS and dividing by the number of TSSs included in each group. Bin size is indicated in each Figure legend.

### Pausing Indices

Read counts were summed in the window from TSS to +200 for each active gene, and from +250 downstream of the TSS to the TES, using the refined annotations described above. The ratio of promoter proximal signal (TSS to +150) over gene body signal (+250 to TES) is the Pausing Index.

### Orthotopic Mammary Fat Pad Injection of NSG Mice and Tumor IHC Staining

For orthotopic mammary fat pad injections we protocol was followed as per Zhang et al.(111). In brief, NSG mice were purchased from Jax Labs at 8 weeks and cells were prepared at a density of 1*10^7^cells/ml in equal volumes of PBS and Matrigel and kept on ice prior to injections. Mice were anesthetized in accordance with IACUC with 2% isoflurane and toe pinch was performed to ensure anesthesia. Fur was shaved off of the belly near the mammary fat pad followed by application of depilatory cream for 30-45 seconds that was then wiped off with a damp KimWipe. The area was sterilized with 70% ethanol and approximately 1*10^6^ cells were injected orthotopically in the mammary fat pad of the mice. Mice were allowed to recover in a warming chamber prior to being placed into isolated cages for monitoring daily. Mice weight was assessed daily and tumor size was assessed weekly using a caliper after day 7. To isolate the tumor, mice were euthanized using isoflurane and the tumor was excised from the mouse. The tumor was placed into 10% formalin buffer for fixation overnight.

Tumor sections from paraffin-embedded tissue blocks were sectioned at 5 μm and incubated on a heat block overnight prior to immunohistochemistry (IHC) staining. Briefly, sections were warmed to 55 °C and subjected to xylene washes for deparaffinization. Tissues were then rehydrated through graded ethanol washes followed by antigen retrieval using Tris-EDTA buffer in a pressure cooker for 5 minutes. Endogenous peroxidase activity was quenched with 3% H_2_O_2_, and sections were incubated with primary antibodies overnight at 4 °C. Following primary antibody incubation, sections were washed with PBS and incubated with biotinylated secondary antibodies, followed by HRP-ABC reagent from Vector Laboratories. Chromogenic detection was performed using a DAB substrate kit from Vector Laboratories, and tissues were counterstained with hematoxylin prior to mounting.

### Statistical analysis

Tumorsphere, ChIP-qPCR, western blot, wound healing and luciferase reporter assay were analyzed through pair-end T-test. For PRO-seq, statistical significance for comparisons was assessed by Wilcoxon (unpaired) or Mann-Whitney (pairwise) tests. The test used and error bars are defined in each Figure legend.

## Acknowledgments

Research reported in this publication was supported by the National Institute of Diabetes and Digestive and Kidney Diseases of the National Institutes of Health under Award Number R01DK139790. The content is solely the responsibility of the authors and does not necessarily represent the official views of the National Institutes of Health.

## References

1. A. I. Ferrer, J. R. Trinidad, O. Sandiford, J. P. Etchegaray, P. Rameshwar, Epigenetic dynamics in cancer stem cell dormancy. Cancer Metastasis Rev 39, 721–738 (2020).

2. B. C. Prager, Q. Xie, S. Bao, J. N. Rich, Cancer Stem Cells: The Architects of the Tumor Ecosystem. Cell Stem Cell 24, 41–53 (2019).

3. A. Ferrer et al., Hypoxia-mediated changes in bone marrow microenvironment in breast cancer dormancy. Cancer Lett 488, 9–17 (2020).

4. X. Hu et al., Induction of cancer cell stemness by chemotherapy. Cell Cycle 11, 2691–2698 (2012).

5. H. Lu et al., Chemotherapy-Induced Ca(2+) Release Stimulates Breast Cancer Stem Cell Enrichment. Cell Rep 18, 1946–1957 (2017).

6. L. Liu et al., Chemotherapy Induces Breast Cancer Stemness in Association with Dysregulated Monocytosis. Clin Cancer Res 24, 2370–2382 (2018).

7. Y. Zhao et al., Chemotherapy exacerbates ovarian cancer cell migration and cancer stem cell-like characteristics through GLI1. Br J Cancer 122, 1638–1648 (2020).

8. H. Liu, L. Lv, K. Yang, Chemotherapy targeting cancer stem cells. Am J Cancer Res 5, 880–893 (2015).

9. I. Ben-Porath et al., An embryonic stem cell-like gene expression signature in poorly differentiated aggressive human tumors. Nat Genet 40, 499–507 (2008).

10. M. L. Suva, N. Riggi, B. E. Bernstein, Epigenetic reprogramming in cancer. Science 339, 1567–1570 (2013).

11. K. Yagita et al., Development of the circadian oscillator during differentiation of mouse embryonic stem cells in vitro. Proc Natl Acad Sci U S A 107, 3846–3851 (2010).

12. F. Agriesti, O. Cela, N. Capitanio, “Time Is out of Joint” in Pluripotent Stem Cells: How and Why. Int J Mol Sci 25 (2024).

13. E. Kowalska, E. Moriggi, C. Bauer, C. Dibner, S. A. Brown, The circadian clock starts ticking at a developmentally early stage. J Biol Rhythms 25, 442–449 (2010).

14. Y. Feng, X. Liu, S. Pauklin, 3D chromatin architecture and epigenetic regulation in cancer stem cells. Protein Cell 12, 440–454 (2021).

15. D. Friedmann-Morvinski, I. M. Verma, Dedifferentiation and reprogramming: origins of cancer stem cells. EMBO Rep 15, 244–253 (2014).

16. K. Guiro, S. A. Patel, S. J. Greco, P. Rameshwar, T. L. Arinzeh, Investigating breast cancer cell behavior using tissue engineering scaffolds. PLoS One 10, e0118724 (2015).

17. E. Nolan, G. J. Lindeman, J. E. Visvader, Deciphering breast cancer: from biology to the clinic. Cell 10.1016/j.cell.2023.01.040 (2023).

18. A. Wiechert et al., Cisplatin induces stemness in ovarian cancer. Oncotarget 7, 30511–30522 (2016).

19. D. Lee, H. S. Jeong, S. Y. Hwang, Y. G. Lee, Y. J. Kang, ABCB1 confers resistance to carboplatin by accumulating stem-like cells in the G2/M phase of the cell cycle in p53(null) ovarian cancer. Cell Death Discov 11, 132 (2025).

20. W. A. Abreu de Oliveira et al., Wnt/beta-Catenin Inhibition Disrupts Carboplatin Resistance in Isogenic Models of Triple-Negative Breast Cancer. Front Oncol 11, 705384 (2021).

21. D. R. Pattabiraman, R. A. Weinberg, Tackling the cancer stem cells - what challenges do they pose? Nat Rev Drug Discov 13, 497–512 (2014).

22. J. Chen et al., A restricted cell population propagates glioblastoma growth after chemotherapy. Nature 488, 522–526 (2012).

23. F. Rijo-Ferreira, J. S. Takahashi, Genomics of circadian rhythms in health and disease. Genome Med 11, 82 (2019).

24. R. Zhang, N. F. Lahens, H. I. Ballance, M. E. Hughes, J. B. Hogenesch, A circadian gene expression atlas in mammals: implications for biology and medicine. Proc Natl Acad Sci U S A 111, 16219–16224 (2014).

25. J. S. Takahashi, Transcriptional architecture of the mammalian circadian clock. Nat Rev Genet 18, 164–179 (2017).

26. J. P. Etchegaray, R. Mostoslavsky, Interplay between Metabolism and Epigenetics: A Nuclear Adaptation to Environmental Changes. Mol Cell 62, 695–711 (2016).

27. K. Straif, The burden of occupational cancer. Occup Environ Med 65, 787–788 (2008).

28. R. G. Stevens et al., Considerations of circadian impact for defining ‘shift work’ in cancer studies: IARC Working Group Report. Occup Environ Med 68, 154–162 (2011).

29. R. M. Lunn et al., Health consequences of electric lighting practices in the modern world: A report on the National Toxicology Program’s workshop on shift work at night, artificial light at night, and circadian disruption. Sci Total Environ 607-608, 1073–1084 (2017).

30. C. Nor et al., Cisplatin induces Bmi-1 and enhances the stem cell fraction in head and neck cancer. Neoplasia 16, 137–146 (2014).

31. X. Chu et al., Cancer stem cells: advances in knowledge and implications for cancer therapy. Signal Transduct Target Ther 9, 170 (2024).

32. M. Kim, L. Bakyt, A. Akhmetkaliyev, D. Toktarkhanova, D. Bulanin, Re-Sensitizing Cancer Stem Cells to Conventional Chemotherapy Agents. Int J Mol Sci 24 (2023).

33. M. Takata et al., Daily expression of mRNAs for the mammalian Clock genes Per2 and clock in mouse suprachiasmatic nuclei and liver and human peripheral blood mononuclear cells. Jpn J Pharmacol 90, 263–269 (2002).

34. M. A. Chico et al., Cancer Stem Cells in Sarcomas: In Vitro Isolation and Role as Prognostic Markers: A Systematic Review. Cancers (Basel) 15 (2023).

35. S. Nallanthighal, J. P. Heiserman, D. J. Cheon, The Role of the Extracellular Matrix in Cancer Stemness. Front Cell Dev Biol 7, 86 (2019).

36. M. Dudek, J. Swift, Q. J. Meng, The circadian clock and extracellular matrix homeostasis in aging and age-related diseases. Am J Physiol Cell Physiol 325, C52–C59 (2023).

37. N. Wang et al., Vascular PPARgamma controls circadian variation in blood pressure and heart rate through Bmal1. Cell Metab 8, 482–491 (2008).

38. P. Charoensuksai, W. Xu, PPARs in Rhythmic Metabolic Regulation and Implications in Health and Disease. PPAR Res 2010 (2010).

39. M. K. Bunger et al., Mop3 is an essential component of the master circadian pacemaker in mammals. Cell 103, 1009–1017 (2000).

40. R. Chen et al., Rhythmic PER abundance defines a critical nodal point for negative feedback within the circadian clock mechanism. Mol Cell 36, 417–430 (2009).

41. L. Fang et al., Circadian Clock Gene CRY2 Degradation Is Involved in Chemoresistance of Colorectal Cancer. Mol Cancer Ther 14, 1476–1487 (2015).

42. Y. Shen et al., NF-kappaB modifies the mammalian circadian clock through interaction with the core clock protein BMAL1. PLoS Genet 17, e1009933 (2021).

43. E. Maury, B. Navez, S. M. Brichard, Circadian clock dysfunction in human omental fat links obesity to metabolic inflammation. Nat Commun 12, 2388 (2021).

44. R. Schneider, R. M. Linka, H. Reinke, HSP90 affects the stability of BMAL1 and circadian gene expression. J Biol Rhythms 29, 87–96 (2014).

45. L. Core, K. Adelman, Promoter-proximal pausing of RNA polymerase II: a nexus of gene regulation. Genes Dev 33, 960–982 (2019).

46. J. P. Etchegaray et al., The Histone Deacetylase SIRT6 Restrains Transcription Elongation via Promoter-Proximal Pausing. Mol Cell 75, 683–699 e687 (2019).

47. Y. Zhang, X. Wang, Targeting the Wnt/beta-catenin signaling pathway in cancer. J Hematol Oncol 13, 165 (2020).

48. N. Takebe, P. J. Harris, R. Q. Warren, S. P. Ivy, Targeting cancer stem cells by inhibiting Wnt, Notch, and Hedgehog pathways. Nat Rev Clin Oncol 8, 97–106 (2011).

49. O. Takase et al., The role of NF-kappaB signaling in the maintenance of pluripotency of human induced pluripotent stem cells. PLoS One 8, e56399 (2013).

50. M. S. Simic et al., Transient activation of the UPR(ER) is an essential step in the acquisition of pluripotency during reprogramming. Sci Adv 5, eaaw0025 (2019).

51. P. Zhang et al., Regulation of induced pluripotent stem (iPS) cell induction by Wnt/beta-catenin signaling. J Biol Chem 289, 9221–9232 (2014).

52. C. A. Dudley et al., Altered patterns of sleep and behavioral adaptability in NPAS2-deficient mice. Science 301, 379–383 (2003).

53. G. Yang et al., Systemic PPARgamma deletion impairs circadian rhythms of behavior and metabolism. PLoS One 7, e38117 (2012).

54. S. De Cosmo, G. Mazzoccoli, Retinoid X Receptors Intersect the Molecular Clockwork in the Regulation of Liver Metabolism. Front Endocrinol (Lausanne) 8, 24 (2017).

55. T. Tamaru et al., Synchronization of circadian Per2 rhythms and HSF1-BMAL1:CLOCK interaction in mouse fibroblasts after short-term heat shock pulse. PLoS One 6, e24521 (2011).

56. C. Lee, J. P. Etchegaray, F. R. Cagampang, A. S. Loudon, S. M. Reppert, Posttranslational mechanisms regulate the mammalian circadian clock. Cell 107, 855–867 (2001).

57. L. Chen, G. Yang, PPARs Integrate the Mammalian Clock and Energy Metabolism. PPAR Res 2014, 653017 (2014).

58. H. Kawasaki, R. Doi, K. Ito, M. Shimoda, N. Ishida, The circadian binding of CLOCK protein to the promoter of C/ebpalpha gene in mouse cells. PLoS One 8, e58221 (2013).

59. N. Koike et al., Transcriptional architecture and chromatin landscape of the core circadian clock in mammals. Science 338, 349–354 (2012).

60. M. P. Antoch et al., Functional identification of the mouse circadian Clock gene by transgenic BAC rescue. Cell 89, 655–667 (1997).

61. W. Zou et al., Association of CD44 and CD24 phenotype with lymph node metastasis and survival in triple-negative breast cancer. Int J Clin Exp Pathol 13, 1008–1016 (2020).

62. S. Ghuwalewala et al., CD44(high)CD24(low) molecular signature determines the Cancer Stem Cell and EMT phenotype in Oral Squamous Cell Carcinoma. Stem Cell Res 16, 405–417 (2016).

63. J. L. Huang, M. Oshi, I. Endo, K. Takabe, Clinical relevance of stem cell surface markers CD133, CD24, and CD44 in colorectal cancer. Am J Cancer Res 11, 5141–5154 (2021).

64. A. Z. Ayob, T. S. Ramasamy, Cancer stem cells as key drivers of tumour progression. J Biomed Sci 25, 20 (2018).

65. C. R. Justus, M. A. Marie, E. J. Sanderlin, L. V. Yang, Transwell In Vitro Cell Migration and Invasion Assays. Methods Mol Biol 2644, 349–359 (2023).

66. Y. Okamoto-Uchida et al., Post-translational Modifications are Required for Circadian Clock Regulation in Vertebrates. Curr Genomics 20, 332–339 (2019).

67. Y. Xi, D. Chen, Physiology. Partitioning the circadian clock. Science 345, 1122–1123 (2014).

68. N. Khaled, Y. Bidet, New Insights into the Implication of Epigenetic Alterations in the EMT of Triple Negative Breast Cancer. Cancers (Basel) 11 (2019).

69. A. Paul, M. Danley, B. Saha, O. Tawfik, S. Paul, PKCzeta Promotes Breast Cancer Invasion by Regulating Expression of E-cadherin and Zonula Occludens-1 (ZO-1) via NFkappaB-p65. Sci Rep 5, 12520 (2015).

70. S. Li et al., Epigenetic regulation of LINC01270 in breast cancer progression by mediating LAMA2 promoter methylation and MAPK signaling pathway. Cell Biol Toxicol 39, 1359–1375 (2023).

71. H. Yoshitane et al., Functional D-box sequences reset the circadian clock and drive mRNA rhythms. Commun Biol 2, 300 (2019).

72. Y. Larriba, I. C. Mason, R. Saxena, F. Scheer, C. Rueda, CIRCUST: A novel methodology for temporal order reconstruction of molecular rhythms; validation and application towards a daily rhythm gene expression atlas in humans. PLoS Comput Biol 19, e1011510 (2023).

73. S. I. Choi et al., HSPA1L Enhances Cancer Stem Cell-Like Properties by Activating IGF1Rbeta and Regulating beta-Catenin Transcription. Int J Mol Sci 21 (2020).

74. O. A. Sandiford et al., Mesenchymal Stem Cell-Secreted Extracellular Vesicles Instruct Stepwise Dedifferentiation of Breast Cancer Cells into Dormancy at the Bone Marrow Perivascular Region. Cancer Res 81, 1567–1582 (2021).

75. S. Mukherjee et al., Breast cancer stem cells generate immune-suppressive T regulatory cells by secreting TGFbeta to evade immune-elimination. Discov Oncol 14, 220 (2023).

76. Y. Duchartre, Y. M. Kim, M. Kahn, The Wnt signaling pathway in cancer. Crit Rev Oncol Hematol 99, 141–149 (2016).

77. G. Hassan, M. Seno, ERBB Signaling Pathway in Cancer Stem Cells. Adv Exp Med Biol 1393, 65–81 (2022).

78. A. L. Rinkenbaugh, A. S. Baldwin, The NF-kappaB Pathway and Cancer Stem Cells. Cells 5 (2016).

79. D. Filipponi, M. Pagnuzzi-Boncompagni, G. Pages, Inhibiting ALK2/ALK3 Signaling to Differentiate and Chemo-Sensitize Medulloblastoma. Cancers (Basel) 14 (2022).

80. S. Diep, M. Maddukuri, S. Yamauchi, G. Geshow, N. A. Delk, Interleukin-1 and Nuclear Factor Kappa B Signaling Promote Breast Cancer Progression and Treatment Resistance. Cells 11 (2022).

81. E. S. Rasmussen, J. S. Takahashi, C. B. Green, Time to target the circadian clock for drug discovery. Trends Biochem Sci 47, 745–758 (2022).

82. Y. Liu, A. Sancar, Biochemical mechanism of the mammalian circadian clock. FEBS Lett 600, 716–731 (2026).

83. H. Tatetsu et al., SALL4, the missing link between stem cells, development and cancer. Gene 584, 111–119 (2016).

84. K. Urbonas et al., Overexpression of EMT-related transcription factors SNAI1 and ZEB1 is associated with more aggressive clinicopathological features of pancreatic cancer. PLoS One 21, e0339964 (2026).

85. B. Wang, C. W. Lee, A. Witt, A. Thakkar, T. A. Ince, Heat shock factor 1 induces cancer stem cell phenotype in breast cancer cell lines. Breast Cancer Res Treat 153, 57–66 (2015).

86. Z. Dong et al., Targeting Glioblastoma Stem Cells through Disruption of the Circadian Clock. Cancer Discov 9, 1556–1573 (2019).

87. R. V. Puram et al., Core Circadian Clock Genes Regulate Leukemia Stem Cells in AML. Cell 165, 303–316 (2016).

88. P. Godwin et al., Targeting nuclear factor-kappa B to overcome resistance to chemotherapy. Front Oncol 3, 120 (2013).

89. B. M. Fortin, A. L. Mahieu, R. C. Fellows, N. R. Pannunzio, S. Masri, Circadian clocks in health and disease: Dissecting the roles of the biological pacemaker in cancer. F1000Res 12, 116 (2023).

90. R. L. B. Costa, W. J. Gradishar, Triple-Negative Breast Cancer: Current Practice and Future Directions. J Oncol Pract 13, 301–303 (2017).

91. J. Yu et al., Induced pluripotent stem cell lines derived from human somatic cells. Science 318, 1917–1920 (2007).

92. A. Balsalobre et al., Resetting of circadian time in peripheral tissues by glucocorticoid signaling. Science 289, 2344–2347 (2000).

93. M. A. Livshits et al., Isolation of exosomes by differential centrifugation: Theoretical analysis of a commonly used protocol. Sci Rep 5, 17319 (2015).

94. L. Gerrard, D. Zhao, A. J. Clark, W. Cui, Stably transfected human embryonic stem cell clones express OCT4-specific green fluorescent protein and maintain self-renewal and pluripotency. Stem Cells 23, 124–133 (2005).

95. L. Mei et al., Long-term in vivo recording of circadian rhythms in brains of freely moving mice. Proc Natl Acad Sci U S A 115, 4276–4281 (2018).

96. N. Huang et al., Crystal structure of the heterodimeric CLOCK:BMAL1 transcriptional activator complex. Science 337, 189–194 (2012).

97. R. Ye, C. P. Selby, N. Ozturk, Y. Annayev, A. Sancar, Biochemical analysis of the canonical model for the mammalian circadian clock. J Biol Chem 286, 25891–25902 (2011).

98. S. Ozgur, A. Sancar, Purification and properties of human blue-light photoreceptor cryptochrome 2. Biochemistry 42, 2926–2932 (2003).

99. L. Warren et al., Highly efficient reprogramming to pluripotency and directed differentiation of human cells with synthetic modified mRNA. Cell Stem Cell 7, 618–630 (2010).

100. B. K. Chou et al., Efficient human iPS cell derivation by a non-integrating plasmid from blood cells with unique epigenetic and gene expression signatures. Cell Res 21, 518–529 (2011).

101. S. Yamazaki, J. S. Takahashi, Real-time luminescence reporting of circadian gene expression in mammals. Methods Enzymol 393, 288–301 (2005).

102. P. Indic, E. N. Brown, Characterizing the amplitude dynamics of the human core-temperature circadian rhythm using a stochastic-dynamic model. J Theor Biol 239, 499–506 (2006).

103. G. Cornelissen, Cosinor-based rhythmometry. Theor Biol Med Model 11, 16 (2014).

104. P. A. Ewels et al., The nf-core framework for community-curated bioinformatics pipelines. Nat Biotechnol 38, 276–278 (2020).

105. Y. Fan et al., LRR1-mediated replisome disassembly promotes DNA replication by recycling replisome components. J Cell Biol 220 (2021).

106. G. Watts, E. Wilkinson, What the NHS is learning from the British army in the covid-19 crisis. BMJ 369, m2055 (2020).

107. A. Colaprico et al., TCGAbiolinks: an R/Bioconductor package for integrative analysis of TCGA data. Nucleic Acids Res 44, e71 (2016).

108. M. Mounir et al., New functionalities in the TCGAbiolinks package for the study and integration of cancer data from GDC and GTEx. PLoS Comput Biol 15, e1006701 (2019).

109. T. C. Silva et al., TCGA Workflow: Analyze cancer genomics and epigenomics data using Bioconductor packages. F1000Res 5, 1542 (2016).

110. K. A. Reimer, C. A. Mimoso, K. Adelman, K. M. Neugebauer, Co-transcriptional splicing regulates 3’ end cleavage during mammalian erythropoiesis. Mol Cell 81, 998–1012 e1017 (2021).

111. G. L. Zhang, Y. Zhang, K. X. Cao, X. M. Wang, Orthotopic Injection of Breast Cancer Cells into the Mice Mammary Fat Pad. J Vis Exp 10.3791/58604 (2019).

