## Supplemental Figures and Legends for "Pluripotency and Transcriptional Pausing Disrupt Circadian Rhythms to Facilitate the Enrichment of Cancer Stem Cells"

### Supplemental Figure S1

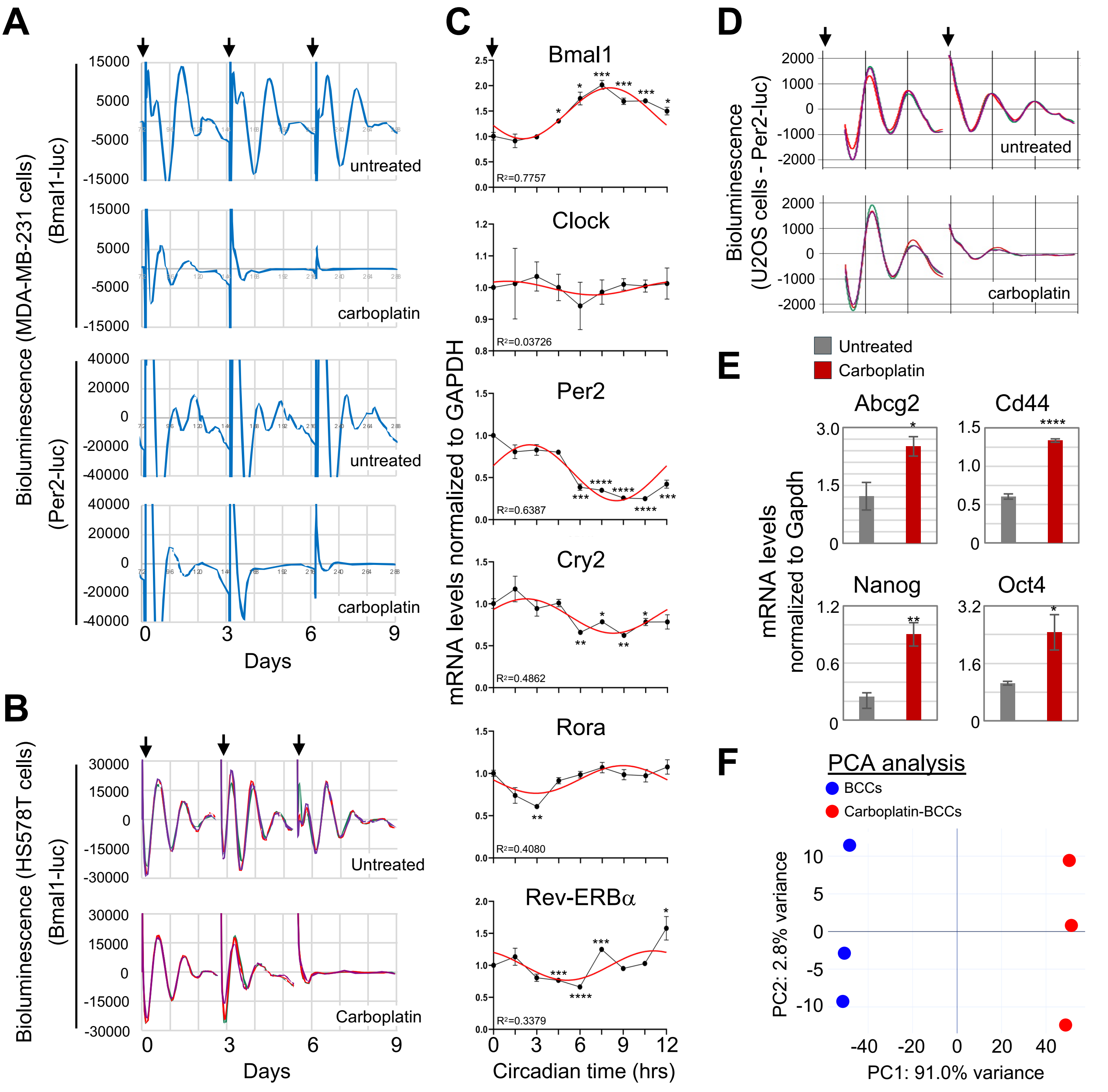

**Supplemental Figure 1. Verification of Circadian Rhythms and Their Loss Mediated by Carboplatin Treatment in Different Cancer Cell Lines.**
(**A**) Bioluminescent circadian profiles of MDA-MB-231 with either Bmal1-luc or Per2-luc circadian reporter with or without carboplatin (50ug/ml). Arrows indicate resynchronization points every 3 days. Bioluminescent plots show an average of 3 independent circadian profiles from each cell line. (**B**) Bioluminescent circadian profiles of another Triple-negative breast cancer cell line and HS578T treated with carboplatin (50ug/ml). Arrows indicate resynchronization points every 3 days. Bioluminescent plots show 3 independent circadian profiles from each cell line. (**C**) Quantification of RT-qPCR showing the normalized gene expression of circadian genes every 3 hours during a 24-hour time-course of MDA-MB-231, normalized to GAPDH. Rhythmicity of gene expression was assessed by nonlinear regression using the Prism sine wave with nonzero baseline model . Error bars represent SEM (n=4), pair-end t-test compared to timepoint 0. (*p<0.05, **p<0.01, ***p<0.001, ****p<0.0001). (**D**) Bioluminescent circadian profile of untreated or carboplatin treated U2OS cells with Per2-Luc reporter. Profiles show 3 independent circadian profiles from each treatment. (**E**) RT-qPCR quantification of CSCs related genes in U2OS cells treated with 50 ug/mL carboplatin for 5 days normalized to GAPDH. Error bars represent SEM, (n=4), Pair-end t-test (*p<0.05, **p<0.01, ****p<0.0001). (**F**) Principal component analysis (PCA) from RNA-seq in MDA-MB-231 and 5-day carboplatin treated MDA-MB-231 (n=3).

### Supplemental Figure S2

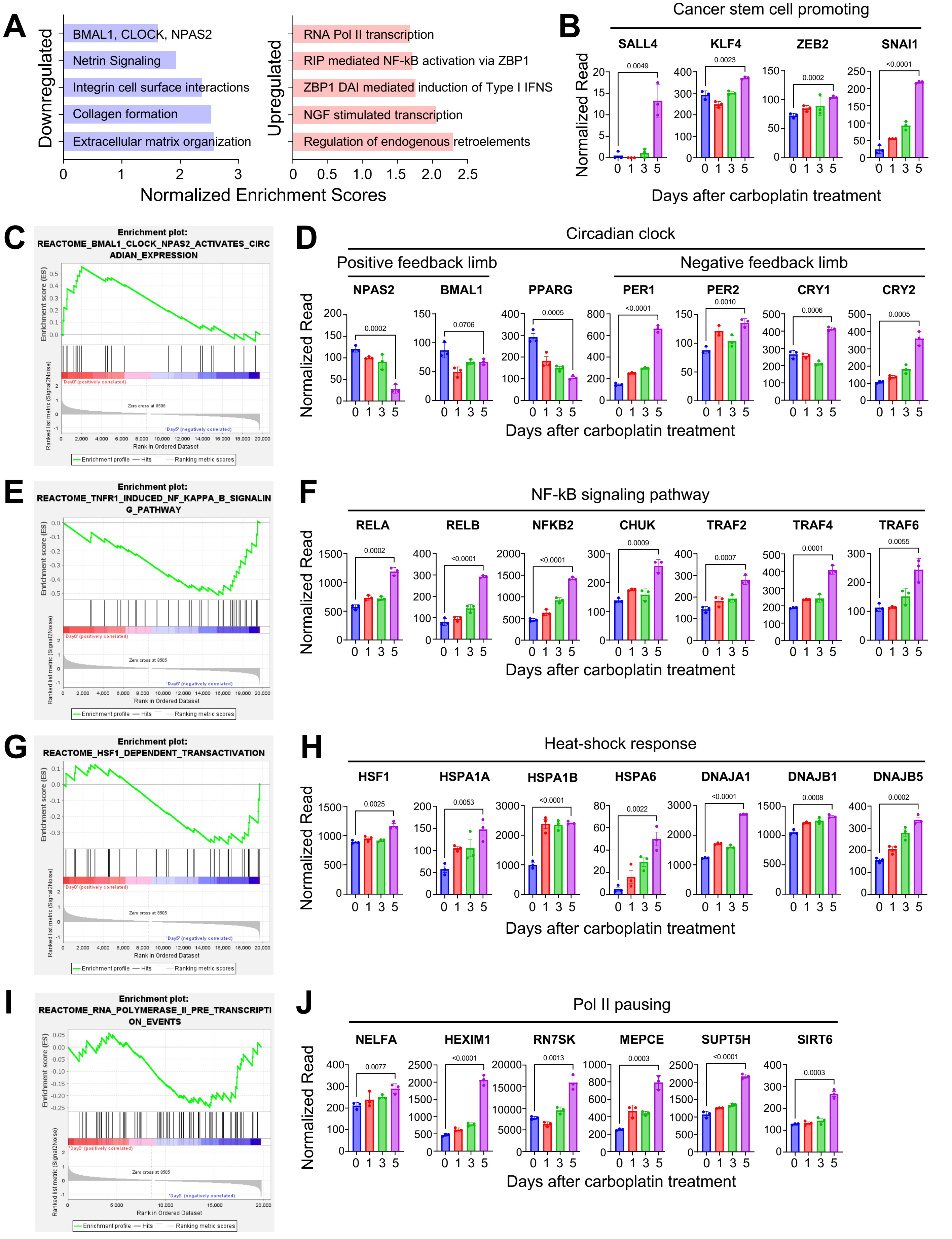

**Supplemental Figure 2. Carboplatin Treatment Induces Gene Expression Changes associated with Cancer Stem Cell Like Progression in MDA-MB-231 Cells.**

(**A**) GSEA analysis on differentially expressed Reactome pathways in MDA-MB-231 before and after carboplatin treatment. (**B**) Bar graph with normalized reads counts of genes associated with cancer stem cells transformation during Carboplatin treatment in MDA-MB-231 cells. (**C**) GSEA plot for circadian pathway of carboplatin treated MDA-MB-231 cells. (**D**) Normalized read counts of Bmal1 dimer components and genes associated with the Bmal1 dimer repression during carboplatin treatment in MDA-MB-231 cells. (**E**) GSEA plot for NFκB signaling pathway in carboplatin treated MDA-MB-231. (**F**) Normalized read counts of genes associated with NFκB signaling during carboplatin treatment in MDA-MB-231 cells. (**G**) GSEA plot for HSF1 dependent transactivation pathway in carboplatin treated MDA-MB-231 cells. (**H**) Normalized read counts of genes associated with heat-shock response pathway. (**I**) GSEA plot for Polymerase II (Pol II) transcription initiation in carboplatin treated MDA-MB-231 cells. (**J**) Normalized read counts of genes associated with Pol II pausing.

### Supplemental Figure S3

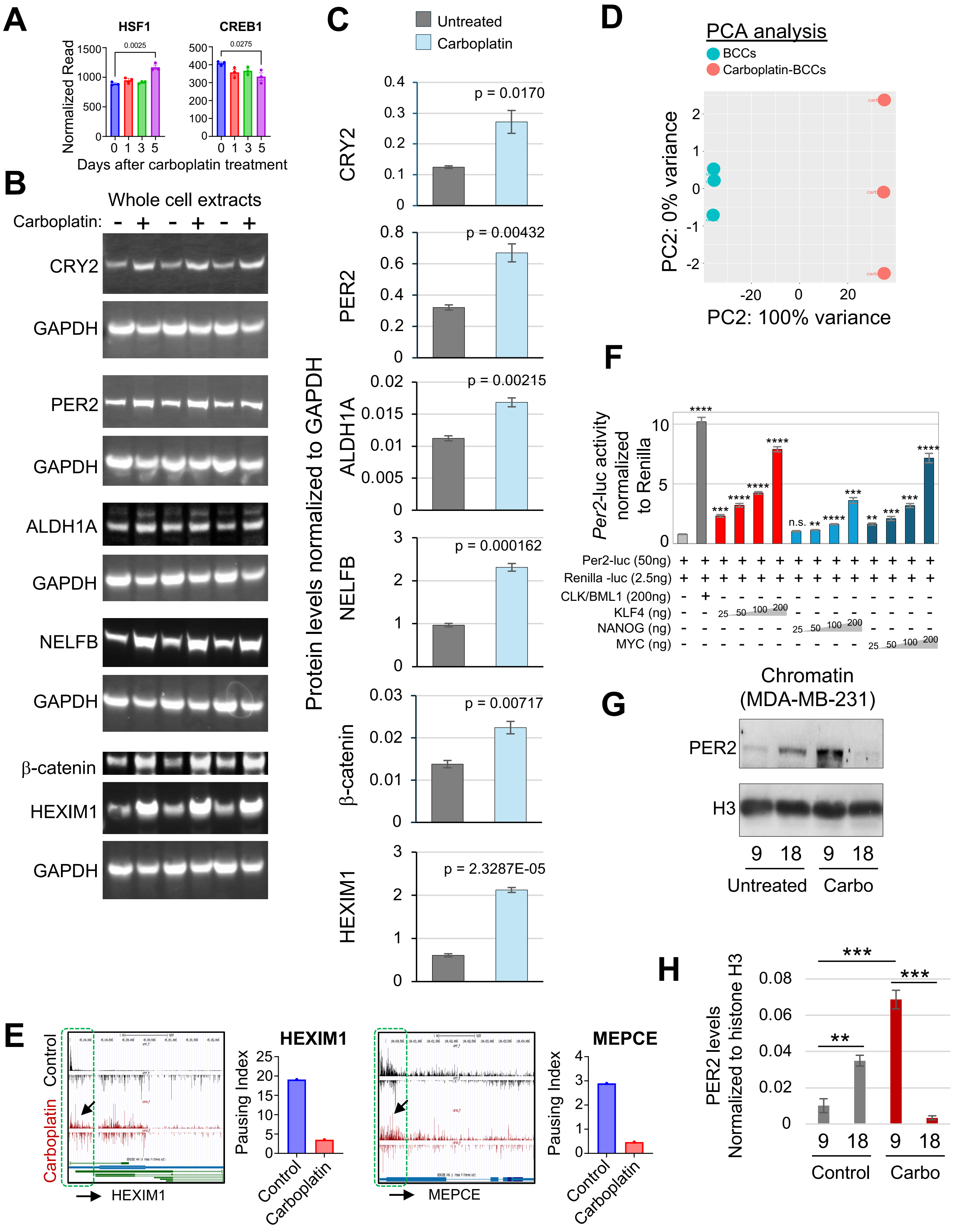

**Supplemental Figure 3**. **Validations of Carboplatin Induced Gene Expression Changes.**

(**A**) Normalized read counts of genes involved in Bmal1-independent activation of PER and CRY genes during carboplatin treatment in MDA-MB-231 cells. (**B**) Western blots for Cry2, Per2, B-catenin, Aldh1a, Nelf-B, Hexim1 and GAPDH (loading control) using whole cell extracts from MDA-MB-231 before and after carboplatin treatment. (**C**) Quantification from Fig. S3B after normalization to GAPDH. Error bars represent SEM, (n=3), pair-end T-test (*p<0.05, **p<0.01, ***p<0.001, *****p<0.00001). (**D**) Principal component analysis (PCA) of PRO-seq data from untreated and 5-day carboplatin treated MDA-MB-231 (n=3).

(**E**) UCSC genome browser images from MDA-MB-231 and 5 day carboplatin treated MDA-MB-231 showing genes involved in transcriptional pausing and their pausing index. (**F**) Dual luciferase assay in HEK293T cells showing the effect of core pluripotency factors KLF4, NANOG, and MYC on circadian gene reporter (Per2-Luc). Luciferase was normalized to Renilla. Error bars represent SEM, (n=4), pair-end-t-test (*p<0.01,**p<0.001,***p<0.0001,****p<0.00001). (**G**) Western blot for BMAL1 and Histone H3 (loading control) using chromatin extracts from HS578T before and after 50ug/ml carboplatin after 5 days synchronized at timepoints 9 and 18 hours. (**H**) Quantification from Fig S3G after normalization to Histone H3. Error bars represent SEM, (n=15), Pair-end t-test (**p<0.01).

### Supplemental Figure S4

A

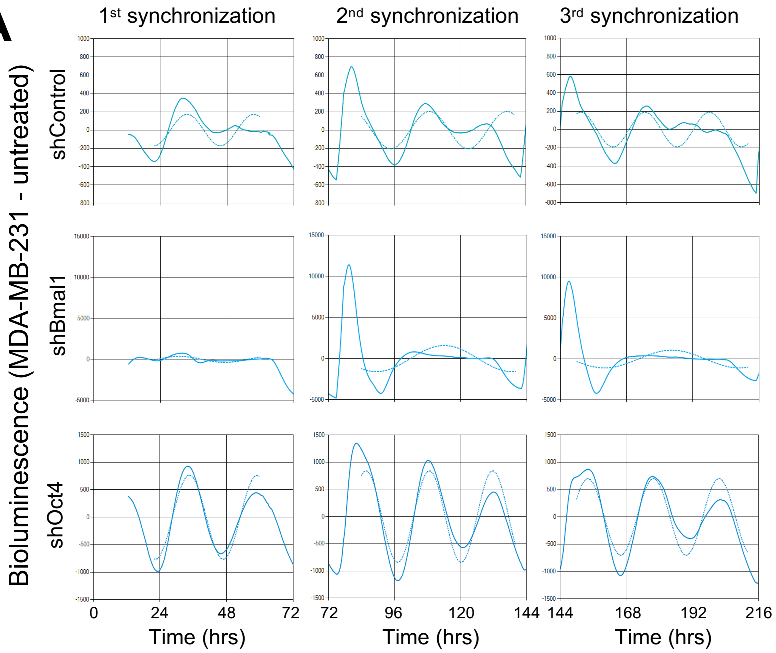


B

shControl

| Sync | Period Calculation | shCtrl | Period | Amplitude |
|---|---|---|---|---|
| 1 | 22-60 | 0-72 | 23.95 | 173 |
| 2 | 84-140 | 72-144 | 28.24 | 204 |
| 3 | 150-212 | 144-216 | 23.25 | 190 |

shBmal1

| Sync | Period Calculation | shBmal1 | Period | Amplitude |
|---|---|---|---|---|
| 1 | 22-60 | 0-72 | 32.64 | 366 |
| 2 | 84-140 | 72-144 | 49.15 | 1,589 |
| 3 | 150-212 | 144-216 | 48.87 | 1,085 |

shOct4

| Sync | Period Calculation | shOct4 | Period | Amplitude |
|---|---|---|---|---|
| 1 | 22-60 | 0-72 | 24.4 | 767 |
| 2 | 84-140 | 72-144 | 23.07 | 844 |
| 3 | 150-212 | 144-216 | 23.81 | 702 |

C

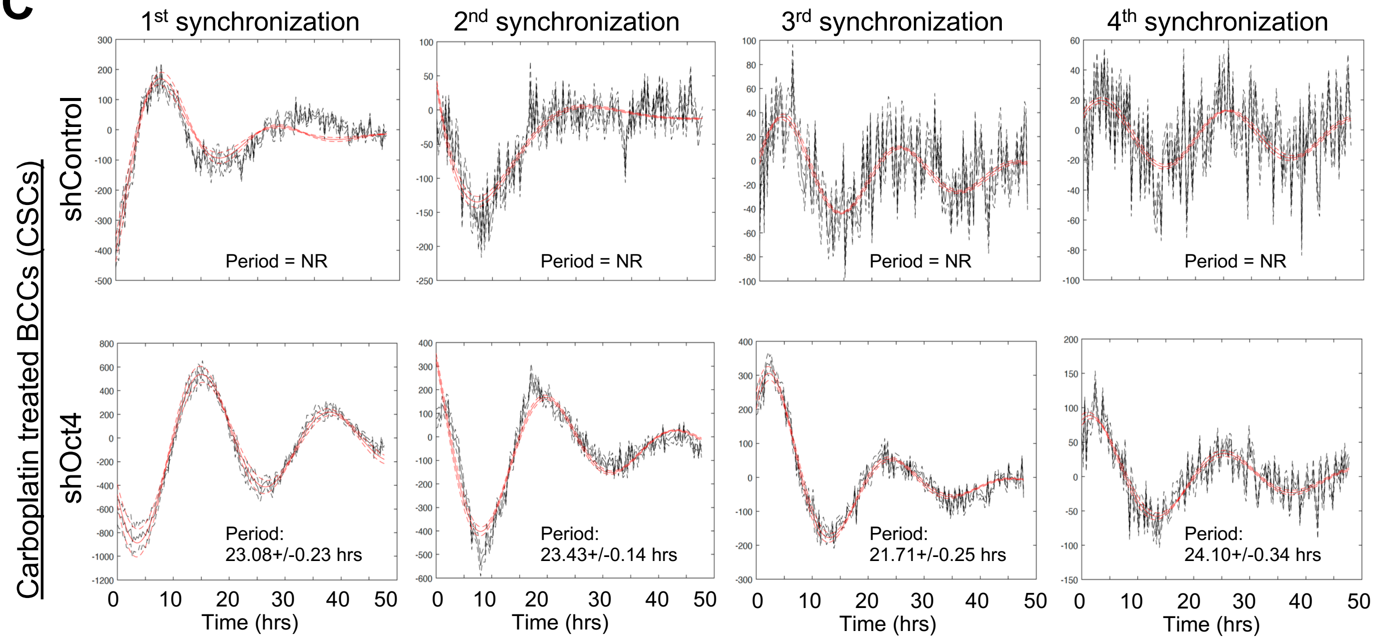

**Supplemental Figure 4**. **Validation of Circadian Rhytmicity in shControl, shBMAL1, and shOCT4.**

(**A**) Bioluminescent profiles derived from KronoAnalyzer software for each synchronization of MDA-MB-231 knocked down for *Bmal1* or *Oct4* in comparison to control. Solid line represents the detrended and smoothed average of 3 independent replicates. Dashed line represents fitted Cosinor analysis as set by KronoAnalyzer software (Atto). (**B**) Quantification of period, acrophase, and amplitude for each of the profiles shown in Fig. S4B. calculated by Cosinor analysis as set by KronoAnalyzer software. Period calculation parameters exclude synchronization peak. (**C**) Evaluation of circadian periodicities in carboplatin treated MDA-MB-231 knockdown for *Oct4* and compared to MDA-MB-231 shControl. Bioluminescent plots show 3 independent circadian profiles from each treatment.

### Supplemental Figure S5

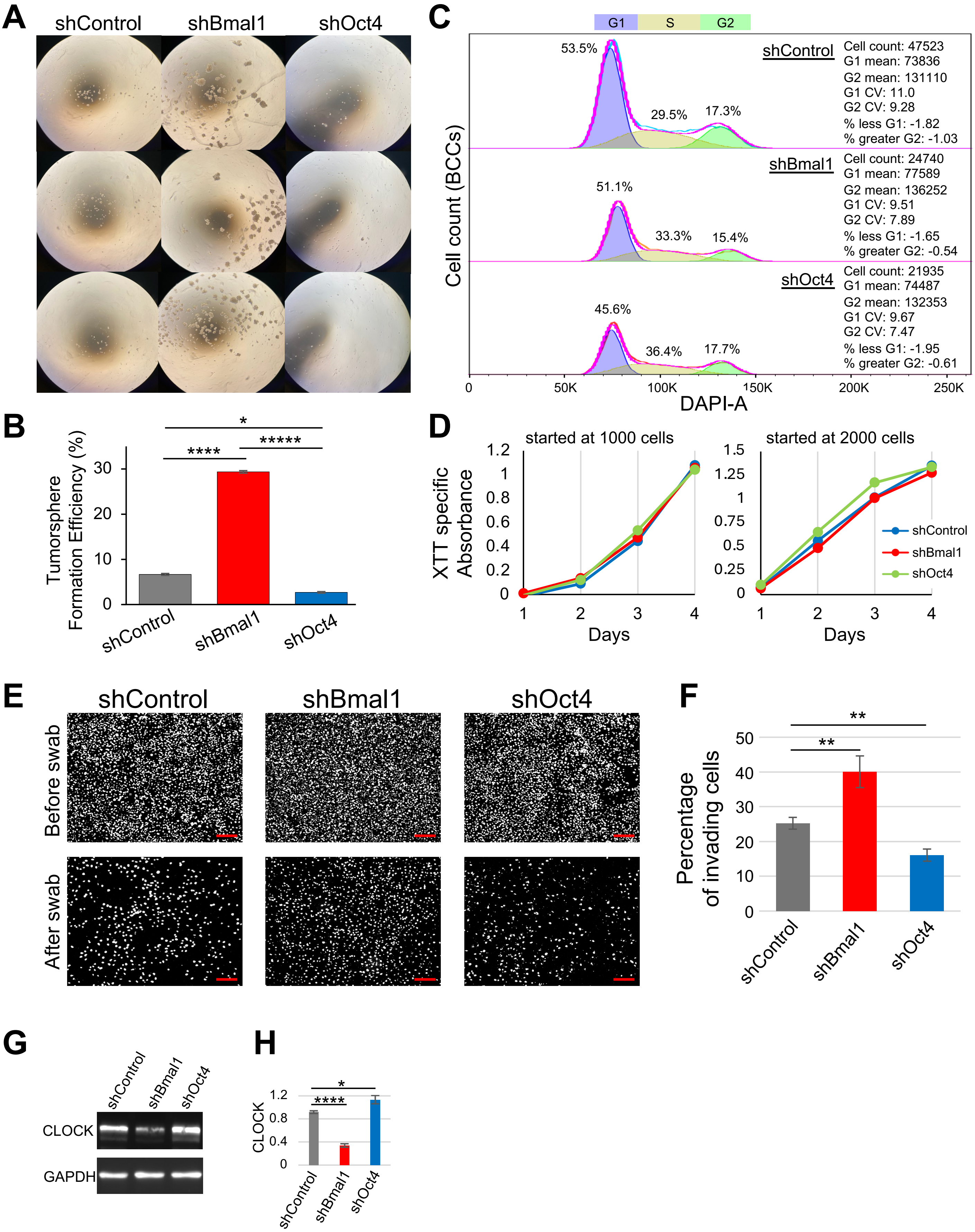

**Supplemental Figure 5. Functional Characterization of shControl, shBMAL1 and shOCT4 MDA-MB-231 cells.**

(**A**) Images of tumorspheres obtained from shControl, shBmal1 and shOct4 MDA-MB-231 during tumorsphere formation efficiency (TFE) assay after 12 days of incubation. (**B**) Quantification of the TFE from Figure S5A. Error bars represent SEM, (n=3), pair-end t-test, (***p<0.001, ****p<0.00001, *****p<0.00001). (**C**) Flow cytometry-based Cell cycle profiling for MDA-MB-231 knockdown for *Oct4* and *Bmal1* in comparison to control. (**D**) XTT assay quantification on shCtrl-, shBmal1-, and shOct4-MDA-MB-231 with two different seeding densities at 1000 and 2000 cells. (**E**) Invasion assay images of cells stained with Hoechest from shControl, shBmal1 and shOct4 before and after swabbing (Scale bar = 200υm). (**F**) Quantification of the invasion assay from Figure S5E. Error bars represent SEM, (n=8), Pair-end t-test ((**p<0.01). (**G)** Western blot for CLOCK and GAPDH (loading control) using whole cell extracts from Control, *Oct4* and *Bmal1* deficient MDA-MB-231. (**H**) Quantification from Fig. S5G after normalization to GAPDH. Error bars represent SEM, (n=3), pair-end T-test (*p<0.05, ****p<0.0001).

### Supplemental Figure S6

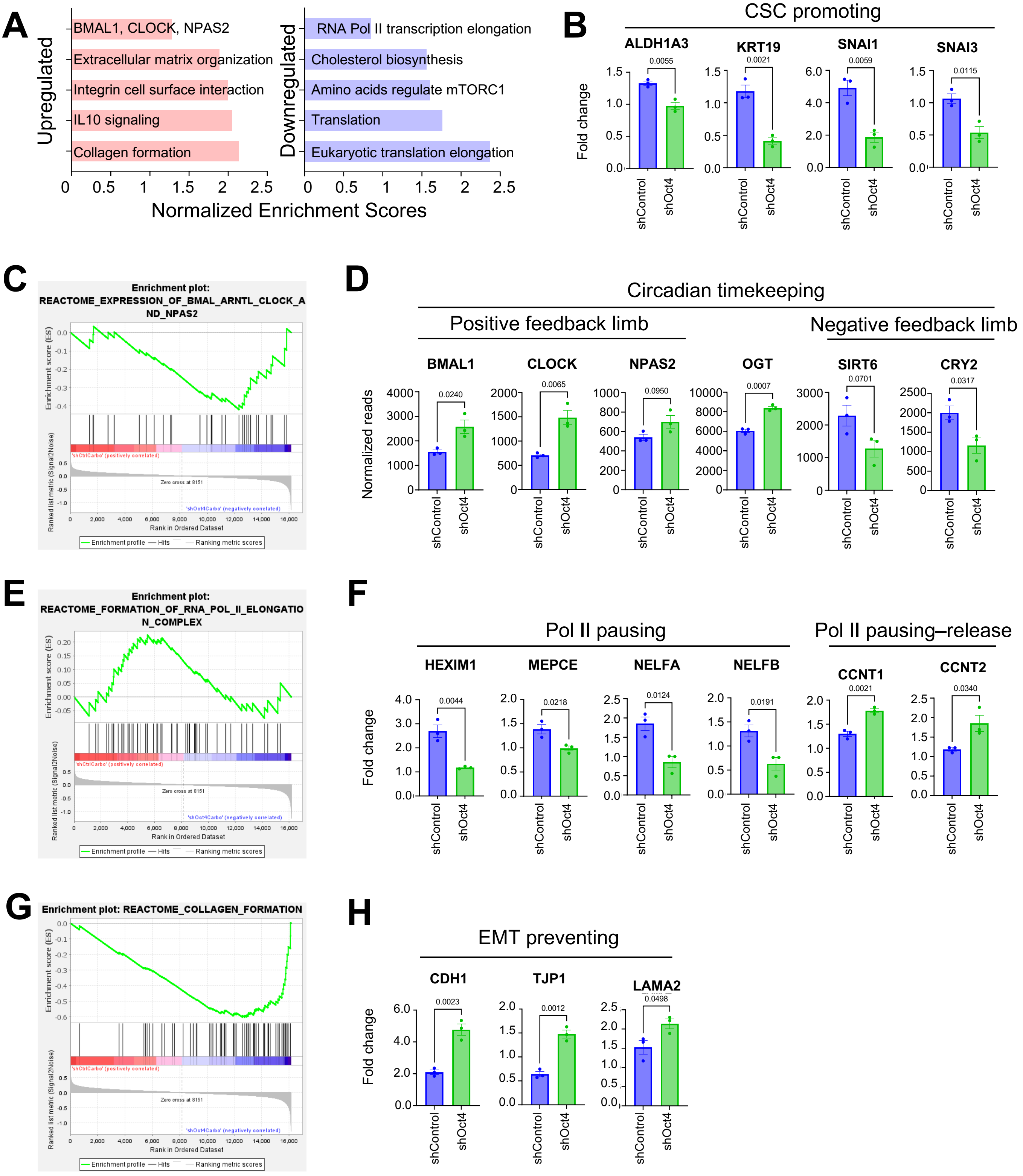

**Supplemental Figure 6. Gene Expression Changes Between shControl and shOCT4 MDA-MB-231 cells.**

(**A**) GSEA analysis on differentially expressed Reactome pathways in MDA-MB-231 cells deficient in *Oct4* and control following 5-day carboplatin treatment. (**B**) Fold Change in normalized read counts before and after carboplatin treatment for genes associated with cancer stem cell promotion in *Oct4* deficient and control MDA-MB-231 cells. (**C**) GSEA plot shows enhanced circadian rhythm pathway in *Oct4* deficient MDA-MB-231 cells post carboplatin treatment. (**D**) Normalized read counts of genes associated with Bmal1 dimer component, repressor and modifier in *Oct4* deficient and control MDA-MB-231 following 5-day carboplatin treatment. (**E**) GSEA plot for Polymerase II (Pol II) transcription elongation of *Oct4* deficient MDA-MB-231. (**F**) Fold change in normalized read counts of genes associated with Pol II pausing and transcription elongation in *Oct4* deficient and control MDA-MB-231 cells. (**G**) GSEA plot for increase enrichment in collagen formation pathway in *Oct4* deficient MDA-MB-231 cells. (**H**) Fold change in normalized read counts of genes associated with suppression of epithelial mesenchymal transition (EMT) in *Oct4* deficient and control MDA-MB-231 cells.

### Supplemental Figure S7

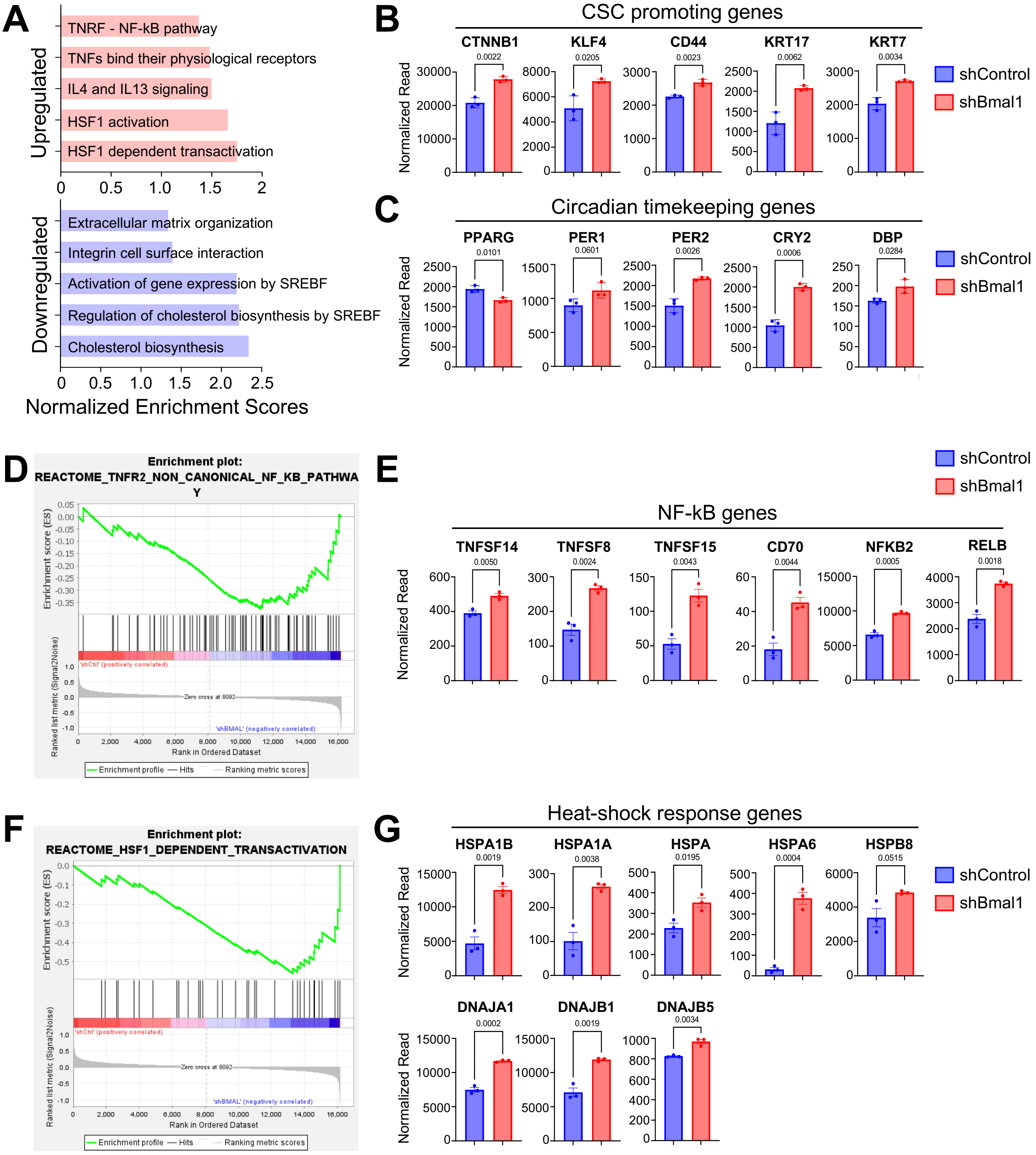

**Supplemental Figure 7. Gene Expression Changes between shControl and shBMAL1 MDA-MB-231 cells.**

(**A**) GSEA analysis of differentially expressed Reactome pathways in *Bmal1* deficient and control MDA-MB-231 cells. (**B**) Normalized read counts of genes associated with cancer stem cell formation in *Bmal1* deficient and control MDA-MB-231 cells. (**C**) Normalized read counts of genes associated with circadian regulation in *Bmal1* deficient and control MDA-MB-231 cells. (**D**) GSEA plot of non-canonical NFκB pathway in *Bmal1* deficient MDA-MB-231 cells. (**E**) Normalized read counts of genes associated with non-canonical NFκB pathway in *Bmal1* deficient and control MDA-MB-231 cells. (**F**)GSEA plot of heat-shock activation pathway of *Bmal1* deficient MDA-MB-231 cells. (**G**) Normalized read counts of genes associated with heat-shock pathway activation in *Bmal1* deficient and control MDA-MB-231 cells.

Supplemental Figure S8

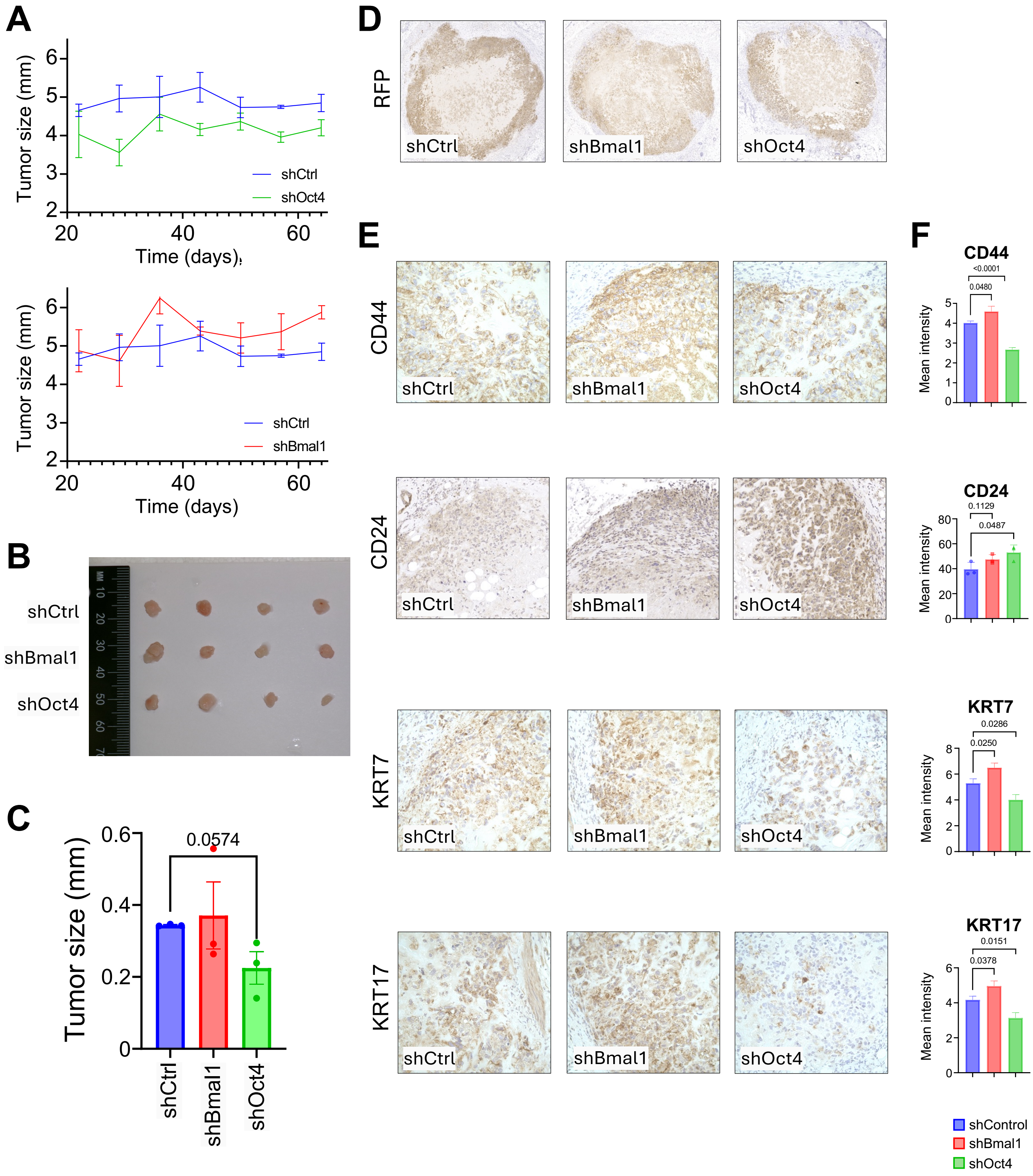

**Supplemental Figure 8. Characterization of Tumors formed by shControl, shBMAL1 and shOCT4 MDA-MB-231 Cells Injected Into Mammary Fat Pad of Mice.**
(**A**) Measurements of orthotopic tumors on severe immunodeficient (SCID) mice injected with *Oct4 deficient, Bmal1* deficient or control MDA-MB-231 cells. (**B**) Representative images of excised tumor from mammary fat pad injection sites. (**C**) Quantification of tumor size from S8B. (**D**) Immunohistochemical (IHC) staining for RFP expression in paraffin-embedded tumor sections. (**E**) IHC staining for CD24/CD44, Ki67, and cytokeratin markers CK7 and CK17 in paraffin-embedded tumors sections. (**F**) Quantification of mean signal intensity from the IHC staining in S8E.

Supplemental Figure S9

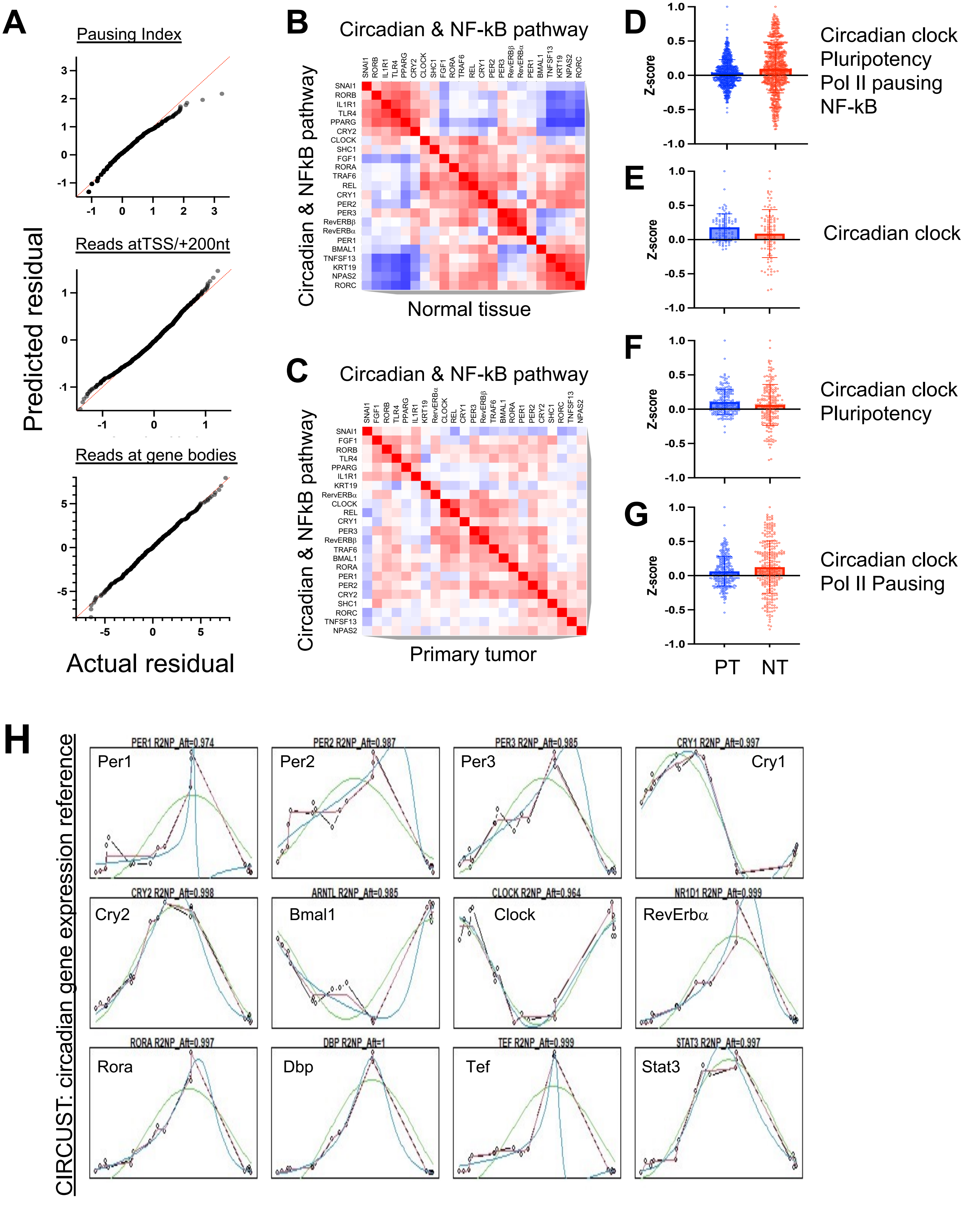

**Supplemental Figure 9. Comparison of Circadian Rhythmicity and Associated Pathways Correlations Between Primary Tumors and Normal Tissue.**

(**A**) QQ plots of PRO-seq analysis showing the normality of the pausing index, assessing the normality of the reads at TSS/+200 nucleotides, and assessing the normality of the reads within gene bodies. (**B-C**) Correlation matrix of Circadian clock and NF-kB pathways in Normal tissue (**B**) and in Primary tumor (**C**) from breast cancer patients in TCGA. (**D-G**) Significance of bidirectional correlations measured by t-test after Fisher Z transform in primary tumor *versus* breast cancer tissues between circadian clock genes, the pluripotency network, transcriptional pausing regulators, and NF-kB pathway ($p$-value$<0.00001$) (**D**), between intrinsic core circadian clock genes ($p$-value$<0.0001$) (**E**), between circadian clock and pluripotency genes ($p$-value$<0.0001$) (**F**), and between circadian clock genes and transcriptional pausing genes ($p$-value$<0.0001$) (**G**) P-value is generated through pair-end T-test. (**H**) CIRCust identifies rhythmic activity in MCF10A samples for 12 circadian genes. Blue, red, and green lines represent predictions from FMM, Cosinor, and modified FMM.

### Supplemental Figure S10

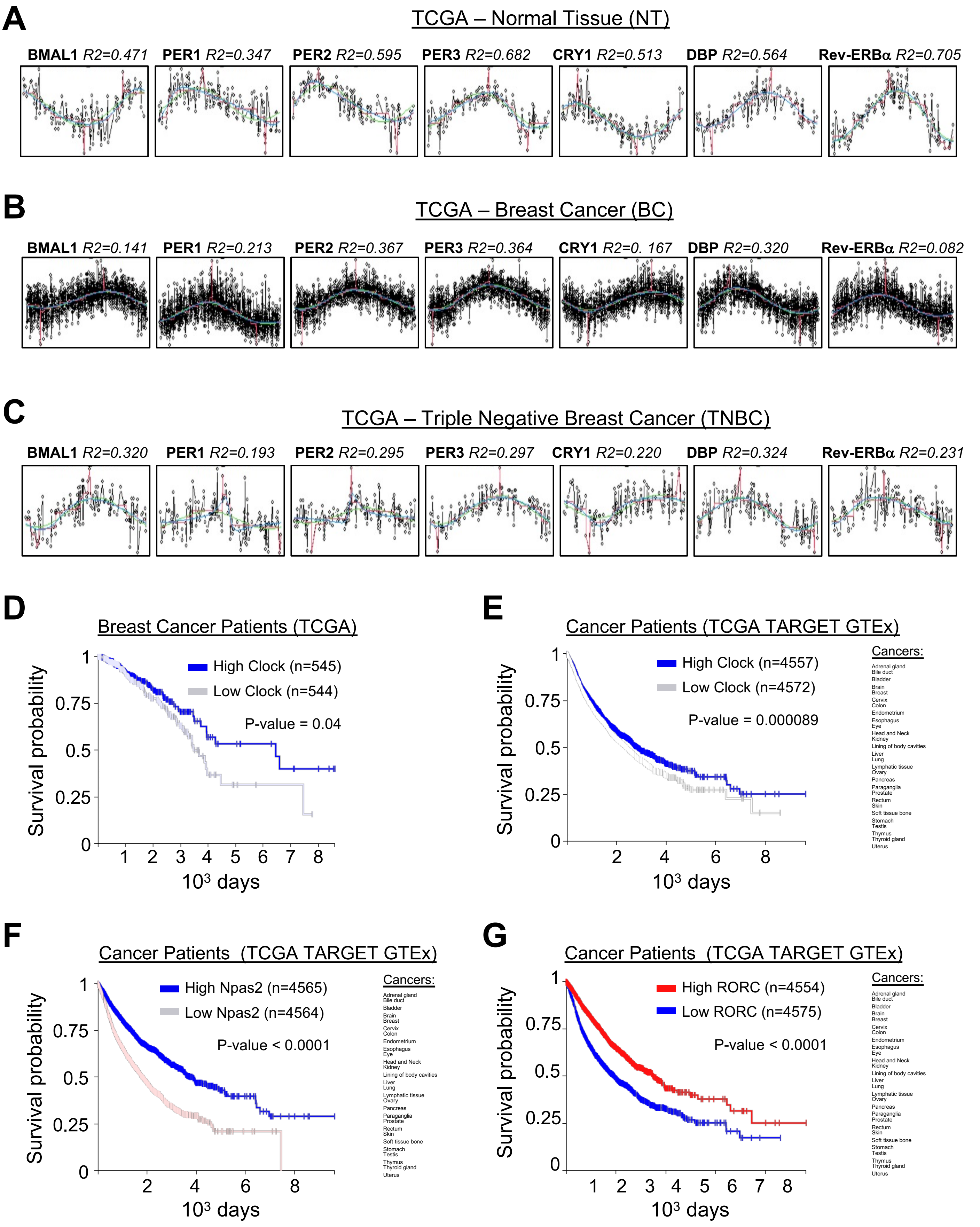

**Supplemental Figure 10. Circadian Robustness in Normal Tissues Compared to Primary Breast Cancer Tumors**

(**A, B, C**) Reordered patient samples based on single circadian gene expression with CIRCust analysis in (**A**) Normal tissue, (**B**) Primary tumor tissue and (**C**) Triple Negative Breast Cancer (TNBC) tissue sample. Blue, red, and green lines represent predictions generated by the FMM, Cosinor, and modified FMM models, respectively. (**D, E**) Kaplan Meier plot from breast cancer patients with high and low levels of *Clock* **(D)** and all cancer patients in TCGA dataset **(E)**. (**F, G**) (Kaplan Meier plot from all cancers patients with low and high Clock (**F**) low and high Npas2 (**G**) low and high RORC in cancer patient in TCGA dataset.

### Supplemental Figure S11

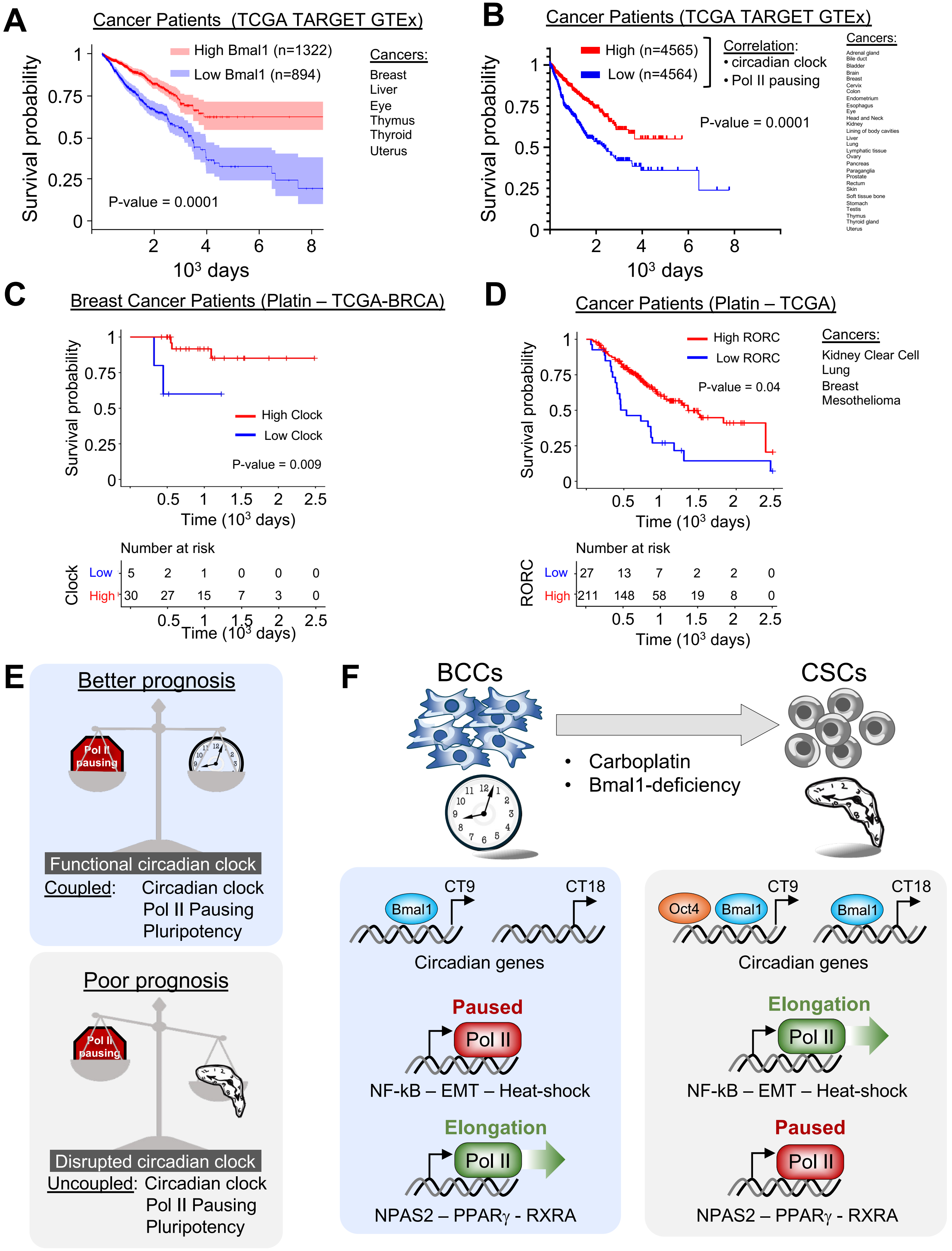

**Supplemental Figure 11. Comparison of Circadian Gene Expression on Patient Outcomes and Proposed Mechanism of Cancer Progression.**
(**A**) Kaplan Meier plot from patients with specific type of cancer from TCGA dataset with low and high Bmal1 expression. (**B**) Kaplan Meier plot for cancer patients in TCGA dataset with low and high gene expression correlation between transcriptional pausing and circadian clock pathway. (**C**) Kaplan Meier survival plot of breast cancer patients treated with platinum-based chemotherapy with high and low expression levels of *Clock gene*. (**D**) Kaplan Meier survival plot of selected cancer types from TCGA dataset treated with platinum-based drugs, stratified by low and high RORC expression levels. (**E**) Schematic illustration depicting an unbalanced relationship between the circadian clock, the pluripotency network and transcriptional pausing in primary tumors from breast cancer patients compared to the balanced interplay between these pathways in normal surrounding tissues. (**F**) Graphical model depicting potential mechanisms underlying the loss of circadian rhythms during formation of CSCs elucidated in this work. Loss of circadian rhythms in CSCs is accomplished by a transcriptional program involving BMAL1 and OCT4 and imbalanced transcriptional pausing causing the upregulation of BMAL1 inhibitors and downregulation of BMAL1 activators.

### Supplemental Figure S12

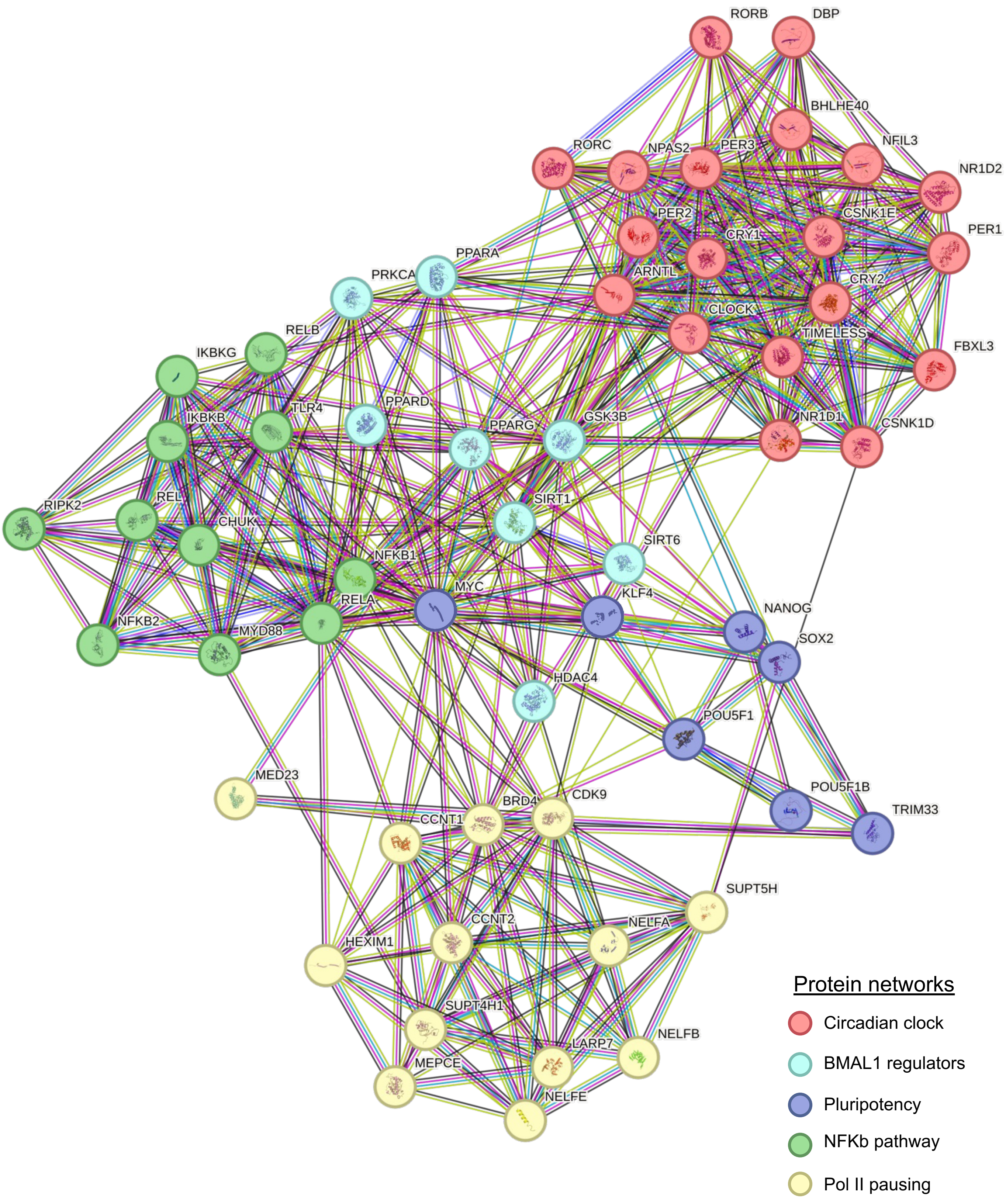

**Supplemental Figure 12.**
**(String Analysis of Genes Involved in Multiple Protein Networks including Circadian Clock, Bmal1 Regulators, Pluripotency, NFkB, and Polymerase II Pausing.**

Supplemental Figure S13

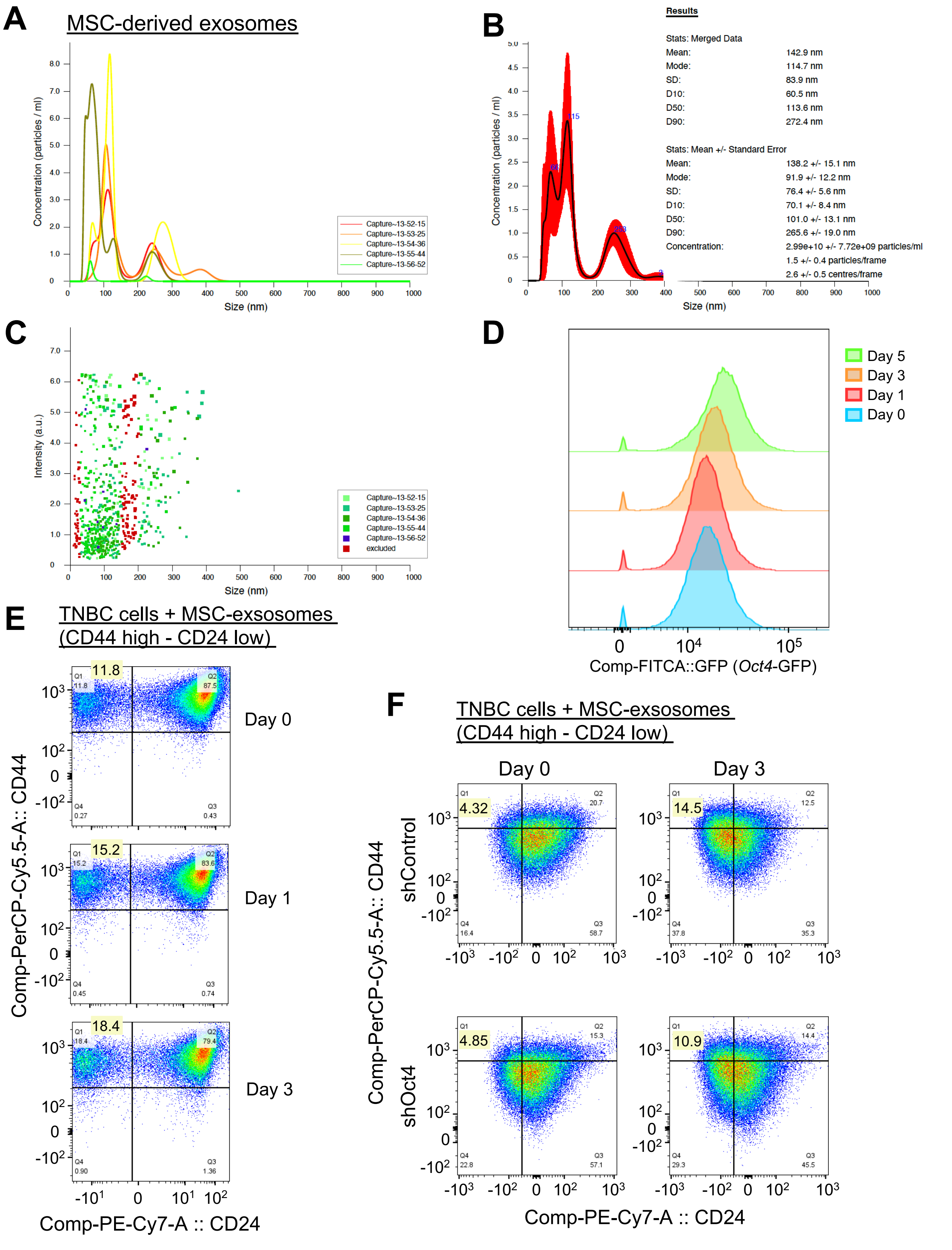

**Supplemental Figure 13. Nanosight Profiles from Purified Exosome Ssolated from MSCs and Their Effect on Cancer Stem Cell like Generation.**

(**A**) Particle concentration and size of MSC derived exosomes quantification of 5 individual nanoparticle tracking captures. (**B**) Averaged particle concentration and size of 5 captures from Fig S1A along with quantitative statistics of MSC derived exosomes. (**C**) Particle size distribution and scattering intensity of MSC derived exosomes quantification of 5 individual nanoparticle tracking captures. (**D**) Flow-cytometry profiling of endogenous Oct4-GFP in MDA-MB-231 treated with $10^8$ MSC derived exosomes for 1, 3, or 5 days. (**E**) Flow cytometry for CD44 and CD24 stained MDA-MB-231 treated with MSC derived exosomes at days 0, 1, and 3. (**F**) Flow cytometry for CD44 and CD24 stained MDA-MB-231 knockdown for *Oct4* and control before and after treatment with MSC derived exosomes.

Supplemental Figure S14

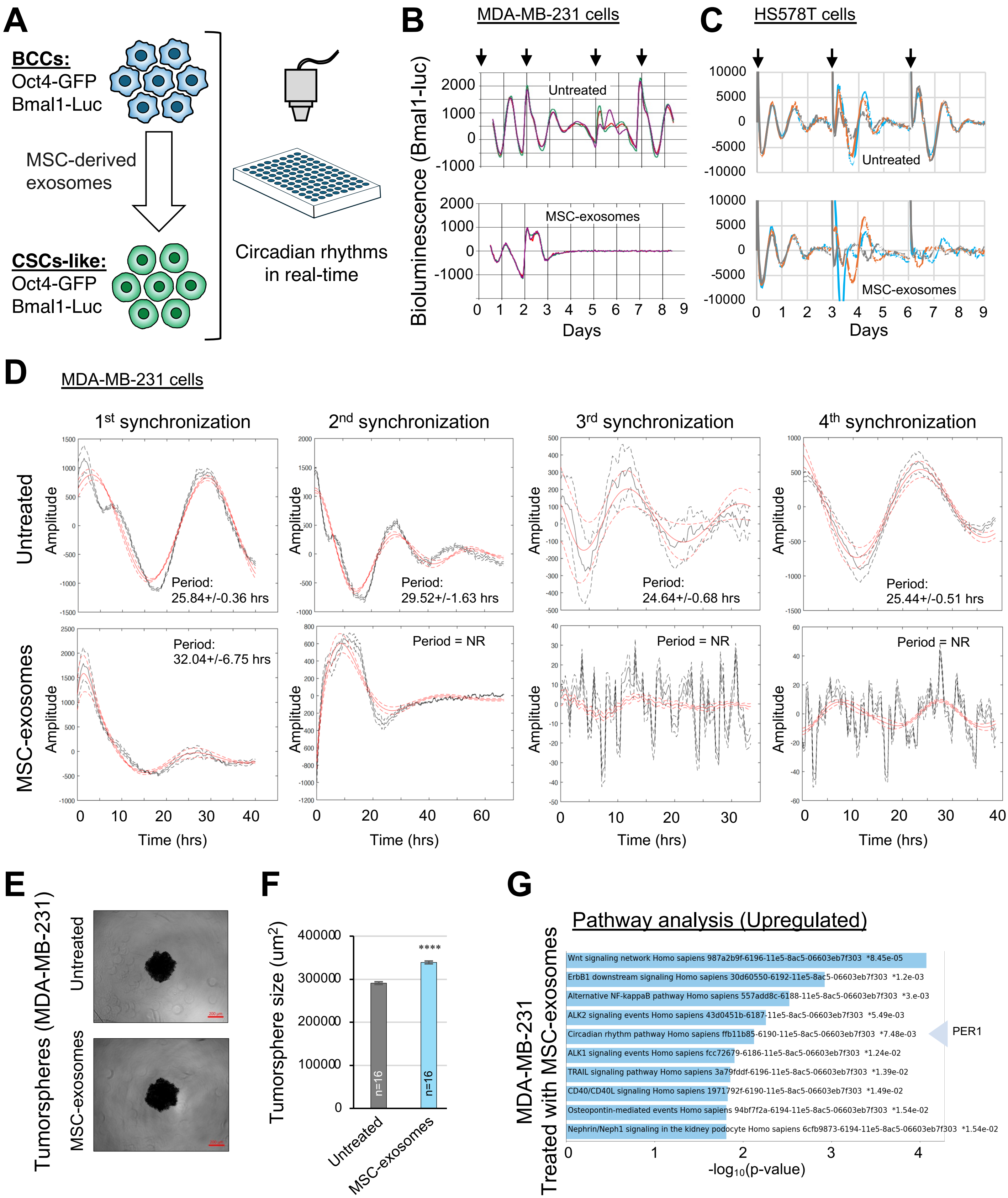

**Supplemental Figure 14. Exosome Effect on Circadian Rhythms and Associated Gene Expression Changes.**
(**A**) Schematic diagram showing the workflow for monitoring circadian rhythms in MDA-MB-231 treated with exosomes derived from mesenchymal stem cells (MSCs) to enrich for CSCs-like cells. MDA-MB-231 expressing *Bmal1-luc* and *Oct4-GFP* reporter are treated with purified exosomes and circadian rhythms are monitored in real-time. (**B**) Bioluminescent circadian profiles of MDA-MB-231 treated with MSC derived exosomes. Arrows indicate resynchronization points. Bioluminescent plots show 3 independent circadian profiles from each treatment. (**C**) Bioluminescent circadian profiles from HS578T MDA-MB-231 treated with $10^8$ MSC derived exosomes. Arrows indicate resynchronization points. Bioluminescent plots show 3 independent circadian profiles from each treatment. (**D**) Evaluation of circadian periodicities in MDA-MB-231 treated with MSC derived exosomes after each synchronization cycle. Circadian periodicity from untreated MDA-MB-231 served as control. (**E**) Representative images of 3D tumorsphere assay using untreated MDA-MB-231 and MDA-MB-231 pretreated with $10^8$ MSC derived exosomes (Scale bar = 200µm). (**F**) Quantification of the surface area for tumorspheres shown in Fig. S14E. Error bars represent SEM, (n=16), pair-end t-test, (**** $p<0.0005$). (**G**) Selected pathway of interest from Reactome Pathway analysis result for MSC derived exosome treated MDA-MB-231.

### Supplemental Figure S15

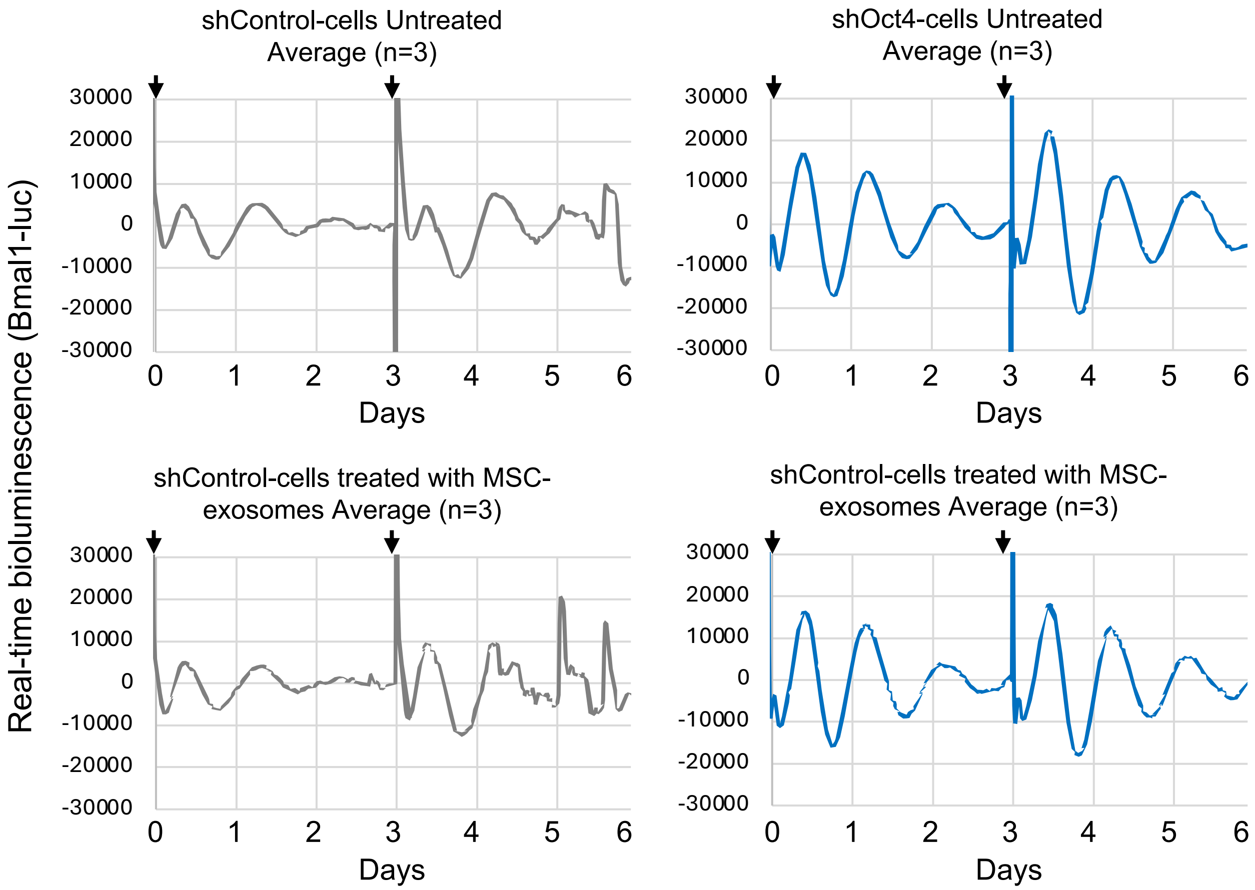

**Supplemental Figure 15. Oct4 deficiency Maintains Circadian Rhythms upon Exposure to MSC-derived Exosomes**

Bioluminescent circadian profiles of MDA-MB-231 deficient in Oct4 or control treated with or without MSC-derived exosomes. Arrows indicate resynchronization points. Bioluminescent plots show an average of 3 independent circadian profiles from each treatment.
